# Cell division protein A (CdpA) organises and anchors the division ring at midcell in haloarchaea

**DOI:** 10.1101/2024.07.17.603854

**Authors:** Yan Liao, Vinaya Devidas Shinde, Dalong Hu, Zhuang Xu, Bill Söderström, Katharine A. Michie, Iain G. Duggin

## Abstract

Many archaea appear to divide through the coordinated activities of two FtsZ homologues (FtsZ1 and FtsZ2) and another bacterial cell division homologue (SepF), which are part of the midcell division ring. Here, we identify an additional protein (HVO_0739, renamed CdpA) that is involved in cell division in *Haloferax volcanii*, with homologues in other Haloarchaea. CdpA localizes at the mid-cell division ring, and this requires the presence of the ring-assembly protein FtsZ1. The division constriction protein FtsZ2 also influenced the proper midcell assembly and structure of CdpA. In the absence of CdpA, cells frequently failed to divide properly, and FtsZ1 formed poorly condensed pseudo-helical structures spanning across a broad region of the cell, whereas FtsZ2 showed mispositioned foci, nano-rings, and filaments. The rate of directional movement of FtsZ1 and FtsZ2 structures around the division ring appeared minimally affected by loss of CdpA, which resulted in continual repositioning of the aberrant FtsZ structures in the cells. In contrast to the FtsZ proteins, CdpA formed relatively immobile foci around the ring. Protein domain function studies, pull-down assays, and multimer structure predictions suggest that CdpA is part of a membrane complex that tethers FtsZ2 and other division proteins to the midcell membrane. Our discovery of an archaeal FtsZ organisation and midcell anchor protein offers new insights into cell division mechanisms that are similar across the tree of life.

## Introduction

Cell division is one of the most fundamental processes in all organisms. It is universally carried out through the localized assembly of cytoskeletal proteins into a ring-like structure linked to the cell membrane and envelope that initiates a constriction process leading to division. In most bacteria, the cytoskeletal protein FtsZ and other cytoskeletal and accessory proteins form the division ring that dynamically drives cell envelope constriction^1^. Bacterial FtsZ appears to perform multiple functions, including acting as scaffolding for protein recruitment to the division site, triggering membrane constriction^2,3^, and dynamically guiding cell wall synthesis to drive its inward growth^4,5^. The relative contributions of these mechanisms to division *in vivo* are yet to be precisely defined.

The eukaryotic homolog of FtsZ is tubulin, which greatly elaborates on the core polymerization function common to this superfamily of proteins, and forms microtubules that provide structural support and play essential roles in diverse cellular processes including intracellular transport and division, although tubulin is not considered a main component of the eukaryotic division ring^6,7^. Multiple tubulin proteins exist in each species, with diversified and specialized functions to support the complex cellular events and differentiation processes that they underpin^8^.

Most bacterial species contain only one FtsZ, but archaea, which are phylogenetically more closely related to eukaryotes, most commonly have two FtsZ homologues belonging to distinct families^9–11^. The dual-FtsZ system might resemble an early stage in the evolutionary diversification and specialization of cytoskeletal functions, as seen in eukaryotes. By using the archaeal model organism, *Haloferax volcanii*, previous work showed that they divide through the coordinated activities of FtsZ1 and FtsZ2^12,13^. The role of archaeal FtsZ1 was primarily ascribed to promoting assembly of the midcell ring, whereas FtsZ2 was more directly implicated in the ring constriction process^13^. Thus, some of the multiple functions of bacterial FtsZ appear to be at least partially separated into specialized homologs in the archaeal dual-FtsZ system. Furthermore, FtsZ2 was found to be specifically absent in certain methanogenic archaea that—rarely in archaea—have a peptidoglycan (PG) cell wall, suggesting that FtsZ2 is needed for division when a PG wall is not present to support the inward growth of the cell envelope^13^.

Current knowledge of FtsZ-based cell division in archaea is limited. There are only a few proteins in this system that have been identified and characterized, despite the many different components present in bacterial and eukaryotic cell division systems. In addition to the two FtsZ proteins, an archaeal homologue of the bacterial division protein SepF^14,15^ and the very recently reported PRC barrel domain proteins (CdpB1/2)^16,17^ were found to have roles in division. *H. volcanii* SepF associates with FtsZ2 at the division site and potentially the cell membrane^14^, and CdpB1 also associates with SepF^16,17^. Crystal structures and AlphaFold2 multimer predictions suggested a tripartite complex formed by CdpB1, CdpB2 and SepF, with CdpB1 and CdpB2 forming a polymer of alternating heterodimers^16^. Currently, there is very little information on what functions these proteins play in the division mechanism or how they are linked to the envelope at midcell.

Other expected components of the FtsZ-based archaeal divisome are yet to be identified, though candidate genes linked to cell division were identified on the basis that their expression or promoter regions were influenced by a transcriptional regulator that appears to control cell division, CdrS, which is encoded by a small ORF in an operon with *ftsZ2*^18,19^. In *H. volcanii*, CdrS-regulated genes included *ftsZ1*, *ftsZ2*, *sepF*, *cdpB1* (Hvo_1691), and the diadenylate cyclase gene (*dacZ*) implicated in signal transduction and regulation of division^19,20^. Another previously uncharacterized gene, HVO_0739, was, with *sepF*, amongst the most strongly regulated by CdrS^19^. Here we have determined the function of HVO_0739, renaming the protein CdpA (<u>C</u>ell <u>d</u>ivision <u>p</u>rotein <u>A</u>), and we propose the Cdp- prefix may be given to other haloarchaeal proteins when they are discovered to participate directly in the division mechanism. We applied genetic and protein structure-function analyses combined with high-resolution fluorescence microscopy to demonstrate the functions of CdpA in organising the assembly of the two FtsZ proteins to anchor them at midcell for division.

## Results

### CdpA is conserved in haloarchaea and required for normal cell division in *H. volcanii*

We identified CdpA (HVO_0739) homologs encoded in 506 published archaeal genomes and performed a phylogenetic analysis of the amino acid sequences. The *cdpA* gene is typically present as a single gene in Haloarchaea, in which it co-exists with the two FtsZs (Fig. 1a) in 96.5% of the species analyzed. Based on protein topology and structure predictions (Fig. 1b), and a multiple sequence alignment (Supplementary Fig. 1), CdpA is comprised of three main regions: a conserved N-terminal transmembrane domain, a less conserved cytoplasmic disordered linker region, and a small cytoplasmic C-terminal domain. CdpA does not have a direct homolog in Bacteria or some of the more distantly related archaeal taxa, although we observed homologous CdpA-like proteins, which showed significant homology but differed in length (± 20%) or number of N-terminal transmembrane segments (≠ 4) compared to the reference CdpA (HVO_0739) in a diverse range of archaea (Fig. 1a). Some Halobacteriota species contained multiple homologs within the genome (Fig. 1a). For example, in *H. volcanii*, a CdpA-like protein (HVO_2016) with four predicted transmembrane segments lies adjacent to HVO_2015, which encodes an actin superfamily protein (volactin)^21^ hinting of a potential common function associated with diverse cytoskeletal systems.

**Fig. 1.**
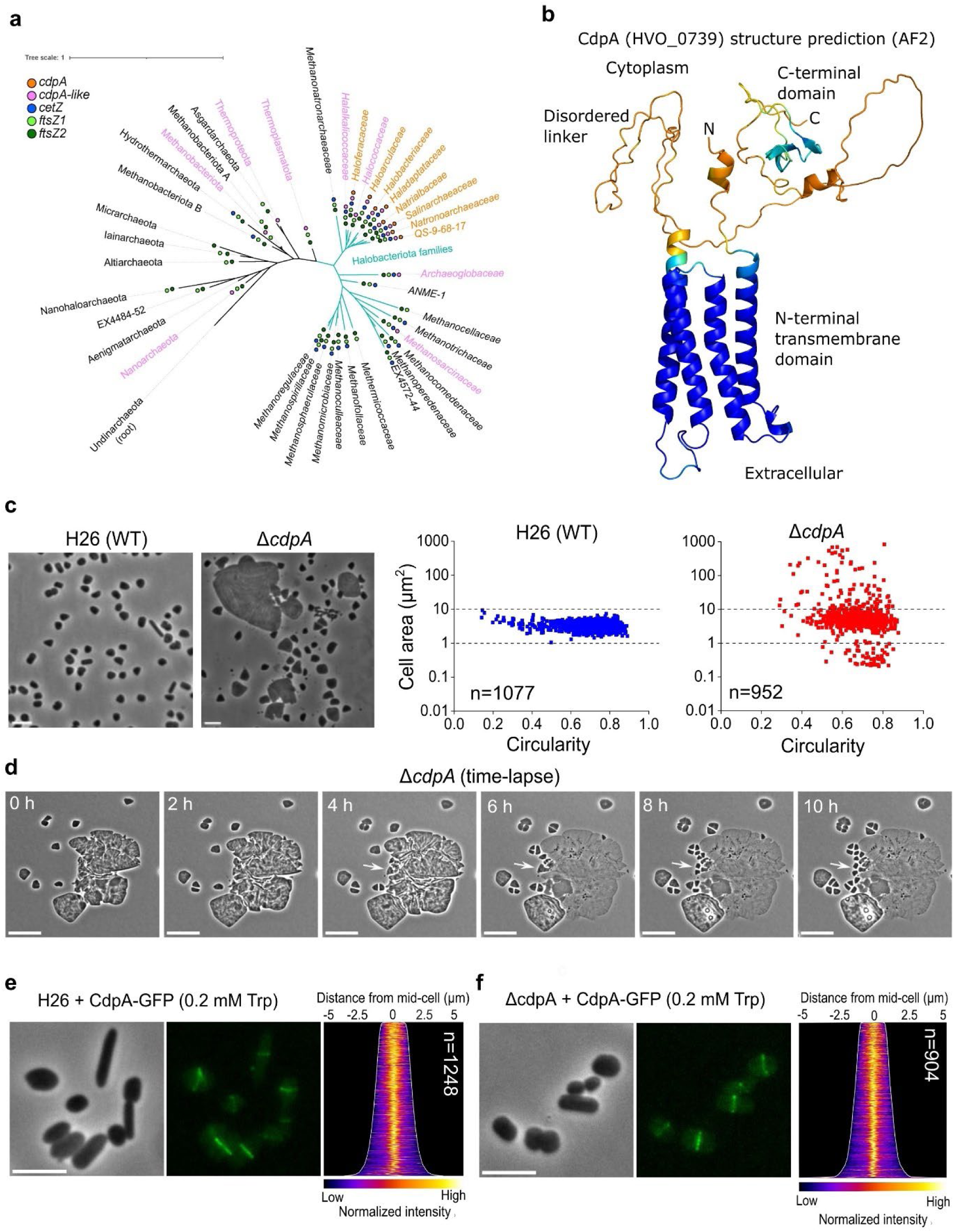
CdpA is cell division protein in haloarchaea. a, Phylogenetic distribution of CdpA (orange circles), CdpA-like (pink), CetZ (blue), FtsZ1 (light green), and FtsZ2 (dark green) on a schematic reference phylogeny of the Archaea (Genome Taxonomy Database, release 207), expanding on Halobacteriota families. The presence /absence is shown at the indicated taxonomic level and is marked as present if identified in at least one genome within that taxon. **b,** Alphafold2 prediction of CdpA monomer structure coloured by model confidence (blue - pLDDT > 90%, orange - pLDDT < 50%) and predicted overall membrane topology. **c**, Phase-contrast microscopy of wild-type (WT) and Δ*cdpA* strains in steady mid-log cultures and the corresponding cell shape analysis. n= number of cells examined over one biologically independent experiment. The data shown is representative of at least two independent experiments. Source data are provided as a Source Data file. **d,** Live-cell time- lapse images of Δ*cdpA* growing on an agarose media pad and showing cell lysis, fragmentation, and division processes (arrows). **e**, **f,** Phase-contrast and fluorescence micrographs of wild-type cells expressing CdpA-GFP (**e**) and Δ*cdpA* expressing CdpA-GFP (**f**) under control of the *p.tnaA* promoter with 0.2 mM Trp induction. Demographic analysis (righthand panels) showing fluorescence intensity along the long axis ordered by cell lengths, top to bottom. n= number of cells examined over one biologically independent experiment. The data shown in panel d, e, f is representative of at least two independent experiments. All scale bars, 5 µm.

To identify the role of CdpA, we investigated the consequences of deleting the *cdpA* gene in *H. volcanii*. Microscopy revealed a cell division defect (Fig. 1c), where the Δ*cdpA* mutant showed a mixture of some greatly enlarged cells that had failed to divide and others with almost normal size. This great heterogeneity suggests an incomplete division defect where failure of cell division in some cells leads to enlarged cells that have a lower probability of division than the smaller cells, which further exaggerates their size defect. Consistent with this, time-lapse microscopy of growing Δ*cdpA* cells showed that some divided at mid-cell and others initially developed division planes but were then unable to complete division (Fig. 1d, Supplementary Video 1). Interestingly, some of the greatly enlarged Δ*cdpA* cells appeared to lyse suddenly but produced cell fragments that were sometimes capable of ongoing growth and division (Fig. 1d, arrows; Supplementary Video 1); in others, blebbing from cellular lobes was seen. These appear to be typical features of *H. volcanii* cell division mutants that are accompanied by ongoing chromosome replication and metabolism to maintain culture viability^13^.

To verify the cell division phenotype attributed to *cdpA*, we reintroduced the *cdpA* gene under the control of the tryptophan-regulated promoter (*p.tna*) on a plasmid in the Δ*cdpA* background. This rescued the cell division defect in the presence of moderate (0.2 mM) or high (2 mM) concentrations of tryptophan (Trp) (Supplementary Fig. 2), confirming that the *cdpA* open reading frame (ORF) is responsible for the observed cell division phenotype. Overexpression of *cdpA* in a wild-type *H. volcanii* background caused no detected cell division or morphological changes (Supplementary Fig. 2a).

### CdpA localizes at the mid-cell division ring

We investigated the subcellular localization of CdpA by expressing GFP-tagged CdpA from a plasmid in wild-type and Δ*cdpA* strain backgrounds. CdpA-GFP fully complemented the cell division defect of the Δ*cdpA* strain in the presence of 0.2 mM Trp induction or greater, showing that the tagged protein is functional in division (Supplementary Fig. 3). Fluorescence microscopy showed that CdpA-GFP in both strains localized as a band at mid-cell in almost all cells (Fig. 1e, Supplementary Fig. 3). With relatively high concentrations of Trp (2 mM), small extracellular fluorescent particles were also seen (Supplementary Fig. 3b), suggesting that excess CdpA-GFP promotes the generation of such particles from cells. Previously, we also detected comparable small extracellular fluorescent particles detached from cells with minimal division defects when expressing mCherry-tagged FtsZ1 GTPase mutant in a wild-type *H. volcanii* strain^13^. However, the mechanisms underlying the formation of these extracellular fluorescent particles remain unclear. One possible explanation is a budding-like process, as previously observed in *ftsZ* knockout strains^13^.

We then investigated the localization of CdpA together with FtsZ1 or FtsZ2 by production of pairs of fusion proteins from a plasmid in the wild-type background. The degree of co-localization observed between CdpA-GFP and FtsZ1-mCherry was strong, with a correlation coefficient of > 0.7 in 79% of cells (Fig. 2a). CdpA-GFP showed a somewhat higher background throughout the rest of the cell than FtsZ1-mCherry, relative to the midcell bands (Fig. 2a, normalized intensity graphs). The co-production of CdpA-GFP and FtsZ1-mCherry did not cause a substantial change in cell shape and size, similar to when FtsZ1-mCherry or FtsZ2-mCherry are moderately produced^13^. The co-production of CdpA-GFP and FtsZ2-mCherry resulted in enlarged or filamentous cells (Fig. 2a), suggestive of a functional interference between these two tagged proteins. Nevertheless, CdpA-GFP and FtsZ2-mCherry still showed strong co-localization, with 84% of cells having a correlation coefficient of > 0.8 (Fig. 2a, righthand panels). In many of these enlarged cells, CdpA-GFP and FtsZ2-mCherry were co-localized around mid-cell, whereas in some cells they were found near the quarter positions of the long axis of the cell (Fig. 2a).

**Fig. 2.**
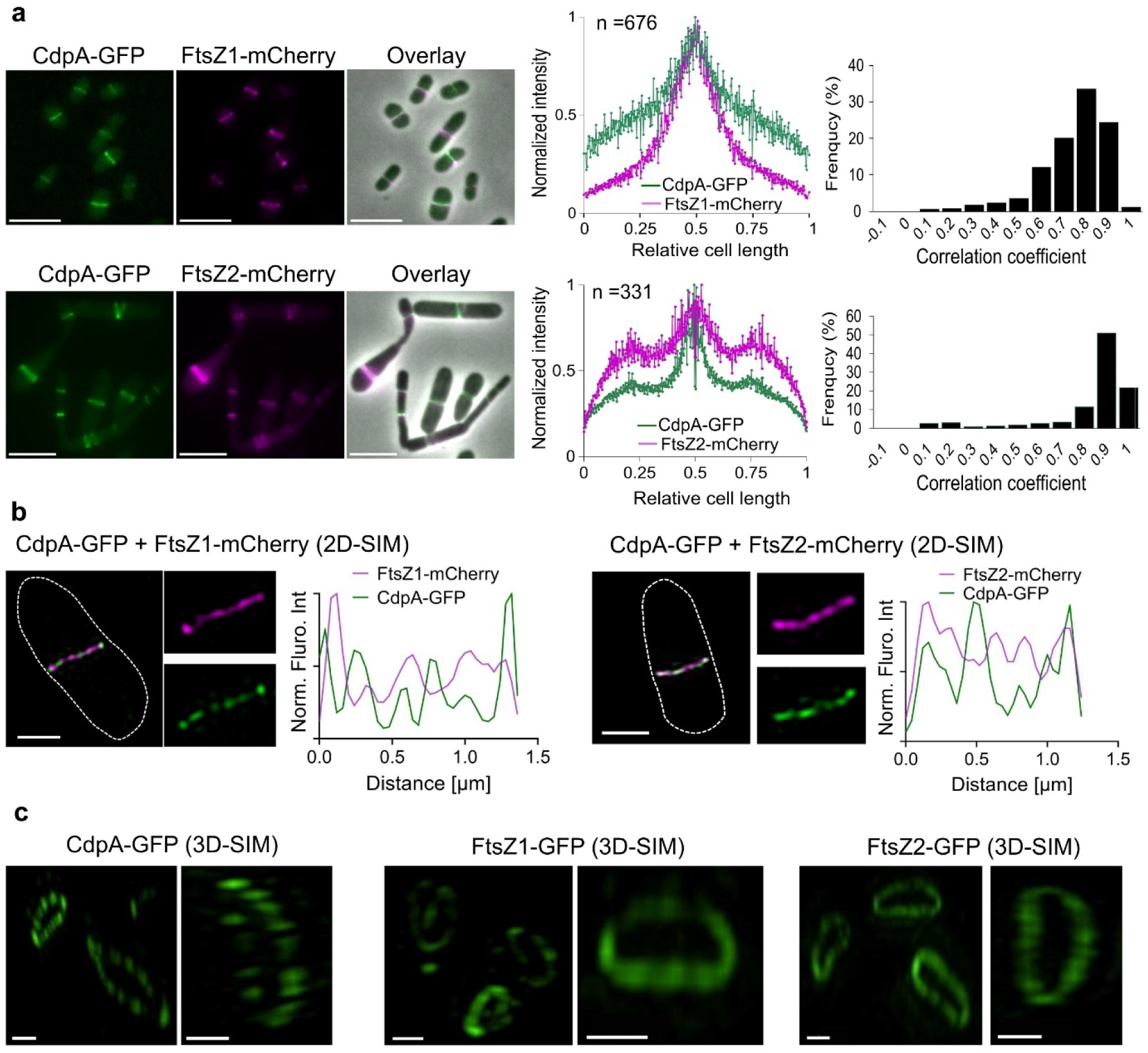
Sub-cellular and sub-ring localization of CdpA with FtsZ1 or FtsZ2. a, Phase-contrast and fluorescence images of *H. volcanii* strains YL6 (H26 + CdpA-GFP + FtsZ1-mCherry dual expression) and YL7 (H26 + CdpA-GFP + FtsZ2-mCherry dual expression) cultured with 0.2 mM Trp during mid-log growth. The middle graphs show the intensity profile of the fluorescence signal (magenta for mCherry, and green for GFP) by plotting the normalized median intensity perpendicular to the long axis of cells. The right graphs show correlation coefficients plotted as a frequency distribution. n = number of cells examined over one independent experiment. The data shown are representative of at least two independent experiments. Scale bars, 5 µm. Source data are provided as a Source Data file. **b,** Dual- colour of 2D-SIM images of mid-log *H. volcanii* YL6 and YL7 grown with 0.2 mM Trp. mCherry (magenta) and GFP (green) fluorescence was quantified by plotting normalized intensity in the respective channel along the ring band. Scale bars, 1 µm. Source data are provided as a Source Data file. **c,** 3D-SIM images (xy-tilted) of mid-log *H. volcanii* YL29 (H26 + CdpA-GFP), YL24 (H26 + FtsZ1-GFP), and YL26 (H26 + FtsZ2-GFP) grown with 0.2 mM Trp. Scale bars, 0.5 µm. The data shown in panel b and c are representative of at least two independent experiments.

We next investigated whether CdpA-FtsZ co-localization depends on the presence of the other FtsZ protein (Supplementary Fig. 4). In the absence of FtsZ1, FtsZ2-mCherry formed patches or foci, with CdpA-GFP co-localizing well with these structures. However, CdpA-GFP also appeared as faint, discontinuous foci along the cell edge or as short patches inside the cells, where FtsZ2-mCherry was not present. In the absence of FtsZ2, CdpA-GFP and FtsZ1-mCherry appeared as broader bands, perpendicular patches or foci, exhibiting substantial co-localization, with some minor regions of weak localization of CdpA-GFP only. The above findings suggest that CdpA has a significant role in organising the assembly of the two FtsZ proteins at the midcell ring.

### CdpA, FtsZ1 and FtsZ2 do not strongly colocalize within the division ring

To investigate protein localization at higher resolution within the division ring, Structured-Illumination Microscopy (SIM) was used to visualize cells co-producing CdpA- and FtsZ- fusion proteins. Live cell 2D-SIM revealed that CdpA-GFP and FtsZ1/2-mCherry showed different patterns of fluorescence within the mid-cell ring (Fig. 2b). Specifically, quantification of fluorescence intensities revealed that CdpA-GFP and FtsZ1- or FtsZ2-mCherry foci were colocalized in some regions but were clearly separated in others (Fig. 2b). Similarly, FtsZ1-mCherry and FtsZ2-GFP exhibited only partial co- localization within the mid-cell ring, as revealed by 2D-SIM (Supplementary Fig. 5).

We then used 3D-SIM, which further resolved the individual ring sub-structures. CdpA-GFP was highly discontinuous within the ring, appearing as distinct small foci with gaps in fluorescence intensity between them (Fig. 2c). FtsZ1-GFP and FtsZ2-GFP rings appeared more continuous but still showed substantial variations in structure and intensity around the ring (Fig. 2c, Supplementary Fig. 6, Supplementary Video 2-4).

### CdpA requires FtsZ1 and FtsZ2 for its proper assembly at midcell

The division ring localization of CdpA, FtsZ1 and FtsZ2, and their different spatial patterns within the ring, prompted us to investigate the dependency of CdpA and FtsZ1/FtsZ2 on each other for proper assembly in the division ring. We assessed the localization of CdpA in the two *ftsZ* knockout strains and vice versa. In the absence of FtsZ1, CdpA-GFP failed to localize as rings and infrequently appeared as a faint midcell focus, and in an *ftsZ1/ftsZ2* double knock-out it showed a diffuse, non-localized signal (Fig. 3a). On the other hand, FtsZ2 deletion did not prevent CdpA-GFP localization although it often appeared as very irregularly structured, poorly condensed or indistinct rings (Fig. 3a). Thus, the proper localization and midcell assembly of CdpA is strongly dependent on the presence of FtsZ1 and FtsZ2.

**Fig. 3.**
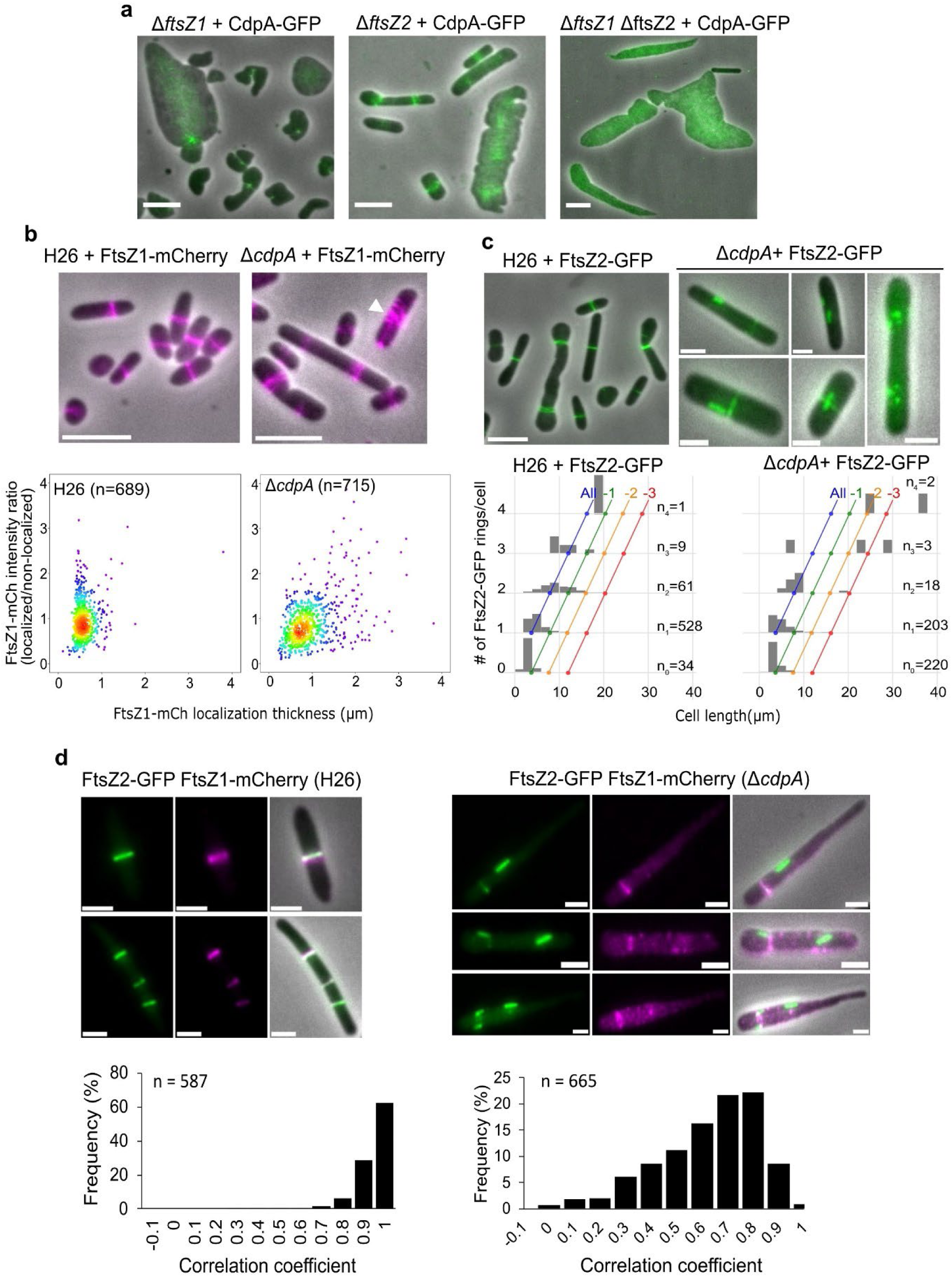
CdpA, FtsZ1 and FtsZ2 show different localization interdependencies and strongly influence each other’s midcell structures. All strains were grown in Hv-Cab medium with SepF, and mid-log cultures were sampled for microscopy. **a,** Phase-contrast and fluorescence composite images of CdpA- GFP in the absence of FtsZ1, FtsZ2, or both. Scale bars, 5 µm. The micrographs shown are representative of at least two independent experiments. **b,** Left: localization of FtsZ1-mCherry in wild-type H26 and Δ*cdpA* backgrounds. The arrowhead indicates a poorly condensed FtsZ1-mCherry structure in Δ*cdpA*. Scale bars, 5 µm. The dot-plots show quantification of the degree of FtsZ1-mCherry localization versus individual localization thickness as a measurement of protein condensation along the long axis in the indicated backgrounds. The data points are coloured with a proximity heatmap. **c,** localization of FtsZ2-GFP in H26 (scale bar, 5 µm) and Δ*cdpA* backgrounds (scale bars, 2 µm). The histograms show the number of FtsZ2-GFP structures detected versus cell length in the indicated backgrounds. Blue diagonals (“All”) represent 1 localization per 4 µm in length (i.e. approximately equivalent to all division sites detected at the normal frequency per cell length in wild-type cells), green (“-1”) indicates one fewer rings/cell length, yellow is two fewer, and red is 3 fewer. n_x_=number of cells for the indicated number(x) of localizations examined over one biologically independent experiment. **d,** Upper: localization of FtsZ1-mCherry and FtsZ2-GFP in H26 and Δ*cdpA* strains. Scale bars, 2 µm, Bottom: correlation coefficient between FtsZ1-mCherry and FtsZ2-GFP plotted as a frequency distribution. The majority (97%) of H26 cells had a correlation coefficient of ≥0.8, while only 32% of Δ*cdpA* cells had a correlation coefficient of ≥0.8. For panel b, c, and d, n= number of cells examined over one biologically independent experiment, and the data shown is representative of at least two independent experiments. Source data are provided as a Source Data file.

### CdpA organizes FtsZ1 and FtsZ2 filaments for proper midcell ring formation

Deletion of *cdpA* had strikingly different effects on FtsZ1 and FtsZ2 function and localization patterns. FtsZ1-mCherry localizations in Δ*cdpA* appeared as rings but with broader zones compared to FtsZ1- mCherry ring structure in wild-type cells and occasional foci seen throughout the cells (Fig. 3b).

Measurements of the ratio of localized to non-localized FtsZ1-mCherry per cell and each individual localization’s thickness (or its width along the medial axis), confirmed that the fluorescence in localized structures was similar in both strains, but the localizations were relatively poorly condensed in many of the *ΔcdpA* cells compared to wild-type (Fig. 3b, graphs). In these strains, the *cdpA* knockout resulted in ∼30% of the total cells that we classified as filamentous (cell area > 5 μm^2^, circularity < 0.3) compared with ∼5% in the wild-type (Supplementary Fig. 7). These findings suggest that CdpA contributes to proper FtsZ1 ring condensation.

Consistent with a previous observation^13^, we saw that production of FtsZ2-GFP caused a division defect in the wild-type background, resulting in somewhat enlarged cells and some showing multiple rings (Fig. 3c). However, in the Δc*dpA* background, FtsZ2-GFP showed fewer rings of normal appearance, and instead frequently assembled as foci, clusters of multiple foci, or short mispositioned filaments located along the cell edge in a perpendicular orientation compared to normal rings (Fig. 3c). Automated detection of the frequency of FtsZ2-GFP rings along the cell length confirmed that there were overall fewer rings in the Δc*dpA* background compared to the wild-type (Fig 3c - compare the number of cells in each group sorted by the number of detected rings). We also observed similar mispositioned or perpendicular structures of FtsZ2-mCherry in the absence of CdpA (Supplementary Fig. 8). These observations suggest that a key role of CdpA is the organisation of FtsZ2 filaments into properly structured and orientated rings.

Highly informative results were obtained when producing both FtsZ1-mCherry and FtsZ2-GFP in the Δ*cdpA* background. The absence of CdpA resulted in obvious uncoupling of FtsZ1-mCherry and FtsZ2- GFP filaments, with a wide range of correlation coefficients (0.3-1) in most cells (Fig. 3d). FtsZ1- mCherry appeared as rings with additional poorly condensed foci throughout the cells, whilst FtsZ2- GFP was seen as foci or mispositioned filaments, as seen during the single expression experiments above (Fig. 3d). Experiments with a lower concentration of tryptophan inducer (0.1 mM Trp), to reduce the concentration of FtsZ1-mCherry and FtsZ2-GFP^13^, still showed similar patterns with broad zones of FtsZ1-mCherry localization and foci or mispositioned FtsZ2-GFP (Supplementary Fig. 9).

We used 3D-SIM to further investigate the unusual FtsZ2-GFP patterns in the absence of CdpA. FtsZ2- GFP bands observed in widefield images were seen as incomplete and poorly organized rings in 3D- SIM (Supplementary Fig. 10-11). Furthermore, some of the FtsZ2-GFP foci in widefield were resolved as nano-rings, -helices, or short filaments in 3D-SIM (Supplementary Fig. 10-11, Supplementary Video 5), which were not associated with division constriction sites. The FtsZ2-GFP nano-rings displayed variation in orientation and size, ranging approximately from 300 to 700 nm (mean ± SEM = 463 ± 18 nm, n = 40). Additionally, the perpendicular filaments of FtsZ2-GFP apparent in widefield images were clearly resolved as mostly straight filaments along the cell edge in 3D-SIM (Supplementary Fig. 10-11).

We conclude that CdpA plays an important role in organization of FtsZ1 and FtsZ2 at midcell by promoting FtsZ1 ring condensation and the correct organization, orientation and assembly of FtsZ2 filaments.

### CdpA is a relatively immobile midcell ring ‘anchor’ that has minimal effects on the dynamics of FtsZ1- GFP and FtsZ2-GFP assemblies

To investigate CdpA and FtsZ movement at midcell, and whether CdpA influences FtsZ dynamics within the division ring, we monitored the fluorescence of CdpA-GFP and FtsZ1/2-GFP in live cells over time. We initially recorded 3D-SIM images with 1 min intervals and saw that FtsZ1-GFP was dynamic within the ring, with changes in the intensity and location of clusters of fluorescence intensity over time (Supplementary Fig. 12, Supplementary Video 6), consistent with previous lower-resolution observations^13^. In contrast, CdpA-GFP showed no such movement in the ring over the same period (Supplementary Fig. 13, Supplementary Video 7).

To better enable cluster tracking and reduce bleaching during image acquisition, we used total internal reflection fluorescence (TIRF) microscopy to record protein movement near the cell surface closest to the coverslip. Visual inspection of kymographs generated from the TIRF data recorded at the division ring using 4 s frame intervals over 15 min, suggested that clusters of FtsZ1-GFP fluorescence were moving in a directional manner (Fig. 4a). Further analysis revealed that these clusters exhibited complex zigzag and crossover patterns, reflecting FtsZ1-GFP cluster movement in both directions around the division plane (Fig. 4a, Supplementary Video 8-9). FtsZ2-GFP showed similar zigzag-like patterns, but they were generally slower than FtsZ1-GFP (imaging required 10 s intervals over 40 min, Fig. 4b, Supplementary Fig. 14, Supplementary Video 10-11). In contrast, visualization of CdpA-GFP with TIRF over the 15- and 40-min periods showed no substantial directional movement and greatly reduced fluctuations of fluorescence intensity over time compared to the FtsZs (Fig. 4c, 4d, Supplementary Fig. 14-15, Supplementary Video 12-15).

**Fig. 4.**
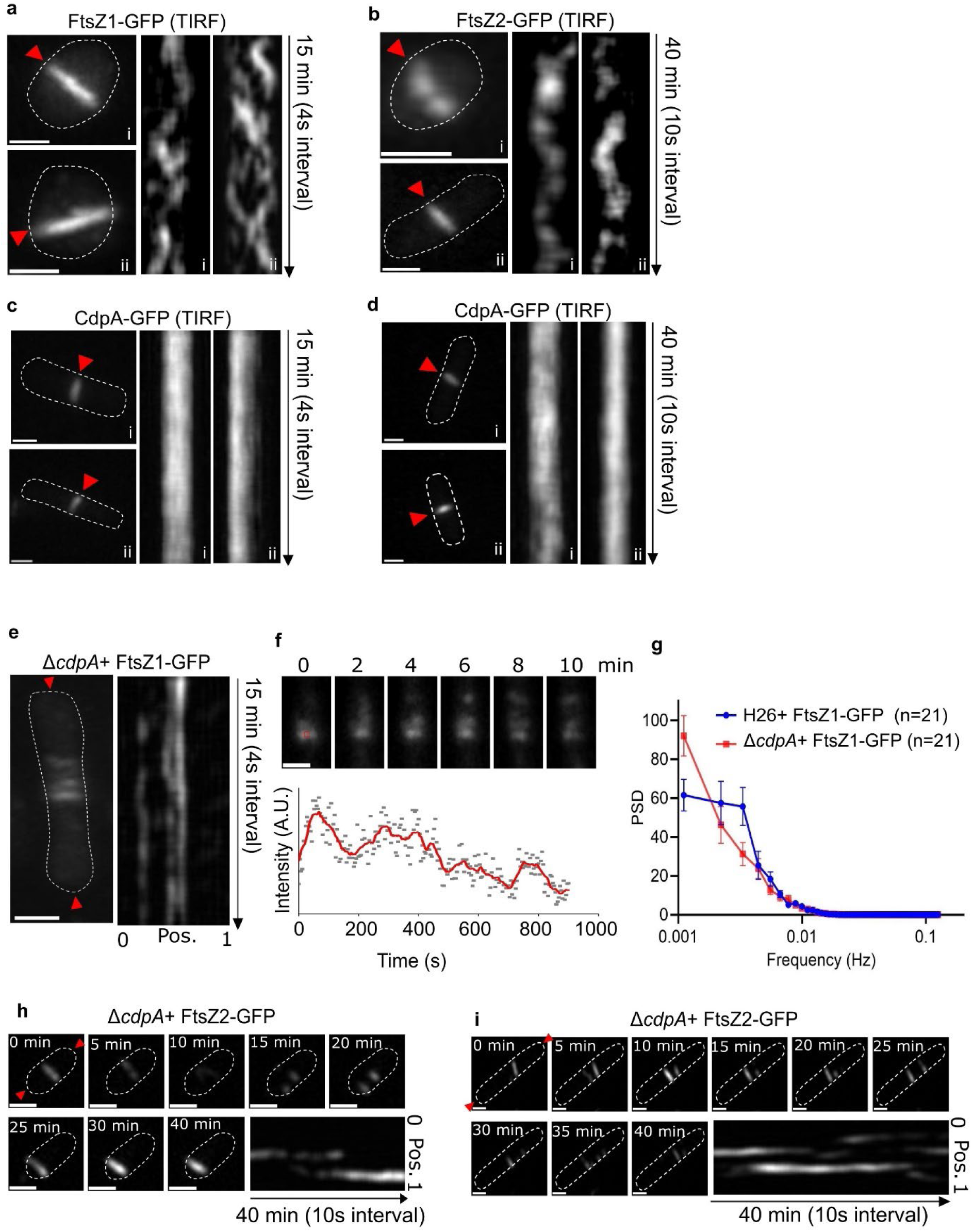
Division protein localization dynamics and the effect of *cdpA* deletion on FtsZ1 and FtsZ2 movement. Cells were grown in Hv-Cab without tryptophan induction to minimize the expression level of fluorescent fusion proteins. Mid-log cultures were sampled for TRIF imaging. The data shown is representative of at least two independent experiments. **a,** Maximum intensity projections (MIPs) of two representative wild-type H26 cells expressing FtsZ1-GFP (i and ii) are shown on the left, with the corresponding kymographs at positions indicated by red arrows in the MIPs shown on the right. **b,** MIPs of two representative wild-type H26 cells expressing FtsZ2-GFP (i and ii) and their corresponding kymographs. **c-d,** MIPs of wild-type H26 cells expressing CdpA-GFP and their corresponding kymographs with different imaging time-window (4 s intervals for 15 min, panel c; 10 s intervals for 40 min, panel d). **e,** MIP of one representative Δ*cdpA* expressing FtsZ1-GFP and its kymograph computed from the intensity along the long axis of cell (between the two red arrows). The relative position (0-1) along the long axis of cell is labelled below the kymograph. **f,** Montages of FtsZ1-GFP dynamics in Δ*cdpA* and the integrated fluorescence time trace for the ROI region, as shown as a small red box around the centre of ring at the start of imaging (Time 0 min). The red curve is the moving average (every 5 points) of the raw intensity (grey dots). Source data are provided as a Source Data file. **g,** The mean power spectral density (PSD) curves (± SEM) of FtsZ1-GFP in wild-type and Δ*cdpA* cells (n=21). Source data are provided as a Source Data file. **h-i,** Montages of representative FtsZ2-GFP dynamics in two Δ*cdpA* cells and their corresponding kymographs indicate the changes in ring position (h) and orientation (i). All scale bars are 1 µm.

It is important to note that the function of FtsZ2-GFP in cell division is more severely affected by the GFP tag than FtsZ1-GFP^13,22^, whereas CdpA-GFP is fully functional (Fig. 1f). The dynamics of FtsZ2-GFP or its response to other proteins might therefore be significantly affected by the tag, compared to the untagged native FtsZ2. Here, our main aim was to determine whether the relative dynamics of each of the existing FtsZ-GFPs is altered by the presence or absence of CdpA. In the absence of CdpA, FtsZ1- GFP still showed clusters with apparently similar movement rates as the wild-type background, and as expected, they often spanned a relatively uncondensed zone around midcell (Fig. 4e, Supplementary Video 16-18). We then measured the FtsZ1-GFP fluorescence intensity over time in a square region near the centre of a localization, and this appeared to show irregular, non-periodic fluctuations (Fig. 4f, Supplementary Fig. 16). To identify any periodicity or dominant frequencies, we plotted the power spectrum density (PSD)^5^ in 21 cells of each strain (Fig. 4g, Supplementary Fig. 17). In the wild-type background, there was an elevated FtsZ1-GFP periodicity at ∼0.003 Hz on average compared to Δ*cdpA* in which the average fluctuations showed a simpler decay-like curve consistent with a lack of periodicity detected within the 15 min. The wild-type signal around 0.003 Hz was dominant in ∼24% of cells, and weak or not evident in ∼76%. This heterogeneity might reflect complex patterns of polymerization within the ring, or cells at different stages of midcell maturation and activity (Supplementary Fig. 17). In the absence of CdpA, FtsZ1 polymerization rates or types might be affected, or the loss of ring structural integrity could also lead to the loss of the localized periodic fluctuations.

With FtsZ2-GFP in Δ*cdpA*, we saw striking dynamic shifts in the cellular position and orientation of FtsZ2-GFP structures over time (Fig. 4h-i, Supplementary Video 19-24), consistent with the strong uncoupling and mispositioning of FtsZ2-GFP seen in the static images (Fig. 3c). However, like FtsZ1- GFP, FtsZ2-GFP clusters clearly displayed movement consistent with dynamic polymerization within the localized structures in Δ*cdpA* that was comparable to the wild-type background (Fig. 4h-i, Supplementary Video 10-11, 19-24). We conclude that CdpA is relatively immobile within the ring but strongly influences the stable organization and positioning of the two FtsZ proteins at midcell whilst having minimal or moderate influence on FtsZ-GFP assembly dynamics.

### The three domains of CdpA make different contributions towards its function

CdpA contains a strongly predicted N-terminal transmembrane domain (amino acids 17-149), a cytoplasmic unstructured linker region (150-284) and a smaller C-terminal domain (285-328) (Fig. 1b, 5a). Together with the results above, this expected topology suggests a model in which CdpA tethers and/or helps assemble and orientate cytoplasmic FtsZ filaments and potentially other components at the midcell membrane. We further investigated this by assessing the contributions of the three main regions to CdpA localization and function.

We initially found that an isolated N-terminal domain (CdpA_NTD_-GFP) localized weakly at mid-cell, a NTD deletion construct (CdpA_ΔNTD_-GFP) failed to localize, and a CTD deletion construct (CdpA_ΔCTD_-GFP) efficiently localized at mid-cell (Fig. 5b). The CdpA_NTD_ is therefore essential for midcell ring localization, but insufficient without the CdpA_Linker_ for the normal degree of localization. It is possible that the linker contributes to interactions promoting CdpA’s assembly into the ring, and/or it alleviates potential steric interference by the GFP when attached directly to the CdpA_NTD_.

**Fig. 5.**
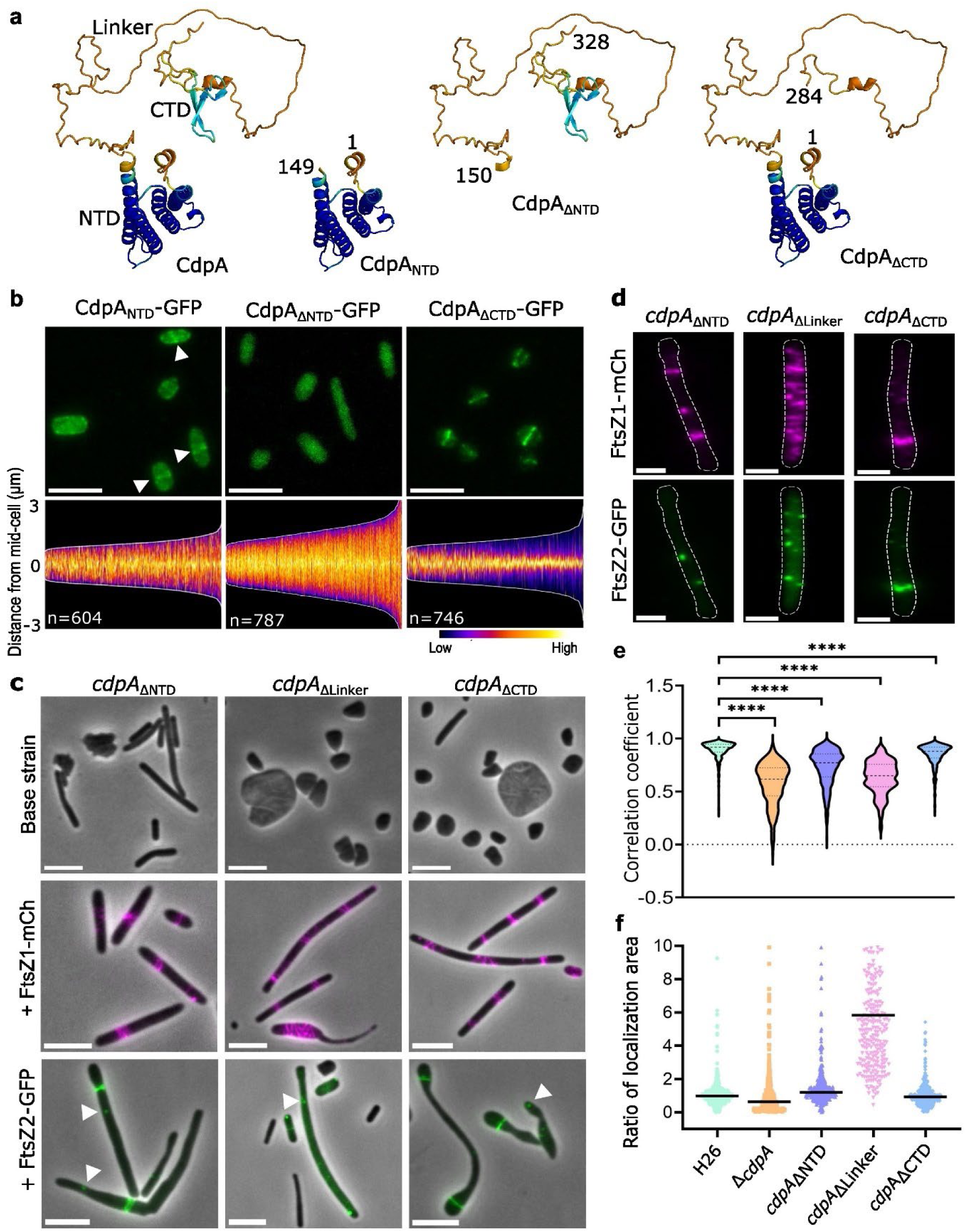
The three domains of CdpA have different contributions to the role of CdpA in of the organisation of FtsZ1 and FtsZ2. a, Alphafold2 models of CdpA showing domain boundaries of the GFP tagged deletion constructs, including the N-terminal domain (NTD), disordered linker, and C- terminal domain (CTD). **b,** Fluorescence microscopy of H26 wild-type strains expressing the indicated constructs during mid-log growth in Hv-Cab medium with 0.2 mM Trp. Arrows indicate regions of weak mid-cell localization. Scale bars, 5 µm. The lower panels show demographic analyses of fluorescence along the long axis, ordered from shortest to longest cells (left to right). n=number of cells examined over one representative experiment. **c,** Phase-contrast and fluorescence micrographs of the indicated *cdpA* deletion strains producing either FtsZ1-mCherry or FtsZ2-GFP, sampled during mid-log phase growth in Hv-Cab + 0.2 mM Trp. Arrows indicate some faint foci of FtsZ2-GFP. Scale bars, 5 µm. **d,** Fluorescence images of the indicated *cdpA* deletion strains co-producing FtsZ1- mCherry and FtsZ2-GFP. Scale bars, 2 µm. **e,** Distributions of cellular correlation coefficients between FtsZ1-mCherry and FtsZ2-GFP and (**f**) ratio of localization area for FtsZ1-mCherry:FtsZ2-GFP in the background strains shown below the lower panel. The sample sizes analysed for each background strain in panel e were as follows: 587 for H26, 665 for Δ*cdpA*, 397 for *cdpA*_ΔNTD_, 424 for *cdpA*_ΔLinker_, and 386 for *cdpA*_ΔCTD._ The statistical test in panel e was performed using an unpaired t-test (two- tailed) in Prism, with **** indicating a significant difference (P<0.0001). The sample sizes analyzed for each background strain in panel f were as follows: 471 for H26, 320 for Δ*cdpA*, 349 for *cdpA*_ΔNTD_, 329 for *cdpA*_ΔLinker_, and 447 for *cdpA*_ΔCTD._ For display purpose (with the maximum y-axis set to 10), 112 data points fell outside the axis limits. The data shown are representative from one of two independent experiments. Source data are provided as a Source Data file.

To investigate the influence of the three CdpA domains on cell division and FtsZ localization, we generated in-frame deletions (scar-less) of each of the three regions in the endogenous chromosomal copy of *cdpA*. In the absence of the NTD (*cdpA*_ΔNTD_), the cells became filamentous (Fig. 5c, Supplementary Fig. 18a); the *cdpA*_ΔLinker_ strain also showed a division defect of enlarged plate cells, and *cdpA*_ΔCTD_ showed a heterogeneous mixture of enlarged plates and filamentous cells. Thus, division defects were clear in all three strains, but they have different cell morphologies. However, these shape differences were neutralized upon transformation with plasmids (containing various FtsZ-FPs), which resulted in a greater frequency of highly filamentous cells (circularity < 0.2) and no enlarged plates (Fig. 5c, Supplementary Fig. 18a). This is consistent with the known influence of *H. volcanii* plasmids on cell shapes^22^.

In all three CdpA domain deletion backgrounds, FtsZ1-mCherry appeared as poorly condensed rings, broad zones of patchy localization or complex helicoid structures, but this was accentuated with *cdpA*_ΔLinker_ (Fig. 5c, Supplementary Fig. 18b), suggesting that this deletion causes additional interference to FtsZ1 localization. Cells producing FtsZ2-GFP in all three domain deletion backgrounds were filamentous, with infrequent rings or localizations usually near one or both poles (Fig. 5c). In the *cdpA*_ΔLinker_ background, rings were very rare and foci were frequently seen near the pole, suggesting this deletion results in additional interference to prevent normal FtsZ2-GFP assembly. In contrast, the *cdpA*_ΔCTD_ background gave the mildest FtsZ2-GFP assembly defects, with rings appearing more frequently than in other mutants, although still less frequently than in the wild-type (Fig. 5c, Supplementary Fig. 18c). Additionally, mispositioned or perpendicular structures of FtsZ2-GFP were also evident in three domain deletion strains (Supplementary Fig. 19).

When we co-produced FtsZ1-mCherry and FtsZ2-GFP in the three domain deletion strains (Fig. 5d, Supplementary Fig. 20), the patterns for each protein were similar but some appeared more accentuated than the corresponding single-FtsZ-FP strains. In the *cdpA*_ΔNTD_ strain, FtsZ1-mCherry localized as rings, poorly condensed rings or foci, while FtsZ2-GFP formed patches, foci, or rare poor rings and they showed a weaker correlation of localization than the wild-type (Fig. 5e). In the *cdpA*_ΔLinker_ strain, FtsZ1-mCherry showed striking irregular bands scattered along the length of the cells, and FtsZ2-GFP showed a combination of diffuse signal with additional scattered foci (Fig. 5d). This strain showed the weakest co-localization amongst the three *cdpA* domain deletions (Fig. 5e) and the FtsZ1- mCherry localizations also occupied a much greater area of the cells than the FtsZ2-GFP localizations (Fig. 5f). In *cdpA*_ΔCTD_, we saw strongly co-localized rings of FtsZ1-mCherry and FtsZ2-GFP, but these still appeared less frequently than in the wild type and were often near one pole (Fig. 5d-f).

The combined results suggest that: 1) The N-terminal transmembrane domain and possibly the linker region of CdpA are important for CdpA midcell localization, FtsZ1 ring condensation, and FtsZ2 ring assembly and positioning; 2) The linker region of CdpA is necessary to prevent strong localization defects caused by direct fusion of the NTD and CTD, possibly by preventing steric hindrance; 3) The C-terminal domain of CdpA does not play a major role in the assembly and coupling of FtsZ1 and FtsZ2 rings, but has at least one other significant yet currently unknown role in cell division.

### CdpA is part of an anchor complex linked to FtsZ2 and other cell division proteins

The SepF division protein *in H. volcanii* has been shown to interact directly or indirectly with FtsZ2 and was suggested to be a membrane anchoring protein for FtsZ2^14^. So we investigated the effects of CdpA absence on SepF localization, and vice versa. In the absence of CdpA, SepF-GFP expressed at moderate levels (0.2 mM Trp) formed sharp midcell bands in the small plate cells, and in giant plate-like cells it appeared as complex clusters that appeared detached from the envelope—similar to the patterns observed for FtsZ1/2 in giant plate cells (Supplementary Fig. 21)^13^. However, higher expression of SepF-GFP (2 mM Trp) resulted in the formation of striking branched and less condensed mid-cell bands in the small plate cells (Fig. 6). Conversely, during SepF depletion (as SepF could not be deleted^14^), CdpA-GFP formed very few rings, which were consistently positioned approximately one wild-type cell length from the poles of the highly filamentous cells (Fig. 6b). Plotting the number of CdpA-GFP rings over the cell length in the cell population confirmed that ring frequency along the cells was much lower than in the wild-type background (Fig. 6b). These findings suggest that correct CdpA assembly depends on SepF, while SepF localization is, in turn, influenced by the presence of CdpA.

**Fig. 6.**
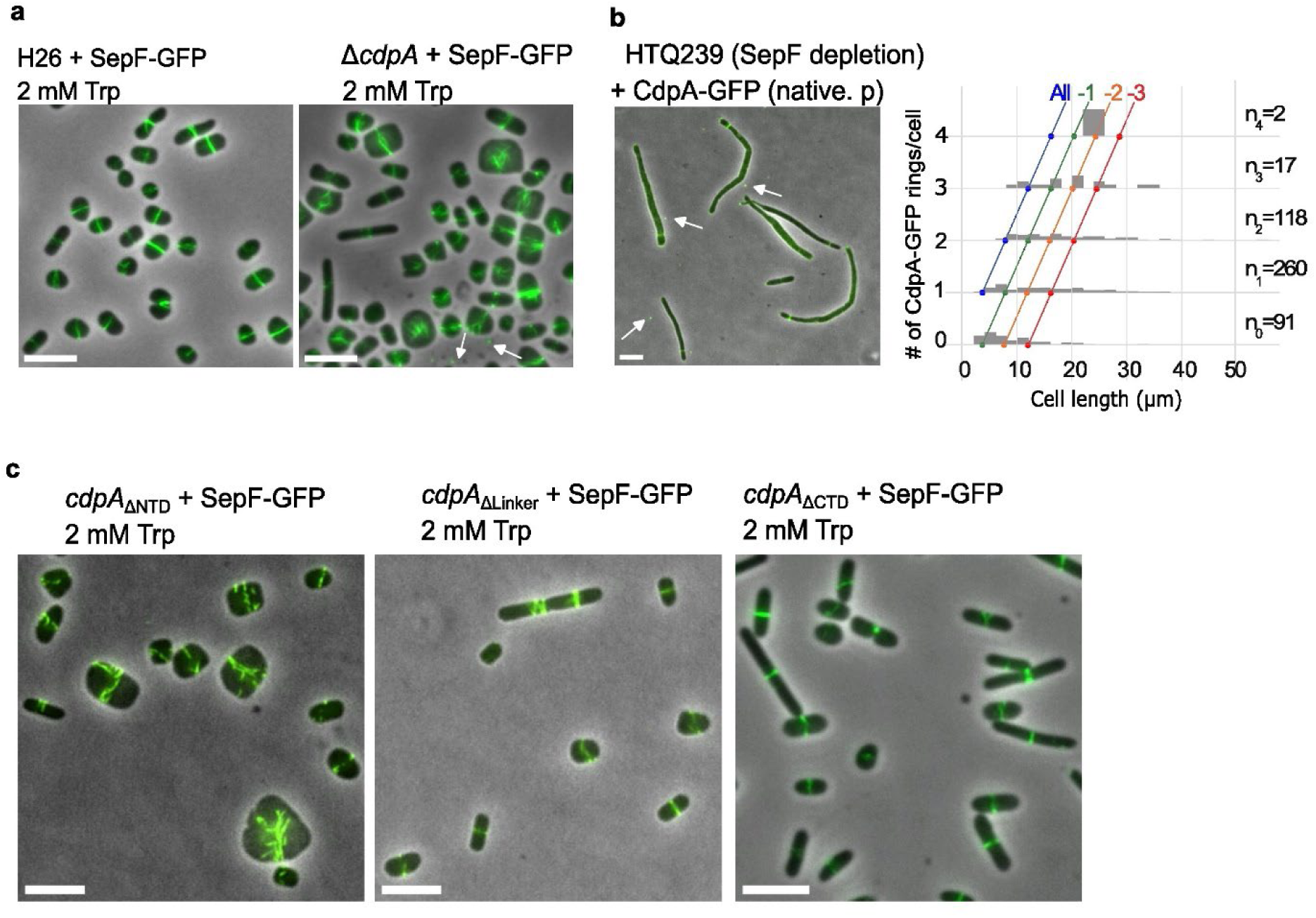
The interdependency of CdpA and SepF localization. a, Phase-contrast and fluorescence micrographs of wild-type H26 and *cdpA* deletion strains producing SepF-GFP, sampled during mid-log phase growth in Hv-Cab + 2 mM Trp. Microscopic images of the strains grown under 0.2 mM Trp can be found in Supplementary Fig. 21. At higher expression levels, SepF-GFP formed branched and less condensed mid-cell bands in the wild-type-like cells in the absence of CdpA. **b,** Phase-contrast and fluorescence micrographs of SepF deletion strain (HTQ239) expressing CdpA-GFP under its native promoter. The histograms show the number (#) of CdpA-GFP rings detected versus cell length. Coloured lines indicate a nominal WT frequency of rings (All), or how many fewer rings were observed, as per Fig. 3c. n= number of cells examined over one biologically independent experiment with indicated ring number, and the data shown is representative of at least two independent experiments. Source data are provided as a Source Data file. **c,** Phase-contrast and fluorescence micrographs of three CdpA domain deletion strains producing SepF-GFP, sampled during mid-log phase growth in Hv-Cab + 2 mM Trp. White arrows indicate small extracellular fluorescent particles. All scale bars, 5 µm. The micrographs shown are representative of at least two independent experiments.

Given the pronounced abnormalities in SepF-GFP localization at higher expression levels in the absence of CdpA, we further examined how the three domains of CdpA influence SepF-GFP localization under these conditions. In the three CdpA domain mutants, SepF expression failed to fully complement the cell division defects, leading to the formation of giant, filamentous cells (Supplementary Fig. 21). In *cdpA*_ΔNTD_, SepF-GFP structure at midcell significantly disrupted, forming branched bands, whereas in *cdpA*_ΔCTD_ and *cdpA*_Δlinker,_ SepF-GFP localizations remained largely unaffected (Fig. 6). These findings suggest that the N-terminal domain (NTD) of CdpA plays a more direct role in mediating SepF assembly at midcell.

Given the above results, we searched for CdpA interactions with other *H. volcanii* proteins by isolating affinity-tagged CdpA (CdpA-HA) from cross-linked *H. volcanii* whole-cell lysates. Proteins remaining bound to the anti-HA magnetic beads after washing with a mild-detergent buffer to reduce non-specific associations were eluted and analysed by western blots probed with antibodies recognising HA, FtsZ1 and FtsZ2 (Fig. 7a). Both FtsZ1 and FtsZ2 were detected in the lysate (L) and unbound (FT) samples, but only FtsZ2 was clearly detected in the bound protein elution (E) sample (Fig. 7a). Neither protein was detected in the elution from the control cells containing the HA vector only (Fig. 7a). These results suggest that CdpA can exist in a complex with FtsZ2.

**Fig. 7.**
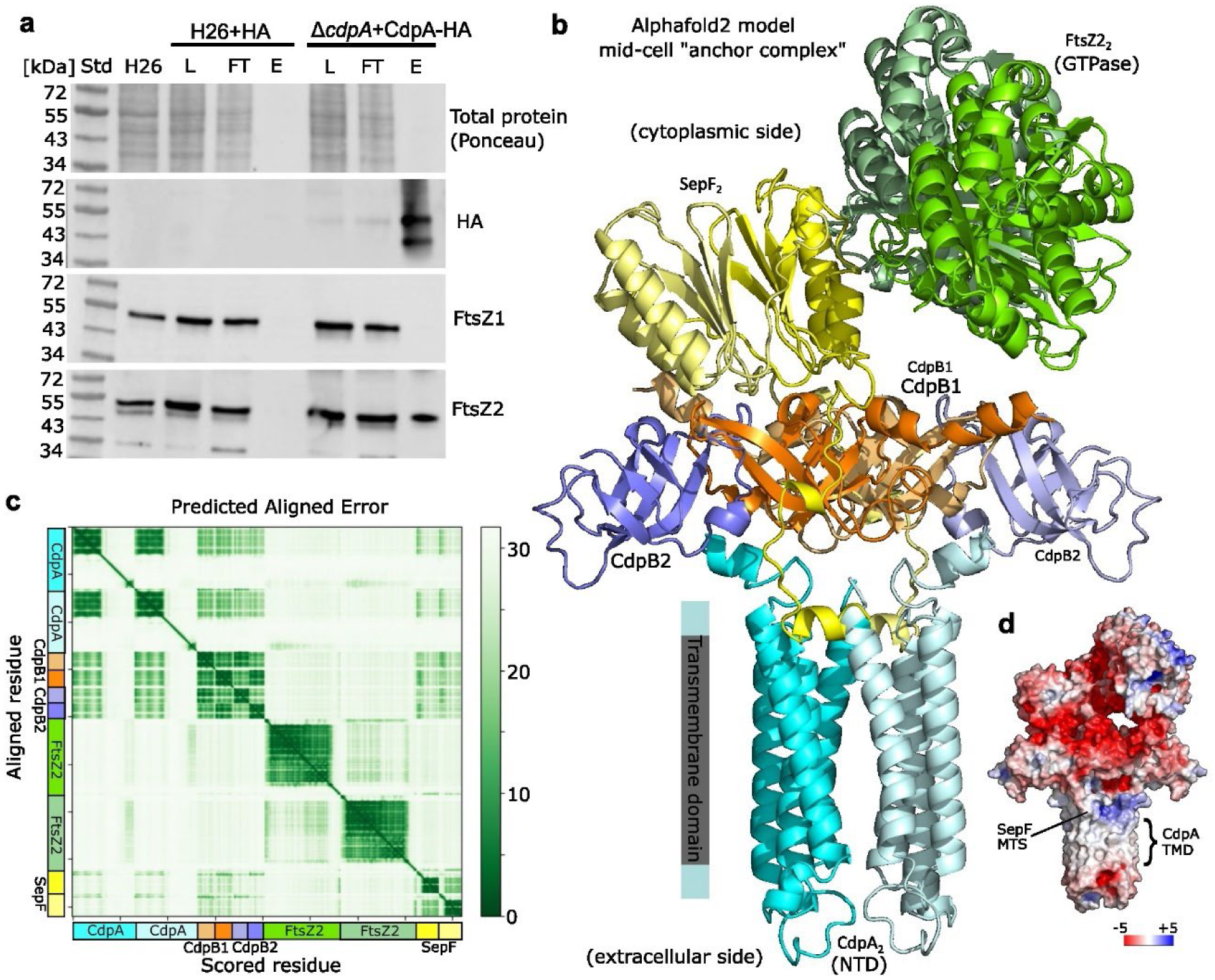
CdpA is associated with FtsZ2 and other division proteins in a putative membrane anchor complex. a, Western blots of one of three replicate co-immunoprecipitation experiments. The cell lysate (L), unbound protein flow through (FT) and bound protein elution fraction (E) from control (H26 + HA vector only) and Δ*cdpA* + CdpA-HA were analysed by 15% SDS-PAGE and western blotting. The total protein stain (Ponceau S) is shown for the same membrane used for the anti-HA probe, and replicate membranes were probed with anti-FtsZ1 or anti-FtsZ2 as indicated. The control (H26 + HA vector only) was not detectable within the size limits of the gel. **b,** Alphafold2 model of the mid-cell anchor complex generated from a sequence input of 2 CdpA, 2 CdpB1, 2 CdpB2, 2 FtsZ2, and 2 SepF. The C-terminal regions of CdpA and the N- and C-terminal extensions of FtsZ2 that are largely disordered are not shown for clarity. **c,** The Alphafold2 predicted aligned error (%) heatmap plot of all input sequences (concatenated), indicating the degree of confidence in pairwise interactions between residues. **d,** The same structure and orientation as in panel b shown with surface electrostatics, indicating the approximate CdpA transmembrane domain (TMD) and SepF membrane targeting sequence (MTS). Unprocessed western blots and the Alphafold2 coordinates are provided in the Source Data file.

We carried out additional pull-down experiments with CdpA-HA and FtsZ2-HA in their corresponding knock-out backgrounds, compared to the wildtype background with HA vector only as the control, and identified bound proteins by mass spectrometry (MS) (Supplementary Table 3). FtsZ1 was detected in the bound-protein elution of both the CdpA-HA and FtsZ2-HA pull-downs, albeit with only moderate statistical support, as assessed by mean MS peak area quantification (Log_2_ fold-change (FC)=2.7, p=0.10 for the CdpA-HA pull-down, and Log_2_FC=3.0, p=0.03 for the FtsZ2-HA pull-down, n=6 independent experiments each). FtsZ2 was identified significantly in the CdpA-HA pulldown (Log_2_FC=5.7, p=0.002), and as expected in the FtsZ2-HA pull-down (Log_2_FC=10.6, p=0.08). The FtsZ2- HA pull-down only identified one other protein with statistical support (i.e., CetZ2, which was not detected in the HA controls; Supplementary Table 3). The low detection rate in the FtsZ2-HA pull- downs might be due to the relatively low production of the FtsZ2-HA fusion (Supplementary Fig. 22), interference by the HA tag (as seen with the function of FtsZ2-GFP) or possibly limited stability of FtsZ2-HA protein complexes in this assay. However, the CdpA-HA construct was produced and complemented the division defect of the Δ*cdpA* background used in the pull-downs (Supplementary Fig. 22). The CdpA-HA pull-down MS data included other proteins previously implicated in cell division, including SepF (HVO_0392, Log_2_FC=4.3, p=0.01), CdpB1 (HVO_1691, Log_2_FC=3.4, p=0.07) and CdpB2 (HVO_1964, Log_2_FC=3.1, p=0.02) (Supplementary Table 3). Other identified proteins recently linked to cell envelope-related functions included actin-like protein volactin (HVO_2015, Log_2_FC=2.5, p=0.06) and small GTPase ArvA (OapA, HVO_3014, Log_2_FC=2.8, p=0.04)^21,23^. It is currently unknown whether these represent direct or indirect associations with CdpA.

To predict how the cell division proteins interact with CdpA, we used Alphafold2 (AF2) Multimer with two molecules each of CdpA, CdpB1, CdpB2, FtsZ2, and SepF, as preliminary AF2 runs and existing homolog structures^16,17^ had indicated that each protein can dimerise or polymerise. This revealed a strikingly coherent predicted structure, consistent with the current experimental results, that we refer to as the anchor complex (Fig. 7b). Most of the visible interactions within the predicted anchor complex are supported by low pairwise alignment error scores between the corresponding regions (Fig. 7c). The main exception to these relatively strong predictions was that the association between the anchor complex and FtsZ2 was quite weak, hinting that interactions between the anchor complex and FtsZ2 might occur via other unknown adaptors. The strongly predicted interactions notably include a direct association of the cytoplasmic short N-terminal helix of CdpA (residues 1-10) with the interface region of each CdpB1/2 heterodimer, which would bring CdpB1/2 close to the inner surface of the membrane (Fig. 7b, 7c). Furthermore, CdpA is predicted to participate in a ternary association with CdpB1 and SepF’s N-terminal extension and membrane targeting sequence (MTS) that places the SepF MTS in an ideal position to penetrate the inner leaflet of the cytoplasmic membrane whilst also bound to CdpA at the interface of cytoplasm and membrane (Fig. 7b, 7c).

These experimental results and structural analysis have indicated that the CdpA NTD acts as a primary membrane anchor located at discrete positions around the division ring, where SepF is expected to play an adaptor-like role (braced by CdpB1/2 multimers) to connect the anchor complex with cytoplasmic ring components including FtsZ2. We found that CdpA has multiple functions, as the inter- domain disordered linker was critical for the proper condensation and positioning of FtsZ1 and FtsZ2 filaments, respectively, whereas the CTD was also important for division in other ways, likely through unknown interaction partners.

## Discussion

In bacteria and eukaryotes, many proteins have been identified to contribute to division ring assembly and anchoring. Here we identify and characterize the function of a ring assembly and anchor protein in haloarchaea, which we named Cell division protein A (CdpA). We found clear roles for CdpA in orchestrating the midcell assembly and co-positioning of FtsZ1 and FtsZ2 filaments to form the division ring in the model archaeon *H. volcanii*. Several other proteins involved in archaeal dual-FtsZ division have recently been identified, including SepF, CdpB1, and CdpB2^14,16,17^, though their functions in the haloarchaeal division mechanism are unclear. Based on the current findings, we developed a structural model in which CdpA acts as the membrane-embedded component of a midcell anchor complex (including SepF, CdpB1, and CdpB2 linked to the membrane though multiple direct interactions with CdpA; Fig. 7). CdpA also functions as a tether and organizing center at relatively immobile and distinct locations around the division ring to promote FtsZ1 and FtsZ2 ring formation and maintenance, and likely other midcell ring proteins, via its cytoplasmic linker and C-terminal domains.

In bacteria, one of the best characterized FtsZ membrane linkage proteins is the widespread FtsA actin-superfamily protein. The essentiality of FtsA varies among different species. For example, in *Escherichia coli*, FtsA is essential^24^, whereas in *Bacillus subtilis*, the *ftsA* gene can be deleted, although this affects Z-ring formation and causes cells to become enlongated^25^. Here, we were able to delete the *cdpA* gene in *H. volcanii*, which caused a division defect resulting in mixed populations of near-normal and greatly enlarged cells. We directly observed some of the greatly enlarged cells forming cellular fragments upon lysis of the main cell body, some of which were viable and capable of ongoing growth and division. This likely explains the survival of many *H. volcanii* cell division mutant strains in culture. Additionally, the *cdpA* deletion strain might survive through the activity of other division site stabilizing or anchoring proteins like SepF^14,26^ or CdpB1/2^16,17^ that potentially compensate for the loss of CdpA. Other currently uncharacterized haloarchaeal proteins, potentially including CdpA-like proteins, may also be involved in midcell ring assembly or cytoskeletal protein anchoring, helping compensate for the loss of CdpA.

Super-resolution imaging revealed that CdpA forms a more discontinuous or patchy ring-like structure at mid-cell compared to FtsZ1 or FtsZ2 rings and differing organization within the ring. In absence of CdpA, FtsZ1 rarely formed normal Z rings and instead formed broad and ill-defined—often helicoid— structures, that still retained dynamic localization patterns. Although typical-looking FtsZ2-GFP rings were sometimes observed in the absence of CdpA, FtsZ2-GFP foci or filaments frequently appeared strongly mispositioned, including perpendicular to the normal ring along the cell edge. FtsZ2 also retained its dynamic properties, resulting in frequent repositioning of the misplaced FtsZ2 foci and filaments. This resulted in FtsZ1 and FtsZ2 becoming uncoupled in their subcellular localization patterns, supporting the notion that they can polymerize independently of one another^13^.

Our protein pull-down experiments detected CdpA in complex with FtsZ2 but not clearly with FtsZ1, despite our finding that the proper localization of CdpA depends on the presence of both FtsZ2 and especially FtsZ1 (and vice versa). Current observations would suggest that FtsZ1 associates only indirectly with the CdpA-anchor complex, via another anchor, or in a manner that is not well maintained during the pull-down assay procedure. The CdpA-HA pull-down assays also identified SepF and CdpB1/2 amongst the interaction partners. Consistent with this, we found that SepF contributes to CdpA assembly at midcell, and thatSepF’s proper localization is strongly affected by the absence of CdpA. Furthermore, the co-localization of CdpA with each FtsZ in strains lacking the other FtsZ suggests a functional interdependence between CdpA and the two FtsZ proteins. Together with the observations of similar poorly-condensed Z-ring structures in the absence of CdpA, SepF^14^, or CdpB1/2^16,17^, the combined results support a model in which these proteins assemble into a complex that functions in Z-ring assembly.

Previous studies reported that *H. volcanii* SepF is part of a complex with FtsZ2^14^, and with CdpB1/2^16,17^. Furthermore, SepF appears to localize at mid-cell via a short N-terminal amphipathic sequence, which was expected to act as a membrane targeting sequence (MTS)^14^. The finding that the SepF MTS alone was sufficient for midcell ring localization^14^ suggests that this short sequence does more than just potential membrane targeting. Additionally, our data indicate that N-terminal domain of CdpA plays a more direct role in mediating interactions with SepF (Fig. 6). Our combined results and structural modelling (Fig. 7) support the existence of a midcell anchor complex that forms through multiple intimate associations, including the N-terminal extensions of CdpA and SepF interacting in different ways with CdpB1/2 heterodimers, and SepF’s MTS (at the very N-terminus) simultaneously associating with the CdpA NTD and partly embedded in the cytoplasmic membrane. The SepF MTS may thereby be sufficient for midcell localization^14^ due to its likely association with CdpA. The main membrane anchoring component appears to be the CdpA NTD, linked to and braced by multimers of SepF and CdpB1/2.

Consistent with the structural model (Fig. 7), our domain localization studies indicated that the CdpA NTD is essential for its midcell localization, whereas the CTD was dispensable for midcell localization but required for normal division. The NTD and linker regions appear to be critical for assembly and anchoring of FtsZ1 and FtsZ2 rings, whereas the CTD seems to not strongly affect FtsZ1 and FtsZ2. These observations suggest a multi-functional role of CdpA in *H. volcanii* division, where the long, disordered linker and C-terminal domain also play important but currently undefined roles in tethering or assembling other proteins at midcell.

CdpA shows some noteworthy similarities and differences to other division ring anchor proteins. In bacteria, the FtsZ ring is typically anchored to the midcell membrane by FtsA, ZipA or SepF, whereas in non-FtsZ archaeal division, based on the ESCRT-III (CdvB) cytoskeletal proteins, CdvA is considered to play a different anchor-like role. FtsA is broadly conserved in most bacteria and contains a short C-terminal MTS and also directly interacts with FtsZ, recruiting it to the inner membrane^27,28^. ZipA is an integral inner membrane protein found in Gram-negative bacteria, and is comprised of a large C-terminal globular cytoplasmic domain linked to a single N-terminal transmembrane domain via an extended linker^29,30^. FtsA and ZipA have overlapping functions but likely serve differing roles in modulating division ring assembly^31,32^. Both FtsA and ZipA interact directly with FtsZ through an interaction hub located at its C-terminal tail^33^. SepF is conserved in Gram-positive bacteria and contains an N-terminal MTS and forms functional dimers, which interact with the C-terminal domain of FtsZ^34,35^. In the Crenarchaeota, *Sulfolobus* CdvA links ESCRT-III (CdvB) proteins to the cell membrane via direct interactions and is thought to assemble on the membrane surface^36^.

Compared to these proteins, CdpA exhibits highest topological similarity to ZipA, suggesting an analogous midcell anchor role. However, ZipA-GFP in *E. coli* showed clear movement around the division ring^37^, yet we found that CdpA-GFP remains relatively stable at mid-cell in *H. volcanii*, suggesting they may operate via different mechanisms within the ring. SepF homologues from different bacterial species can polymerize into large protein rings associated with cell wall remodelling^38^, whereas there is no evidence to show that archaeal SepF is capable of polymerization^14,15^. Our discovery of CdpA as an archaeal division ring organisation and midcell anchoring protein indicates that FtsZ assembly and its attachment to the membrane via a transmembrane anchor are common and significant features shared by both the archaeal and bacterial domains of life.

## Methods

### Identification, phylogeny, and structural analysis of CdpA and CdpA-like sequences

The reference amino acid sequence of HVO_0739 (CdpA) was first input into JackHMMER^39^ to search against UniProtKB database^40^ with three iterations restricted by taxonomy in Archaea (taxid:2157). The output HMM profile was used to scan the amino-acid sequences of all proteins in 1,587 high quality diverse archaeal genomes from NCBI RefSeq database^41^ with a significance threshold of an E-value of 10^-5^. The identified sequences were then confirmed using a BLASTP v2.13.0^42^ search against the reference CdpA sequence with an E-value cut-off of 10^-5^; sequences giving no hit in the BLASTP search result were removed. The remaining 1,336 sequences were taken as CdpA candidates. Of these, proteins with a length between 80% to 120% of HVO_0739 (i.e., 260-400 residues) and with four transmembrane segments (predicted using TMHMM v2.0)^43^ within the N-terminal 75% of the sequences were taken as CdpA family proteins (470 sequences), and the remaining 866 were designated CdpA-like proteins.

Multiple alignment of the 470 CdpA sequences was then performed using MUSCLE v3.8.31^44^. A neighbour-joining tree based on the amino-acid alignment was constructed using PHYLIP v3.698^45^ with the default model and 1,000 bootstrap replicates. HVO_2016 (*H. volcanii* CdpA-like protein) was included as the outgroup to root the tree. The alignment was then reordered by tree structure. A reduced version of the alignment was generated by removing columns that had a gap in the *H. volcanii* CdpA reference sequence. This alignment was displayed using UGENE v46.0^46^. Taxonomy annotation was based on the NCBI Taxonomy database. PsiPred v4.02^47^ was used to estimate the secondary structure of CdpA along the alignment.

Protein 3D structure predictions were downloaded in monomer form from the Alphafold structure database (https://www.alphafold.ebi.ac.uk/entry/D4GTB2, accessed December 2023) or generated as complexes using Alphafold v2.3.1 (Multimer)^48^ using default settings with the full BFD reference database. Structures were visualised and rendered with PyMOL^49^ v2.5.1. Transmembrane topology was predicted with Phobius^50^.

### Growth of *H. volcanii* strains

The *H. volcanii* cultures were grown in Hv-Cab medium^12,51^. Where necessary as auxotrophic requirements, media were supplemented with uracil (50 µg/mL) for Δ*pyrE2* strains. To control gene expression via the *p.tnaA* promoter, 0.2 mM or otherwise indicated concentration of L-tryptophan (Sigma–Aldrich) was supplemented in these cultures. For microscopic imaging, the cultures were incubated at 42°C with rotary shaking (200 rpm) and were maintained in log growth (OD600 < 0.8) for at least 2 days before sampling for analysis of mid-log cultures. Representative data displayed in this paper are representative of at least two independent biological replicate experiments.

### Construction of plasmids for gene deletion and genomic modification in *H. volcanii*

The plasmids, oligonucleotide sequences, and strains are given in the Supplementary Tables 1 and 2. The plasmid pTA131 was used as the basis to construct the plasmids for gene deletion^52^. To make the plasmid for *cdpA* deletion, the upstream and downstream flanks for homologous recombination on either side of *cdpA* were PCR amplified from *H. volcanii* H26 genomic DNA, using the primers given in Supplementary Table 1. The upstream flank fragment and downstream fragments were digested with HindIII and BamHI, respectively, and cloned into pTA131 (at HindIII-BamHI) using Gibson assembly reagents (NEB), resulting in pTA131_up_down_0739. The plasmids pTA131_CdpA_delNTD for N- terminal domain (17-149 aa) deletion and pTA131_CdpA_delLinker for linker region deletion (150-284 aa) were obtained synthesised and cloned (Genscript). To make the plasmid pTA131_CdpA_delCTD for C-terminal domain (285-328 aa) deletion, the upstream flank and downstream flanking fragments of the CdpA C-terminal domain were amplified by using the primers in Supplementary Table 1 and cloned into pTA131 (at HindIII-BamHI) using Gibson assembly (NEB) as described above.

Demethylated plasmids pTA131_up_down_0739, pTA131_CdpA_delNTD, pTA131_CdpA_delLinker, and pTA131_CdpA_delCTD were used to transform *H. volcanni* H26 separately by using the pop-in-pop-out method as described previously to delete the corresponding gene fragments^52^. Allele- specific PCR was used to verify the expected deletion chromosomal structure, resulting in H26-base strains carrying the *cdpA* mutations: Δ*cdpA*, *cdpA*_ΔNTD_, *cdpA*_ΔLinker_, and *cdpA*_ΔCTD_ (Supplementary Table 2).

### Construction of plasmids for gene expression in *H. volcanii*

The plasmids pTA962^53^ or pIDJL40^12^ (containing GFP) were used as the basis to construct plasmids for controlled expression of the *cdpA* genes or modified versions. To construct pTA962-*cdpA*, the *cdpA* ORF (HVO_0739) was amplified from *H. volcanii* H26 genomic DNA using primers HVO_0739_r and HVO_0739_r, then the product was cloned between the NdeI and BamHI sites of pTA962. The *cdpA* ORF was also amplified without its stop codon, then cloned between the NdeI and BamHI sites of pIDJL40, to create the *cdpA*-gfp fusion. For dual expression of *cdpA*-gfp and *ftsZ1/2*-mCherry, PvuII fragment of pTA962-*FtsZ1-mCherry* or *ftsZ2*-mCherry^13^ (containing a *p.tnaA*-*ftsZ*-mCherry gene fusion) were ligated into HindIII-cut (Klenow blunt-ended) pIDJL40_HVO_0739 containing (*cdpA*-GFP), creating plasmid with two independent copies of the CdpA-GFP and FtsZ1/2-mCherry.

For expression of *cdpA*-GFP under its native promoter, the *cdpA* ORF, along with the intergenic region of HVO_0739 and HVO_0738 (including native promotor of *cdpA*), was amplified using primers CdpA_nativeP_F and HVO_0739 (NS)_r, then the product was cloned between ApaI and BamHI of pIDJL40 (remove p.tnaA promoter).

To construct the CdpA_NTD_-GFP fusion, the N-terminal domain of CdpA (1-149 aa) was amplified from *H. volcanii* H26 genomic DNA using primers HVO_0739_f and CdpA_NTD_R, then the product was cloned between NdeI and BamHI sites of pIDJL40. The mutant CdpA without the C-terminal domain and without the N-terminal sequences were amplified from *H. volcanii* H26 and mutant *cdpA*_ΔNTD_ genomic DNA, respectively, by using the primers in Supplementary Table 1, and then cloned between NdeI and BamHI sites of pIDJL40, to create CdpA_ΔCTD_-GFP and CdpA_ΔNTD_-GFP fusions. To construct pHVID1-HA, synthetic DNA fragments encoding the HA tag with EcoRI and NotI overhangs were obtained from Integrated DNA Technologies (IDT). These fragments were subsequently cloned into the pHVID1 vector. To generate CdpA and FtsZ2 HA tagged plasmids, the respective genes amplified from pIDJL40_*cdpA* and pIDJL40- *ftsZ2* were cloned between NdeI and BamHI sites of pHVID1-HA plasmid, respectively. All plasmids were demethylated by passage through *E. coli* C2925 and re-purified before transfer to *H. volcanii* H26 by polyethylene glycol-mediated spheroplast transformation^52^.

### Phase-contrast and epifluorescence microscopy

A 2 µl sample of mid-log culture was loaded on a 1% agarose pad containing 18% buffered saltwater (BSW) on a glass slide at room temperature, and a #1.5 glass coverslip was placed on top^13^. The images were acquired on a GE DeltaVision Elite microscope with a pco.edge 5.5 sCMOS camara and an Olympus 100X Plan Apo 1.4 NA objective using a FITC filter set (ex. 464-492 nm, em. 500-549 nm) or an mCherry filter set (ex. 557-590, em. 598-617 nm).

The analysis of cell shape (area vs circularity), FtsZ1-mCherry intensity ratio (localized/diffuse fluorescence), and FtsZ1-mCherry localization thickness measurements were performed as described^13^. For counting the number of FtsZ2-GFP rings per cell, the number of correctly positioned rings were manually counted to exclude the abnormal localizations, and cell length was determined by a custom script in FIJI^13^.

The measurement of localized and diffuse FtsZ1- and FtsZ2-FP fluorescence areas were analyzed in MATLAB 2019b (MathWorks). The phase-contrast images were first pre-processed using a bottom-hat transform (*imbothat* function) with a disk-shaped structuring element of a radius of 30 pixels for the kernel. These were binarized and individual cells were segmented (*bwconncomp* function). Cell debris were excluded by removing objects with area less than 1 μ*m*^2^ (∼237 pixels). The filtered cell outline was parametrically smoothed by fitting each component of cell outline (i.e., x- and y-components in the Cartesian coordinate system) with two Fourier expansions up to the 8th order mode. The corresponding fluorescence images were pre-treated by subtracting the average intensity of the background (i.e., excluding regions of the detected cells). Additionally, a top-hat transform (*imtophat* function) was applied with a disk-shaped structuring element of a radius of 10 pixels. FtsZ1 and FtsZ2 clusters within each cell were identified and segmented by converting the processed fluorescence images to 8-bit and applying the extended-maxima transform (*imextendedmax* function) with a threshold value of 1. Small fluctuations in FtsZ1-FP and FtsZ2-FP fluorescence signals were neglected by removing clusters less than 4 pixels. The localization area of FtsZ1 and FtsZ2 in each cell was then defined as the total area of all FtsZ1 or FtsZ2 clusters found within the cell outline. The ratio of FtsZ2 localization area to FtsZ1 localization area was then determined for each cell. Cells were excluded from the analysis if either FtsZ1 or FtsZ2 clusters were undetectable within the cell or if the cell exhibited a signal-to-noise ratio of less than 2.

CdpA-GFP localization along the medial axis in filamentous SepF depleted cells was analysed using the same segmentation approach described above with MATLAB. After rolling ball background subtraction (radius: 20 pixels, ∼1.4 µm), CdpA-GFP intensities were mapped along the medial axis by averaging pixel intensity along the transverse direction and smoothed using a moving average filter (window size: 10 pixels, ∼0.7 µm). Peak localizations of CdpA were identified using the *findpeaks* function and manually corrected to avoid foci or other aberrant structures.

### Time-lapse Microscopy

The sample was prepared by using submerged sandwich technique as previously described^13^. A 2 µl sample of mid-log culture was placed on a 0.3% w/v agarose pad (containing the medium for the growth) prepared on an 8-mm glass-based (#1.5) FluoroDish (WPI). A 4 mL volume of medium was added to the FluoroDish to cover the agarose pad assembly, and the lid was placed on the FluoroDish to minimize evaporation. The images were captured at 42°C every 10 min for 24 h on a Nikon TiE2 inverted fluorescence microscope with a 100 X oil-immersion phase-contrast NA 1.45 objective.

### 2D- and 3D-Structured Illumination Microscopy (SIM)

Samples were prepared on agarose pads as described above for phase-contrast and epifluorescence microscopy. Images were captured on a Delta Vision OMX SR 3D Structured Illumination Microscope with a 60 X Plan 1.42 NA Oil Immersion Objective at room temperature for visualizing fluorescent fusion proteins, or at 42°C for 3D-SIM time-lapse imaging. GFP and mCherry signals were collected through a FITC and a TRITC filter, respectively. 3D-SIM images were sectioned using a 125 nm z-step. The reconstructed images-stacks were 3D rendered and visualized by using IMARIS x64 9.6.0 (Bitplane Scientific). The maximum diameters of FtsZ2-GFP nano-rings were manually measured using the “Measurement Points” function in IMARIS. 3D-SIM time series (acquired every 30 s – 2 min as required) were compiled from single time points using IMARIS to create a time-lapse series, followed by bleach correction (Histogram Match) and registration (using ‘translation’ in TurboReg) in Image J 1.53v. The 3D intensity plots were generated by using ‘3D surface plot” in ImageJ 1.53v. Two regions of interest were monitored overtime show how FtsZ-GFP or CdpA-GFP fluorescence fluctuates in the Z ring.

### Total Internal Reflection fluorescence (TIRF) microscopy

Samples were prepared as described for time-lapse microscopy. The TIRF imaging was performed using a Nikon TiE-2 N-STORM microscope operated in TIRF mode on a stage top incubator (OKO-lab) set to 42°C, equipped with a 100 X 1.49 NA TIRF objective. The TIRF angle was set to optimize the signal to noise ratio at a fixed angle of 100°. The imaging settings were chosen to minimize photobleaching with the power intensity set at 5% and the exposure time was 150 – 200 ms. FtsZ1- GFP was imaged at 4 s intervals for 15 min, and FtsZ2-GFP were imaged at 10 s intervals for 40 min. CdpA-GFP was imaged either at 4 second intervals for 15 min or 10 second intervals for 40 min.

For the analysis of TIRF data^5^, fluorescence images were stabilized, and registered (StackReg, Translation), and corrected for photobleaching (simple ratio), interpolated to 20 nm/pixel via the bicubic method in Image J, and moving averaged over a 12-frame window of time. The fluorescence intensity of an FtsZ polymer along the direction of its movement in each frame was determined from the intensity along a line with a width of 25 interpolated pixels (∼ 500 nm) manually drawn across the full length of the path of the FtsZ polymer. This line was then used to plot the kymograph (line width = 25) then background subtraction rolling ball =25, and smooth (for display purpose). The speeds of clusters were calculated by manually measuring the slopes of leading and trailing edges of clear fluorescence zigzags in the kymograph. The distributions of speed of FtsZ-FP clusters were plotted in OrginPro (2021). For analysis of Z-ring intensity fluctuations during TIRF time-lapse imaging, briefly, a cell of interest was cropped out from both fluorescence and bright-field images. Second, the cell was rotated to align the Z-ring horizontally. Fluorescence background within the cell was calculated by averaging the total integrated intensity over the area defined by the cell contour. A square region of interest (ROI) with 3×3 pixels in the centre of the ring was then chosen manually. The corrected intensity was calculated as the integrated intensity within the ROI after background subtraction and smoothed by a moving-average filter with a window size of 5 time points in OriginPro (2021).

The Power Spectral Density (PSD) analysis was conducted using custom MATLAB scripts. Firstly, a cell of interest was cropped out from both fluorescence and bright-field images. The fluorescence images were then registered to the bright-field image and corrected for sample/stage drift and photobleaching. Following this, the cell was rotated manually to align the Z-ring horizontally. To determine fluorescence background within the cell, the total integrated intensity over the area defined by the cell outline was averaged. A square region of interest (ROI) with 3×3 pixels at the center of the ring was then chosen manually. The local FtsZ-GFP (*I*) was computed as the integrated intensity within the ROI after background subtraction.

Using fluorescence intensity time trace, *I*(*t*), from the center of the division ring, the PSD curves were calculated. This involved subtracting the mean value 〈*I*(*t*)〉 from *I*(*t*) and subsequently applying the Fast-Fourier Transform (FFT) to the shifted time trace (fft function in MATLAB). For each cell, the PSD curve across different frequencies *f* was then calculated as follows:

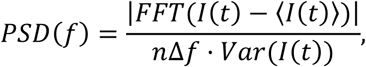

where *n* is the total number of frames of the trace and Δ*f* is the frame rate (1/4 *s*^−1^). Note that the PSD was normalized by the squared magnitude of the intensity trace (variance of *I*(*t*)). The mean PSD curves were generated by averaging the PSD curves of individual cells in each strain.

### Immunoprecipitation of HA-tagged CdpA

The method was based on that described^14^ but with modifications. H26-HA and CdpA-HA strains were grown in 300 mL of Hv-cab medium supplemented with 2 mM Trp until their OD_600_ reached 0.5. The cells were harvested by centrifugation at 6,000 x g for 15 min. The pelleted cells were resuspended in 30 mL of 18% buffered salt water supplemented with 10 mM HEPES. For crosslinking 1.5% (w/v) formaldehyde was added to the cell suspension, and the mixture was incubated at room temperature for 10 min with constant shaking. The crosslinking reaction was stopped by adding 20 mL of 18% buffered salt-water containing 1.25 M glycine. The cells were centrifuged again for 15 min at 6,000 x g. The cell pellet was washed twice with 18% buffered salt water with 1.25 M glycine. After the final wash, the cell pellet was resuspended in 3 mL of lysis buffer (1X PBS, 10 mM MgCl2, 10 µg/ml DNase I and protease inhibitor cocktail). The cells where lysed by sonication for 5 min, with 5 sec on and 5 sec off cycle at 35% amplitude. To remove the cell debris, lysed cells were centrifuged at 6,000 g for 15 min at 4 °C in a microcentrifuge. The supernatant, expected to contain cytosolic and fragmented membrane fractions, was used for immunoprecipitation with washed anti-HA magnetic beads (Thermo Scientific) as described in the manufacturer’s instructions. Briefly, the supernatant (1 mL) was added to 0.5 mg of anti-HA beads and incubated at room temperature for 30 min with gentle mixing. The beads were washed twice with buffer (Tris–buffered saline with 0.05% Tween-20) and proteins were eluted with 0.1 M glycine pH 2.0, and then neutralized and denatured by boiling for 5 min with SDS sample buffer, which was also expected to disrupt crosslinking.

### Western blotting

Proteins were separated by sodium dodecyl-sulphate polyacrylamide gel electrophoresis (SDS-PAGE) and then electro-transferred to a nitrocellulose membrane using a Bio-Rad trans-blot turbo transfer system. The membrane was blocked overnight in 5% (w/v) skim milk in Tris-buffered saline with 0.05% Tween-20 (TBST). A primary antibody recognizing heamagglutanin (HA) (Thermo Fisher Scientific, Cat# 26183) (1:10,000), FtsZ1^13^ (1:1,000) and FtsZ2^13^ (1:2,000) were incubated with membrane for 2 h with constant shaking at RT. Following that the membrane was washed three times, 5 min each with constant shaking at RT, with TBST. The membrane was then incubated with the secondary antibody against mouse (Goat anti-mouse IgG-HRP; AbCam 6789) for 1 h in a 1:5,000 dilution. The membrane was washed five times (5 min each) with TBST before detection. Bands were detected with enhanced chemiluminescence reagents (Thermo-Fisher) using an Amersham Imager 600 system.

### Mass spectrometry (MS)

Fifty micrograms of eluted proteins from the pull-down assay were subjected to trypsin digestion using Promega Trypsin Gold, mass spectrometer grade (1 µg/µL) in 50 mM acetic acid. The digested samples were processed for clean-up using solid-phase extraction (SPE). An equivalent amount of protein (15 µg) for each sample was loaded onto the SPE columns, which were then washed, and peptides eluted. Columns were manually prepared using Empore SDS-RPS (Styrenedivinylbenzene reverse phase sulfonate) with a pipette tip. The SDB-RPS based tips were used for peptide clean up.

Eluted samples were dried in a vacuum centrifuge, and peptides were reconstituted in the injection solvent suitable for the LC/MS system. Analysis was performed on a Q Exactive Plus mass spectrometer (Thermo Fisher Scientific), and the MS/MS data files were analyzed using PEAKS Studio against *Haloferax volcanii* predicted proteome database using Uniprot. The MS peak areas obtained for the peptides were then used for statistical analyses.

### Data availability

The data generated in this study, including microscopy image analyses and western blots are provided in the Supplementary Information and Source data files with this paper. The mass spectrometry proteomics data have been deposited to the ProteomeXchange Consortium via the PRIDE^54^ partner repository with the dataset identifier PXD062651. Other data and biological materials are available from the corresponding author upon reasonable request.

## Acknowledgments

This study was supported by the University of Technology Sydney Chancellor’s Research Fellowship (PR021-13161 to Y.L.) and by the Australian Research Council (FT160100010 to I.G.D, DP220101143 to B.S., and DP250100999 to I.G.D & Y.L.). For technical and facilities management support, we thank Matthew Padula from the Proteomics Core Facility, UTS, Pauline Coulon for assistance with proteomics data handling, Matthew Pittorino for discussion of TIRF image analysis, and Amy Bottomley, Christian Evenhuis, and Louise Cole from the Microbial Imaging Facility, UTS. We thank Sonja-Verena Albers at the University of Freiburg for providing the sepF depletion strain (HTQ239) and plasmid for expression of sepF-GFP (pSVA3942).

## Author Contributions Statement

Y.L and I.G.D. designed the research. Y.L. constructed most of the plasmids, strains and conducted the primary microscopy imaging and analyses. V.D.S constructed plasmids and strains for pull-down experiments and performed immunoprecipitation, western blot and mass-spectrometry analyses. D.H performed phylogeny analysis of CdpA and CdpA-like sequences and contributed the production of some image analysis figures using R programming. Z.X. performed PSD analysis and quantified the localization area of FtsZ1/FtsZ2 and ring number of CdpA-GFP in HTQ239 strain in MATLAB. B.S. contributed to the TIRF imaging. K.A.M, and I.G.D. performed Alphafold2 Multimer prediction and analysis. Y.L. and I.G.D. interpreted results and wrote the original draft. All authors reviewed and gave scientific input to revise the manuscript. Y.L and I.G.D managed and supervised the project and acquired funding.

## Competing Interests Statement

The authors declare no competing interests.

## Notes

### Competing Interest Statement

The authors have declared no competing interest.

### Summary of Updates

The main text has been revised to include a new figure along with the corresponding results and discussion section. Supplemental files have also been updated accordingly.

