## Supplementary Information for "Cell division protein A (CdpA) organises and anchors the division ring at midcell in haloarchaea"

#### Supplementary Figures

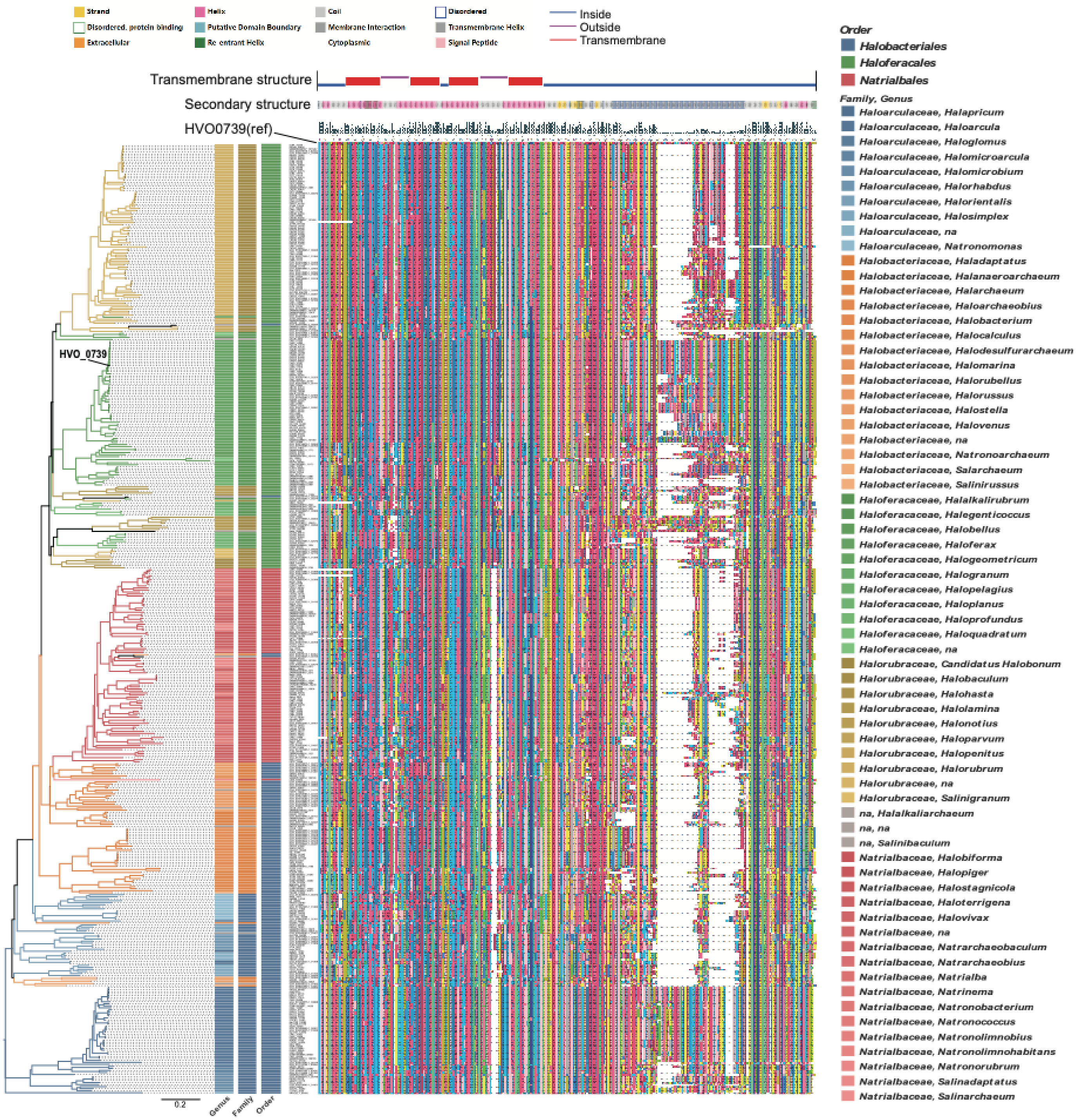


**Supplementary Figure 1. Phylogenetic tree and conserved regions of 470 CdpA amino-acid sequences.** The multiple-sequence alignment only shows positions aligned with the reference sequence HVO_0739 to show consensus regions of CdpA. The annotation of the reference sequence is shown above, including transmembrane structure, secondary structure, and consensus level of the alignment. A neighbour-joining tree based on the multiple sequence alignment is shown at left; the taxonomy of the source genome of each protein sequence was used to annotate the tree using the indicated colour coding at order, family, and genus levels.

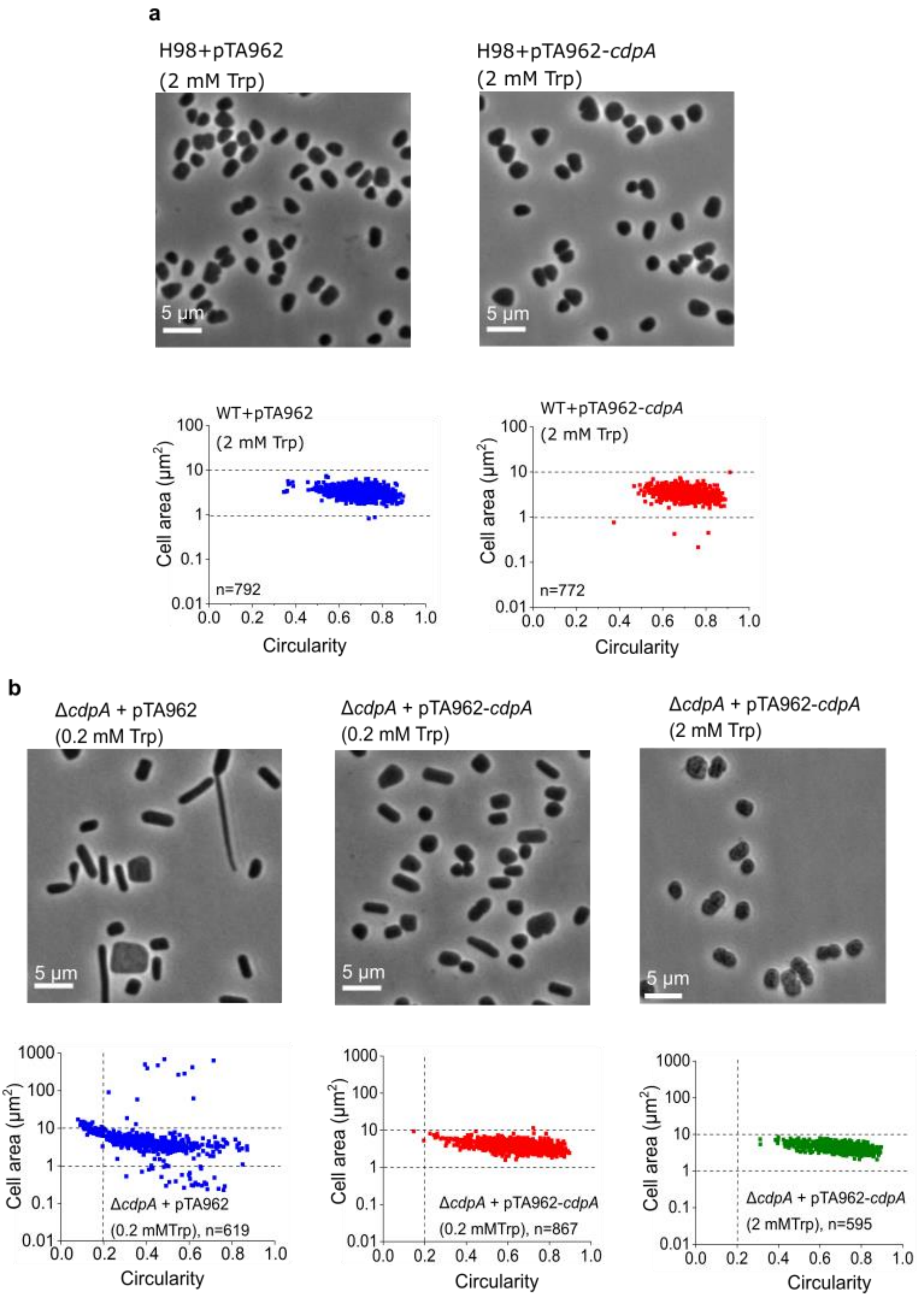


**Supplementary Figure 2. Overproduction of CdpA in the wild-type background and complementation of the *cdpA* cell division phenotype in the Δ*cdpA* strain. a,** Phase-contrast micrographs and corresponding cell size/shape analysis of the *H. volcanii* wild type (H98 + pTA962) and CdpA overproduction (H98 + pTA962-*cdpA*) strains from steady mid-log cultures in Hv-Cab medium supplemented 2 mM Trp. Scale bars, 5 µm. **b,** Phase-contrast micrographs and corresponding cell size/shape analysis of the *H. volcanii* CdpA complementary (Δ*cdpA* + pTA962-*cdpA*) and control (Δ*cdpA* + pTA962) strains from steady mid-log cultures with Hv-Cab medium supplemented 0.2 mM Trp or 2 mM Trp. Scale bars, 5 µm. n=number of cells examined over one biologically independent experiment. The data shown is representative of at least two independent experiments. Source data are provided as a Source Data file.

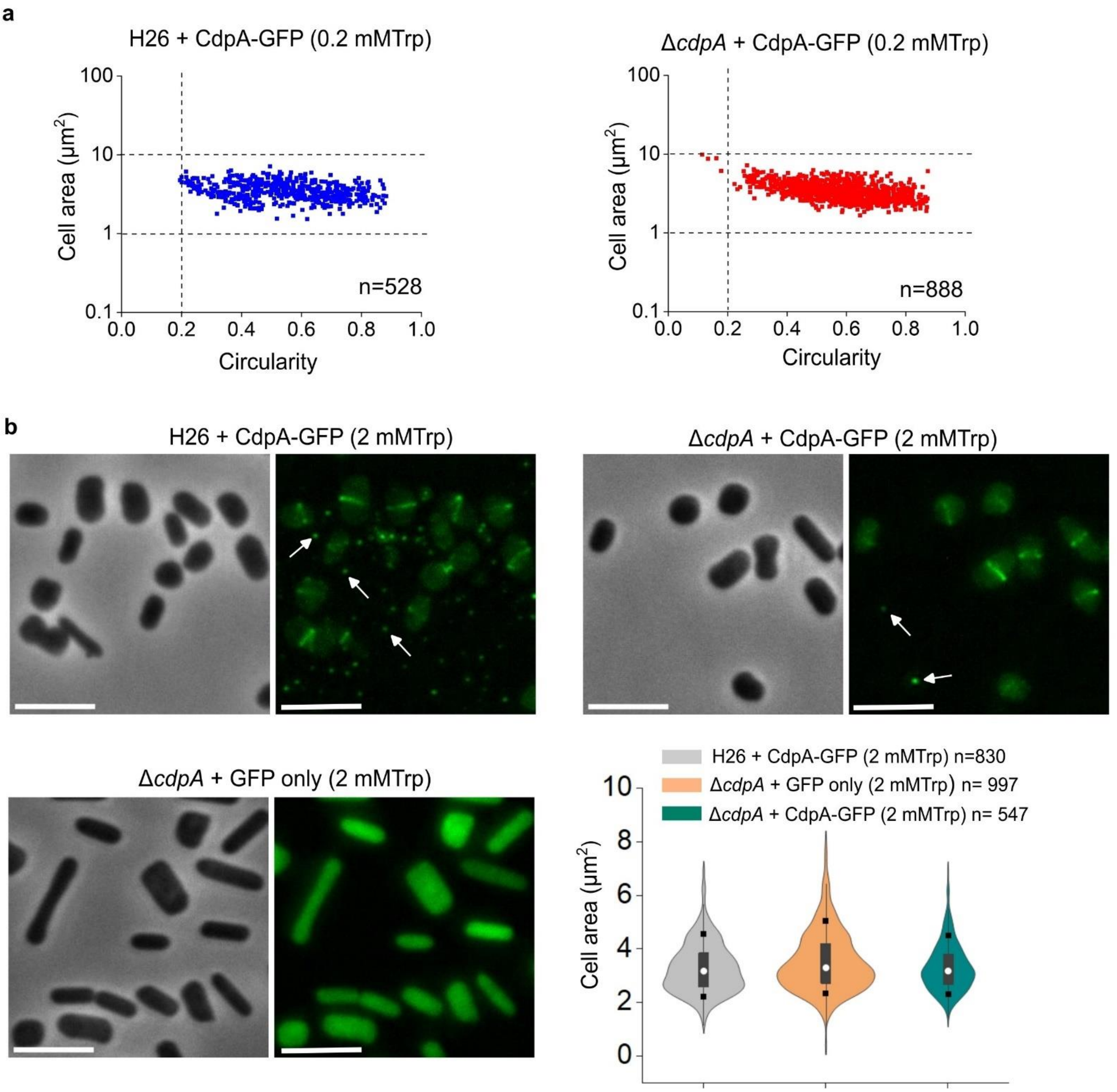


**Supplementary Figure 3. Expression of CdpA-GFP in wild-type and Δ*cdpA* strains. a,** Cell area and shape (circularity) were determined for individual cells, as per Fig. 1 e and f (0.2 mM Trp). **b,** Phase-contrast and fluorescence micrographs of cells producing CdpA-GFP in wild-type and Δ*cdpA* backgrounds. The expression of GFP only from pTA962 in Δ*cdpA* strains showed diffusion localization. White arrows indicate the extracellular particles. Scale bars, 5 µm. Violin plots of the cell area of the three strains were shown; the median is indicated by a white dot, the thick bar represents the interquartile range (IQR), the thin grey line indicates 1.5 times the IQR, and the black square boxes represent $10^{th}$-$90^{th}$ percentile range. n=number of cells examined over one biologically independent experiment. The data shown is representative of at least two independent experiments. Source data are provided as a Source Data file.

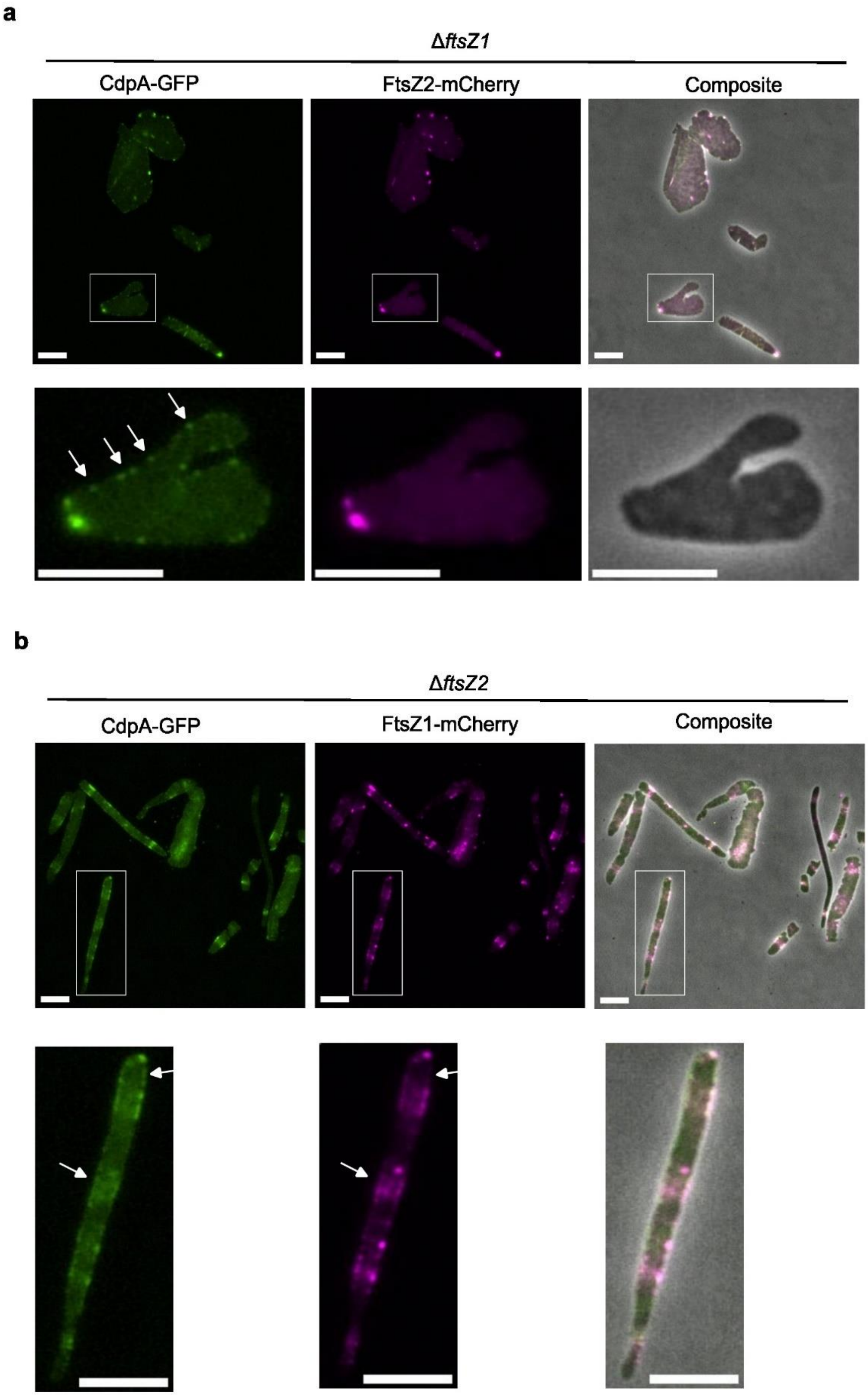


**Supplementary Figure 4. Co-localization of CdpA-GFP and each FtsZ-mCherry in the absence of the other FtsZ. (a)** *H. volcanii* Δ*ftsZ2* producing both CdpA-GFP and FtsZ1-mCherry. **(b)** *H. volcanii* Δ*ftsZ1* producing both CdpA-GFP and FtsZ2-mCherry. The cells were grown in Hv-Cab medium with 0.2 mM Trp and sampled for imaging during mid-log phase. The bottom panel shows magnified views of the insets from the upper panel. The bottom panels shows magnified views of the insets from the upper panels. In both strains, CdpA-GFP and FtsZ-mCherry showed considerable overlap but not fully co-localized. White arrows highlight regions where CdpA-GFP and FtsZ1- or FtsZ2-mCherry signals did not co-localize. All scale bars, 5 µm.

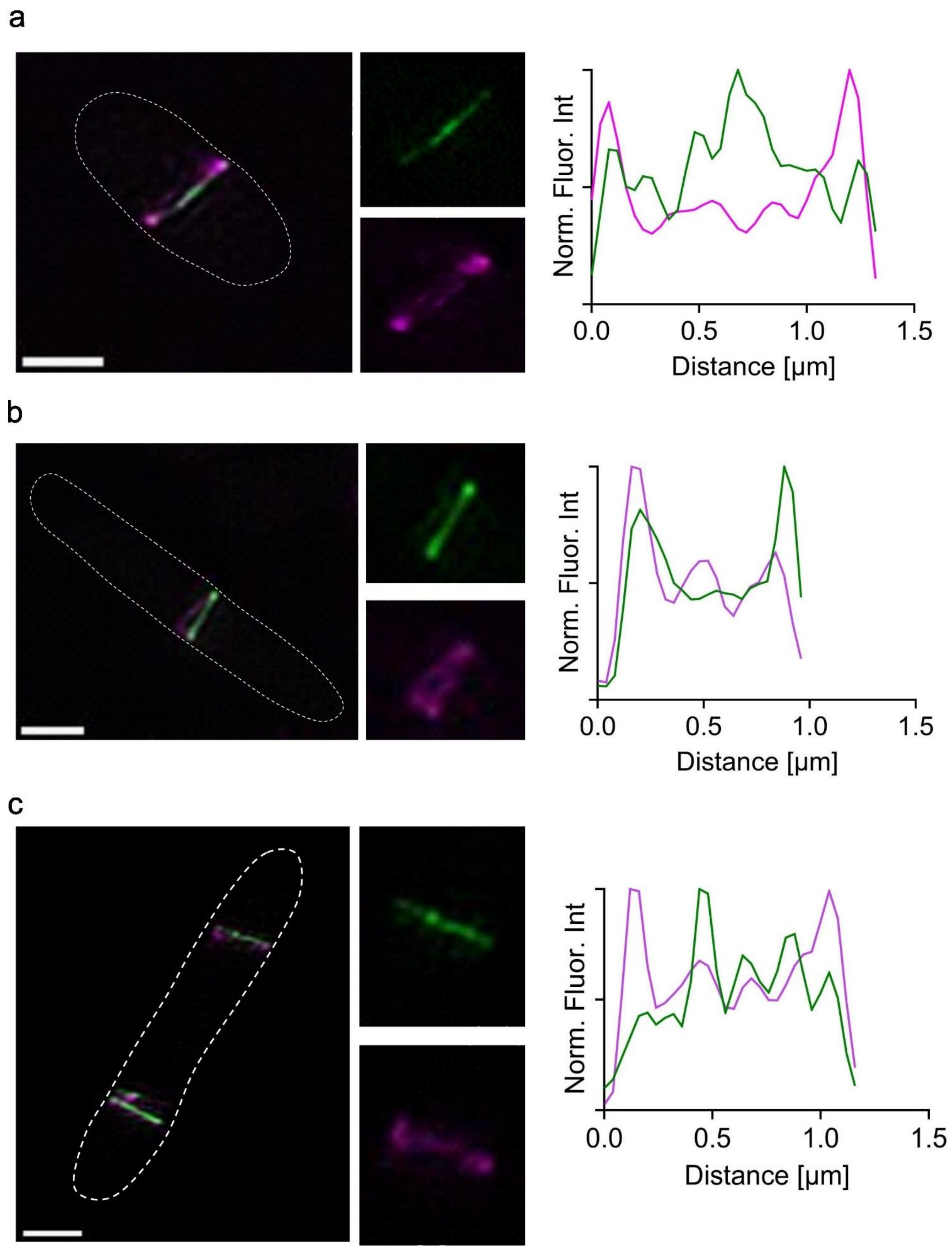


**Supplementary Figure 5. Three examples of 2D-SIM images of *H. volcanii* H26 expressing both FtsZ1-mCherry and FtsZ2-GFP. during mid-log growth with 0.2 mM Trp.** The cells were grown in Hv-Cab medium with 0.2 mM Trp and sampled for imaging during mid-log phase. mCherry (magenta) and GFP (green) fluorescence was quantified by plotting normalized intensity in the respective channel along the ring band. In panel c in a cell with two cell division rings, the fluorescence intensity of upper ring band was quantified. All scale bars, 1 µm. Source data are provided as a Source Data file.

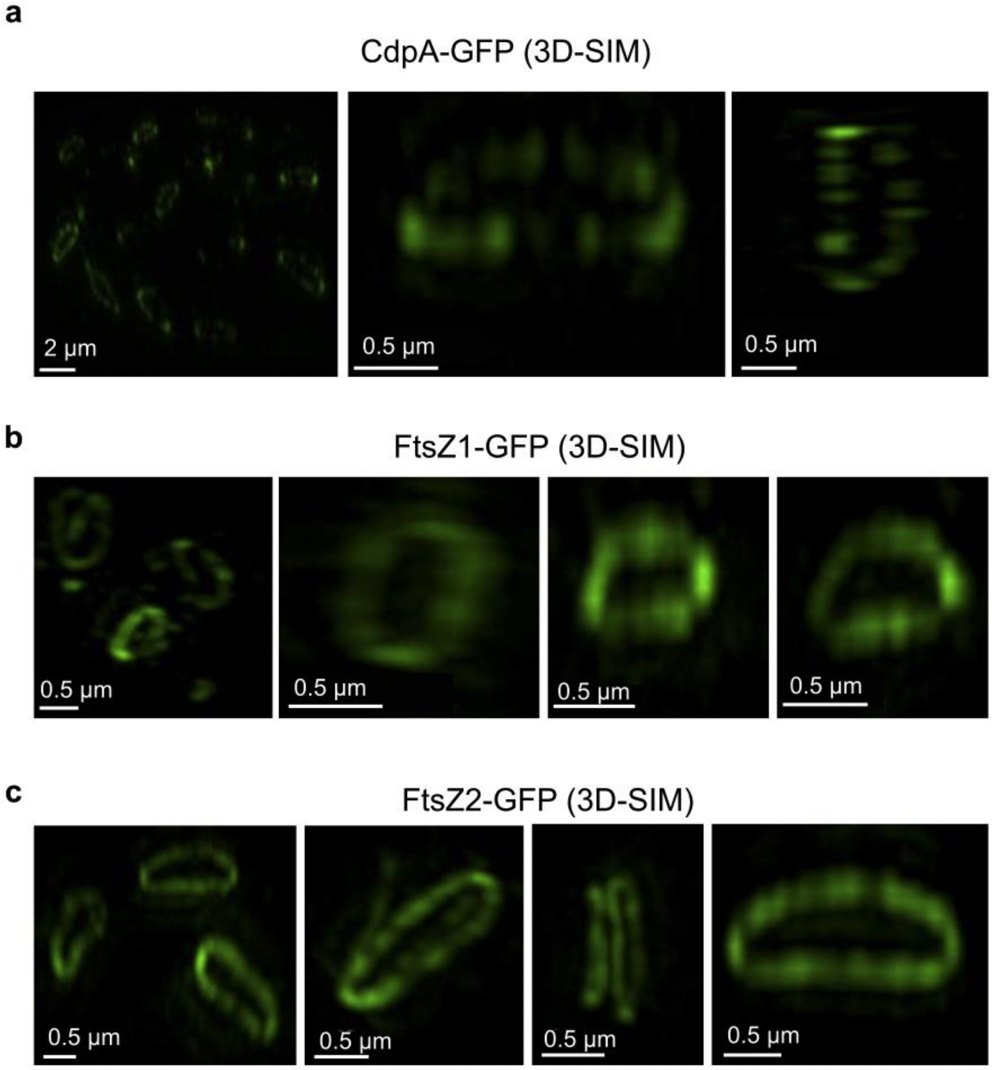


**Supplementary Figure 6. Examples of 3D-SIM images of *H. volcanii* H26 backgrounds expressing CdpA-GFP, FtsZ1-GFP, or FtsZ2-GFP (as per Fig. 2c) during mid-log growth with 0.2 mM Trp.** The distribution of CdpA in the Z ring was patchier and more discontinuous compared with that of FtsZ1 and FtsZ2.

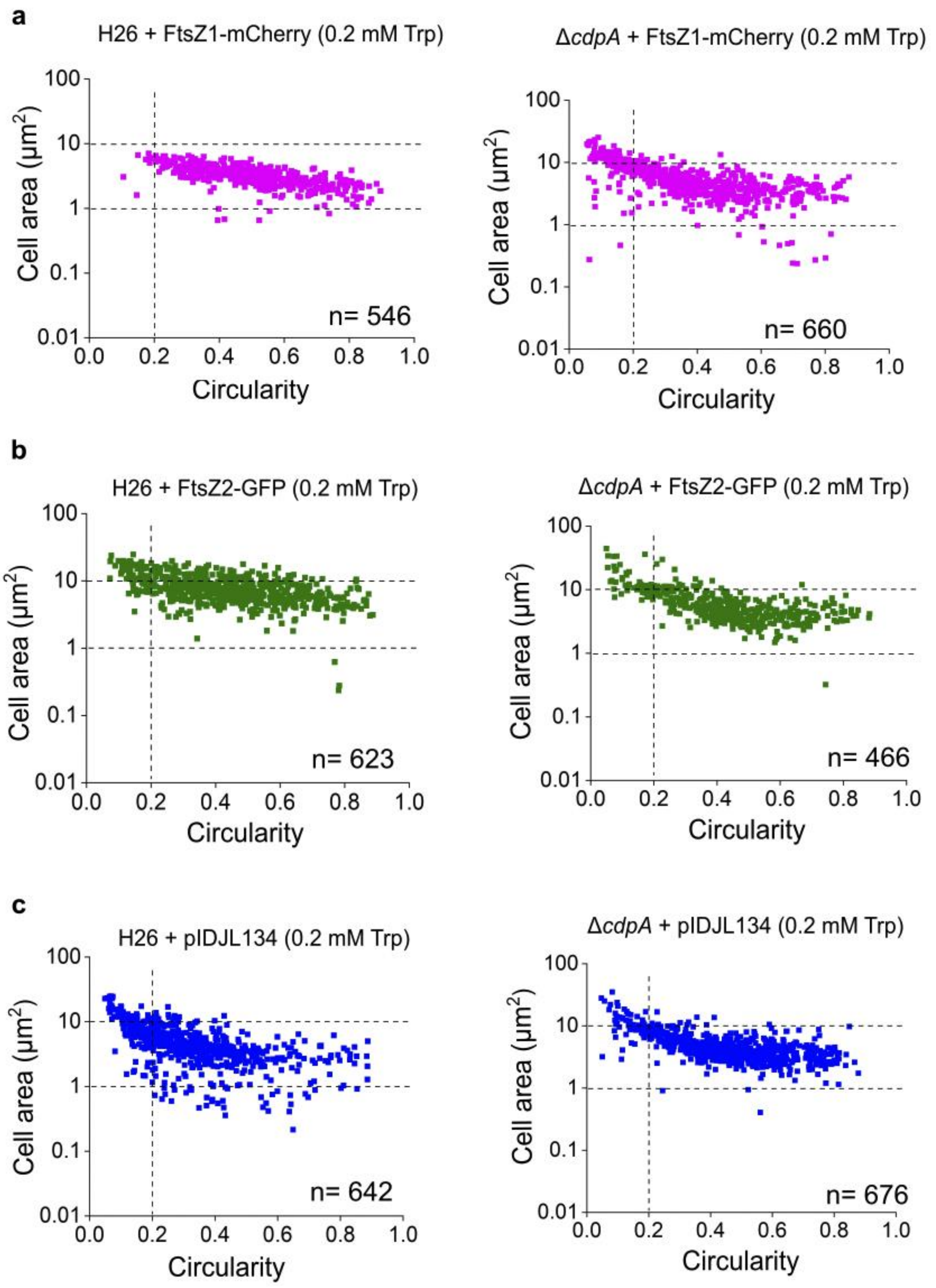


**Supplementary Figure 7. Cell size and morphology analysis of expression of FP proteins in wild-type and Δ*cdpA* strains.** Cell shape quantification scatterplots (circularity vs cell area) were generated from analysis of phase-contrast images of the indicated mid-log strains, cultured with induction under 0.2 mM Trp. n=number of cells examined over one biologically independent experiment. **a,** FtsZ1-mCherry expression in wild-type H26 and Δ*cdpA* strains. **b,** FtsZ2-GFP expression in wild-type H26 and Δ*cdpA* strains. **c,** Expression of FtsZ1-mCherry + FtsZ2-GFP (from pIDJL134) in wild-type H26 and Δ*cdpA* strains. n = number of cells examined over one biologically independent experiment. The data shown is representative of at least two independent experiments. Source data are provided as a Source Data file.

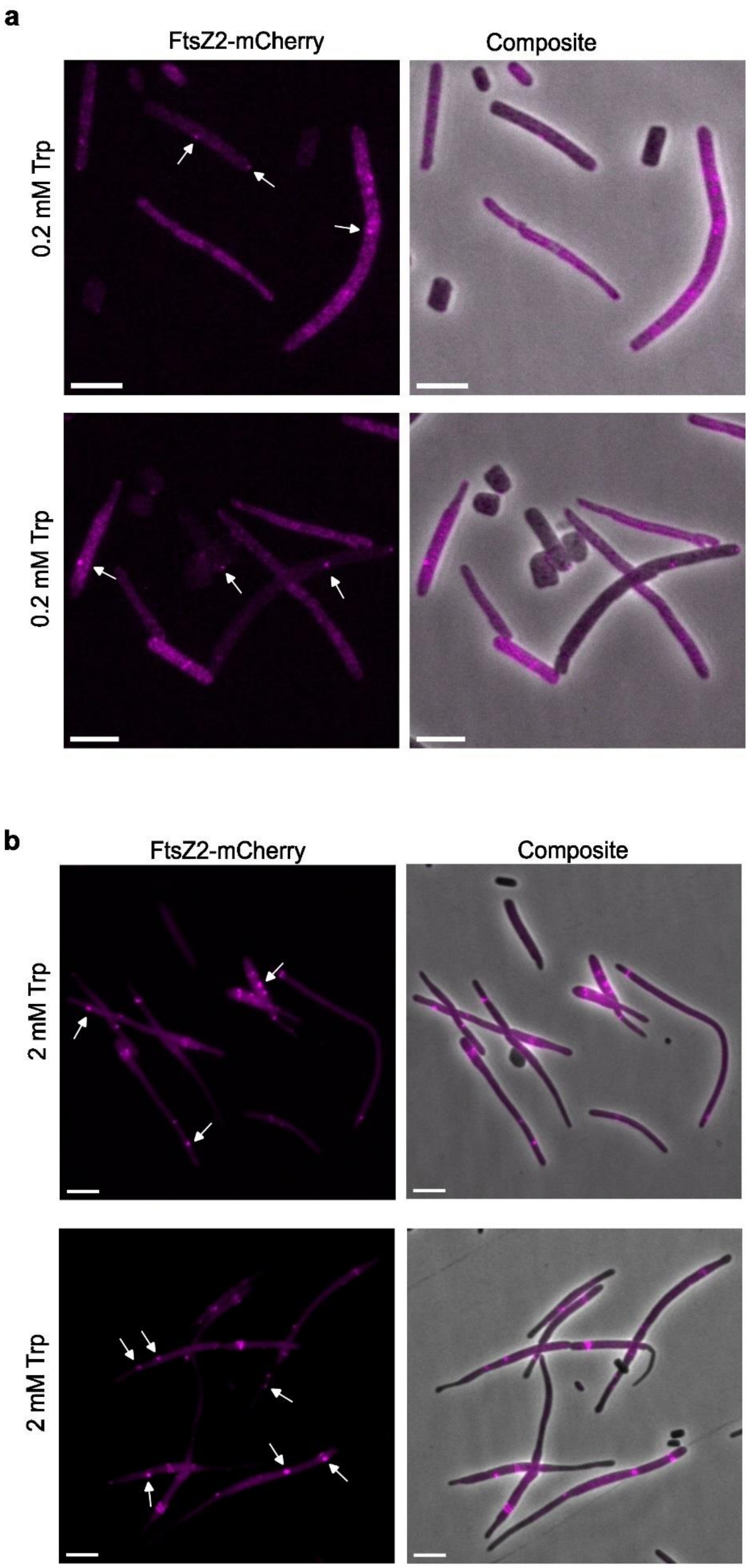


**Supplementary Figure 8. Examples of microscopy images of Δ*cdpA* cells expressing FtsZ2-mCherry.** The cells were grown in Hv-Cab medium with 0.2 mM Trp **(a)** or 2 mM Trp **(b)** and sampled for imaging during mid-log phase. At 0.2 mM Trp, FtsZ2-mCherry expression was low; however, weakly mispositioned, perpendicular, or focal FtsZ2-mCherry structures were still observed. At 2 mM Trp induction, where FtsZ2-mCherry expression was higher, the mispositioned structures of FtsZ2-mCherry became more prominent. White arrows indicate mispositioned localizations of FtsZ2-mCherry. All scale bars, 5 µm.

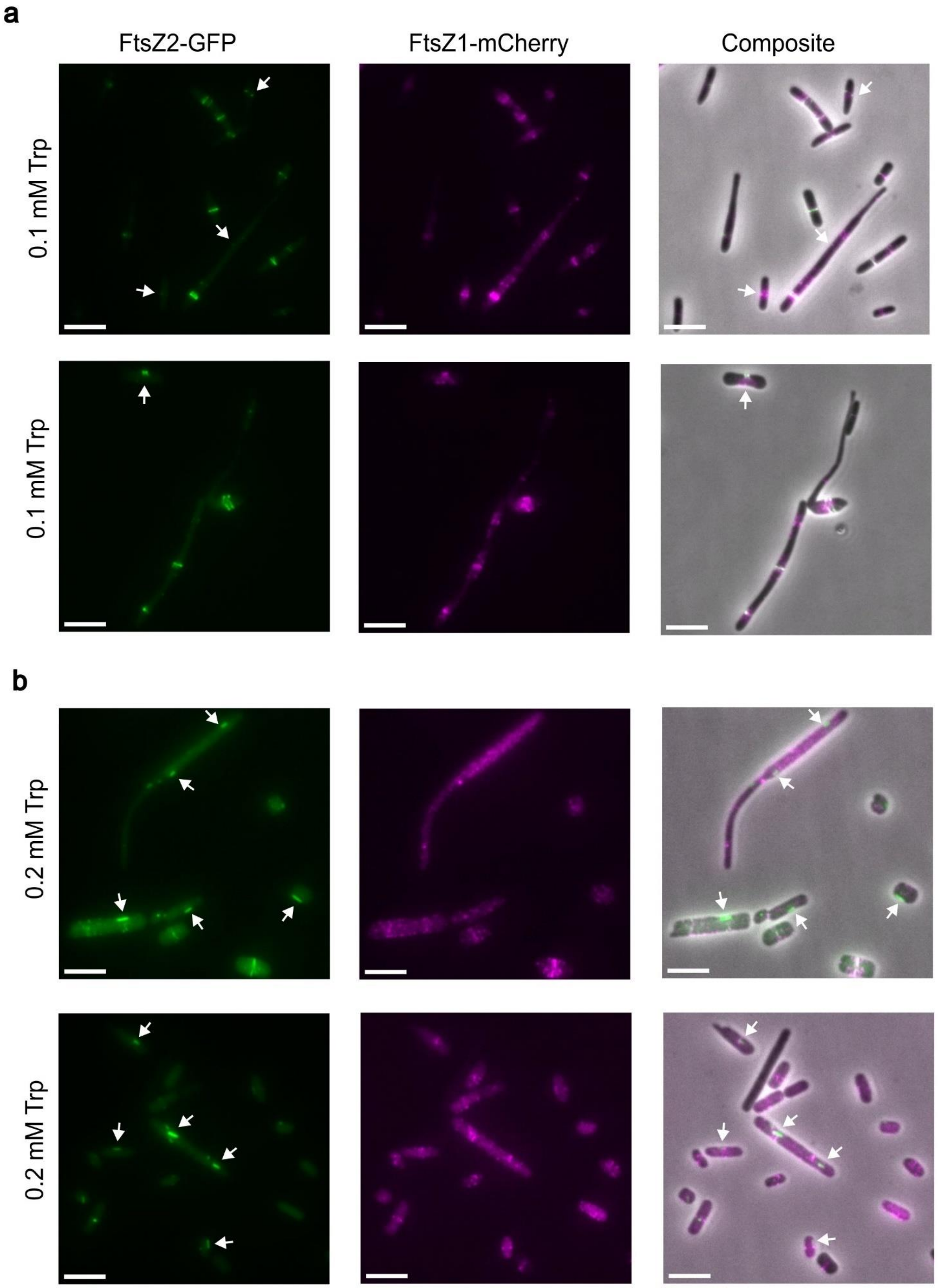


**Supplementary Figure 9. Examples of microscopy images of Δ*cdpA* cells expressing both FtsZ1-mCh and FtsZ2-GFP grown under (a) 0.1 mM tryptophan induction and (b) 0.2 mM tryptophan induction, respectively.** The cells were grown in Hv-Cab medium with 0.1 mM Trp or 0.2 mM Trp and sampled for imaging during mid-log phase. White arrows indicate the foci or mispositioned localization of FtsZ2-GFP. All scale bars, 5 µm.

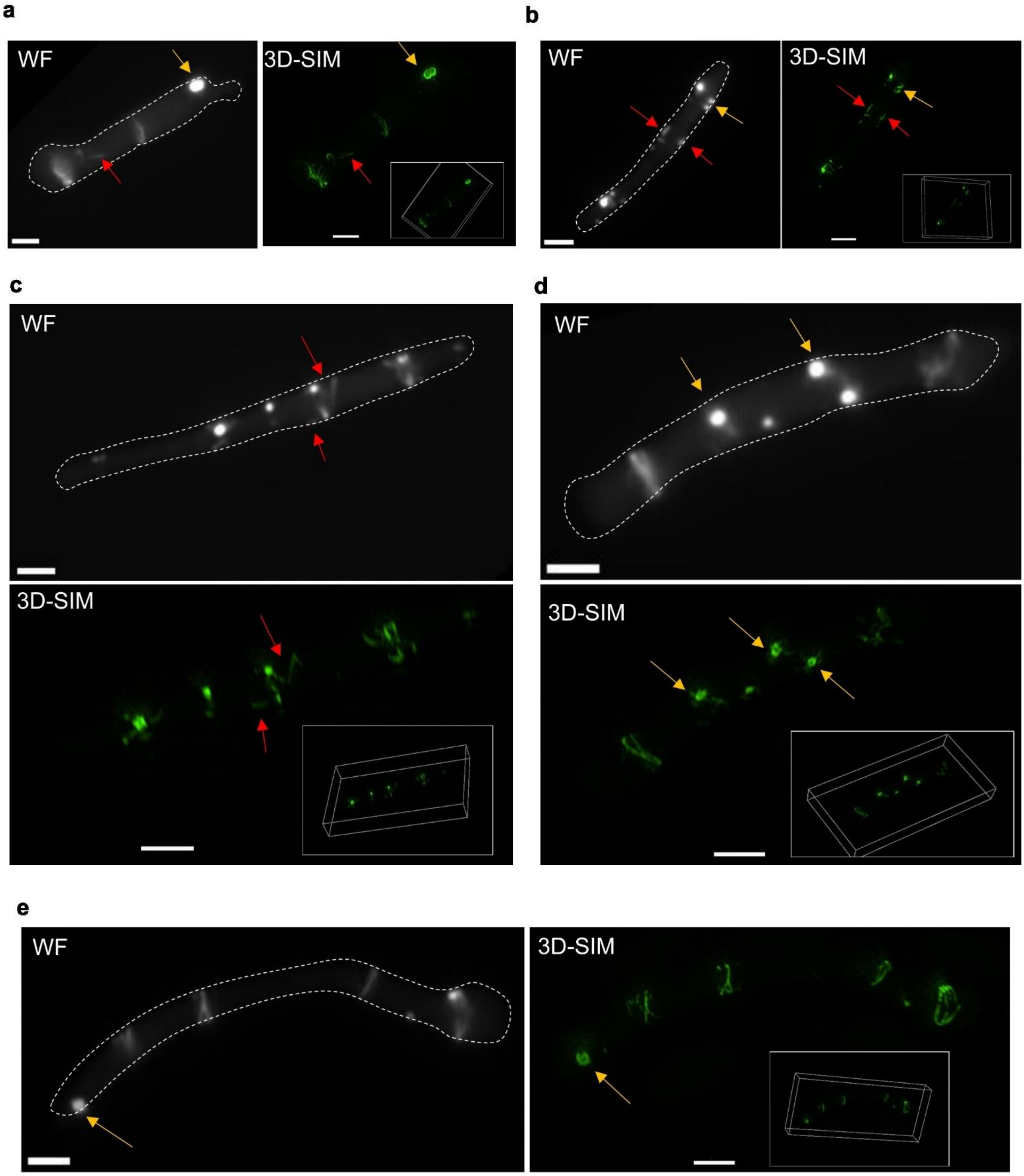


**Supplementary Figure 10. Comparison of conventional widefield fluorescence (WF) and 3D-SIM images of Δ*cdpA* cells producing FtsZ2-GFP.** The cells were grown in Hv-Cab medium with 0.2 mM Trp and sampled for imaging during mid-log phase. The corresponding wild-field images were shown alongside 3D-SIM images. The yellow arrows indicate the foci structures in WF but formed mini-rings or small portions of ring-like structures near cell membrane in 3D-SIM. The red arrows indicate perpendicular patches seen in WF corresponded to filamentous structures in 3D-SIM. All scale bars, 2 µm.

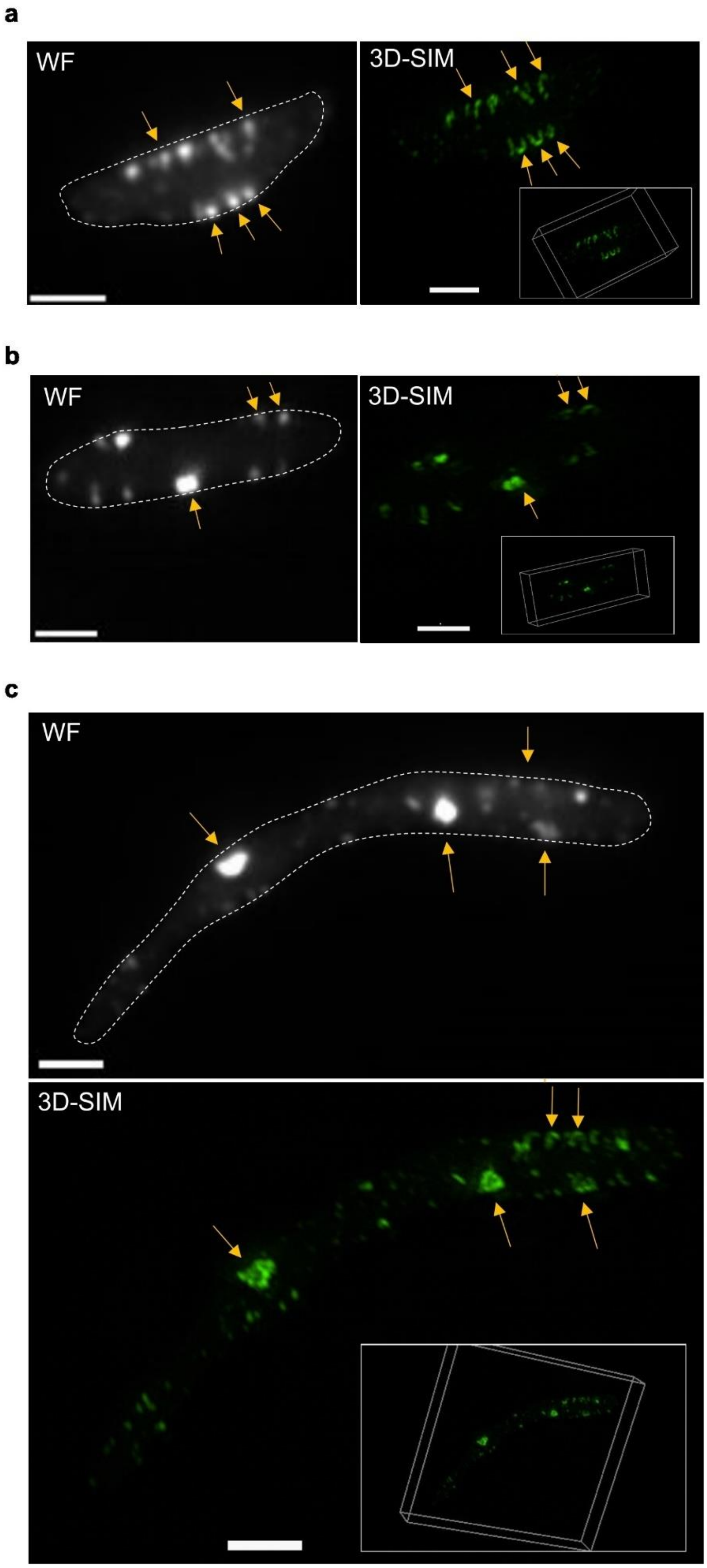


**Supplementary Figure 11. Examples of 3D-SIM images of Δ*cdpA* cells expressing both FtsZ1-mCherry and FtsZ2-GFP.** The cells were grown in Hv-Cab medium with 0.2 mM Trp and sampled for imaging during mid-log phase. Only FtsZ2-GFP were visualized by 3D-SIM due to the quick bleaching of FtsZ1-mCherry. The corresponding wild-field images were shown alongside 3D-SIM images. The yellow arrows indicate the foci structures in WF but formed nano-rings or small portions of ring-like structures near cell membrane in 3D-SIM. All scale bars, 2 µm

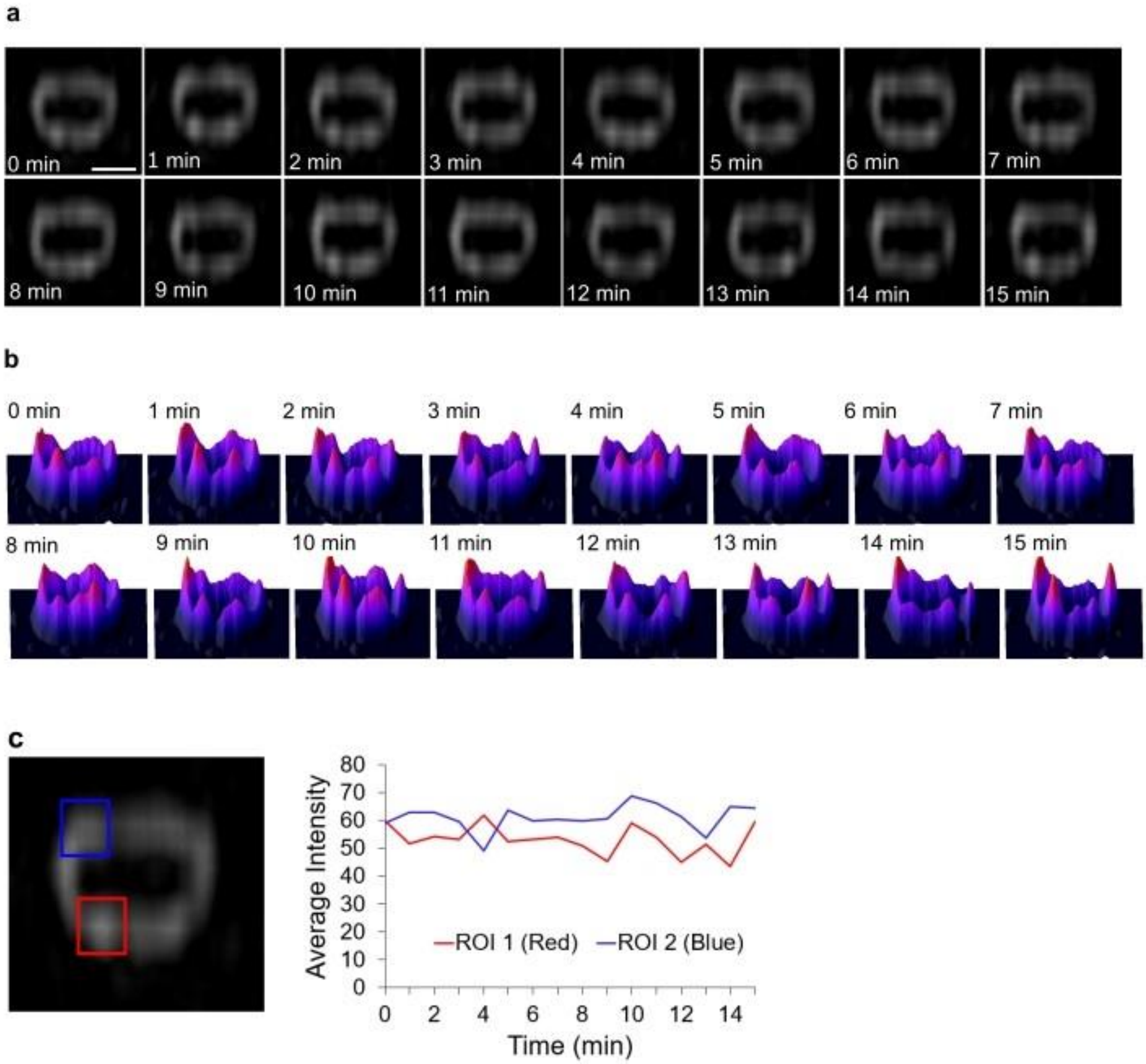


**Supplementary Figure 12. 3D-SIM Time-lapse analysis of FtsZ1-GFP localization within the division rings of *H. volcanii*. a,** Changes in the distribution of FtsZ1-GFP were evident in the regions of the Z ring in the *H. volcanii* YL 24 strain grown in Hv-Cab medium supplemented with 0.2 mM Trp at 42 °C. Scale bar, 0.5 µm. **b,** 3D intensity plots (fire LUT) show that the distribution of FtsZ in the Z ring remained heterogeneous and dynamic. **c,** Two regions of interest were monitored over time, showing fluctuation of FtsZ-GFP fluorescence signal in the Z ring. Source data are provided as a Source Data file.

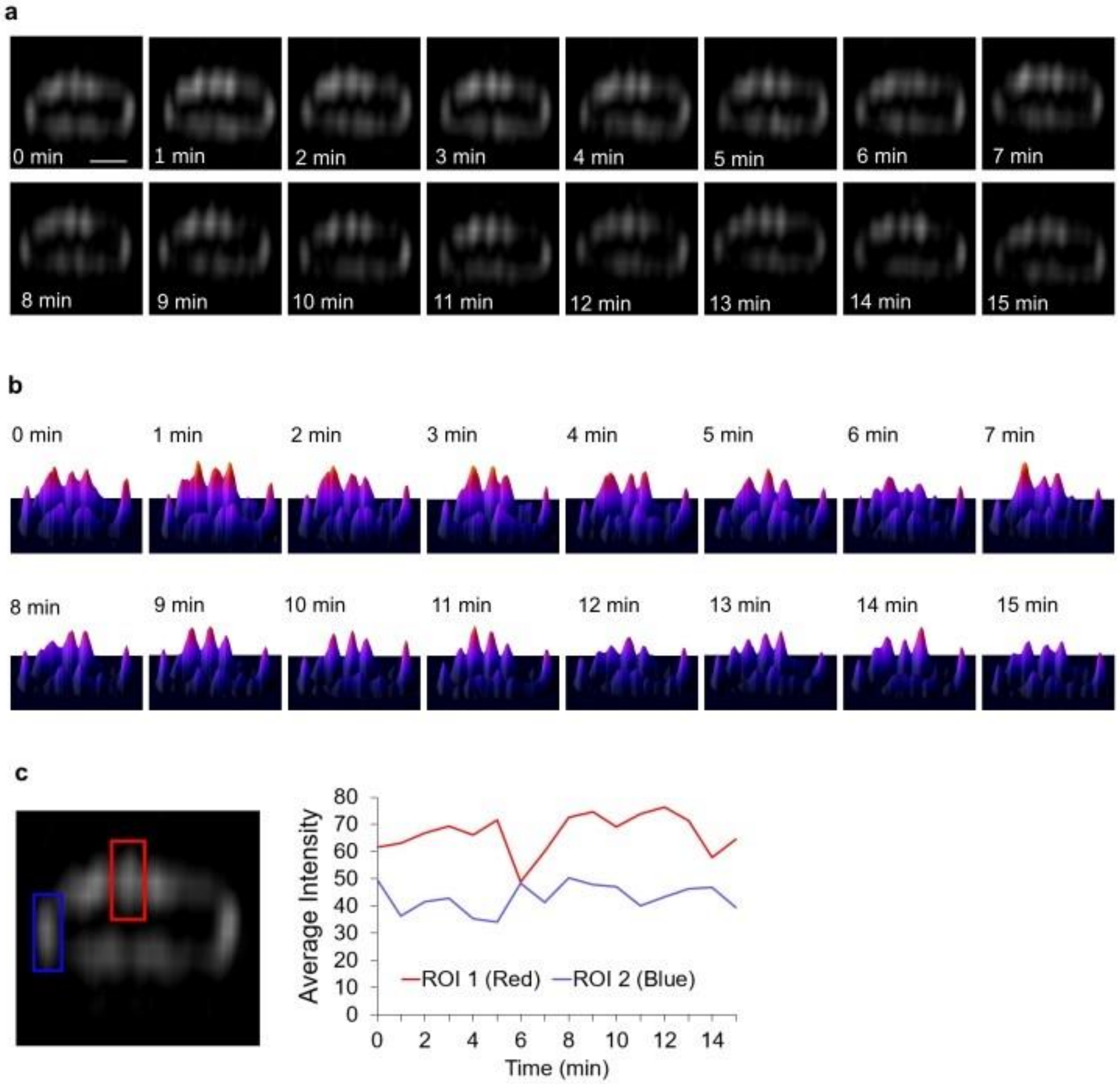


**Supplementary Figure 13**. **3D-SIM time-lapse analysis of CdpA-GFP localization in *H. volcanii.*** The wild-type H26 cells expressing CdpA-GFP were grown in Hv-Cab medium under 2 mM Trp induction to ensure sufficient CdpA-GFP proteins for imaging. The culture was sampled for imaging during mid-log phase. **a,** The distribution and position of CdpA-GFP within the ring did not obviously change from one frame to the next. Scale bar, 0.5 µm. **b,** 3D intensity plots (fire LUT) show that the position of CdpA-GFP molecules within the ring structure may remain constant over time and the intensity of CdpA-GFP fluorescence was not highly dynamic and constantly changing. **c,** Two regions of interest were monitored over time, showing fluctuation of CdpA-GFP fluorescence signal. Source data are provided as a Source Data file.

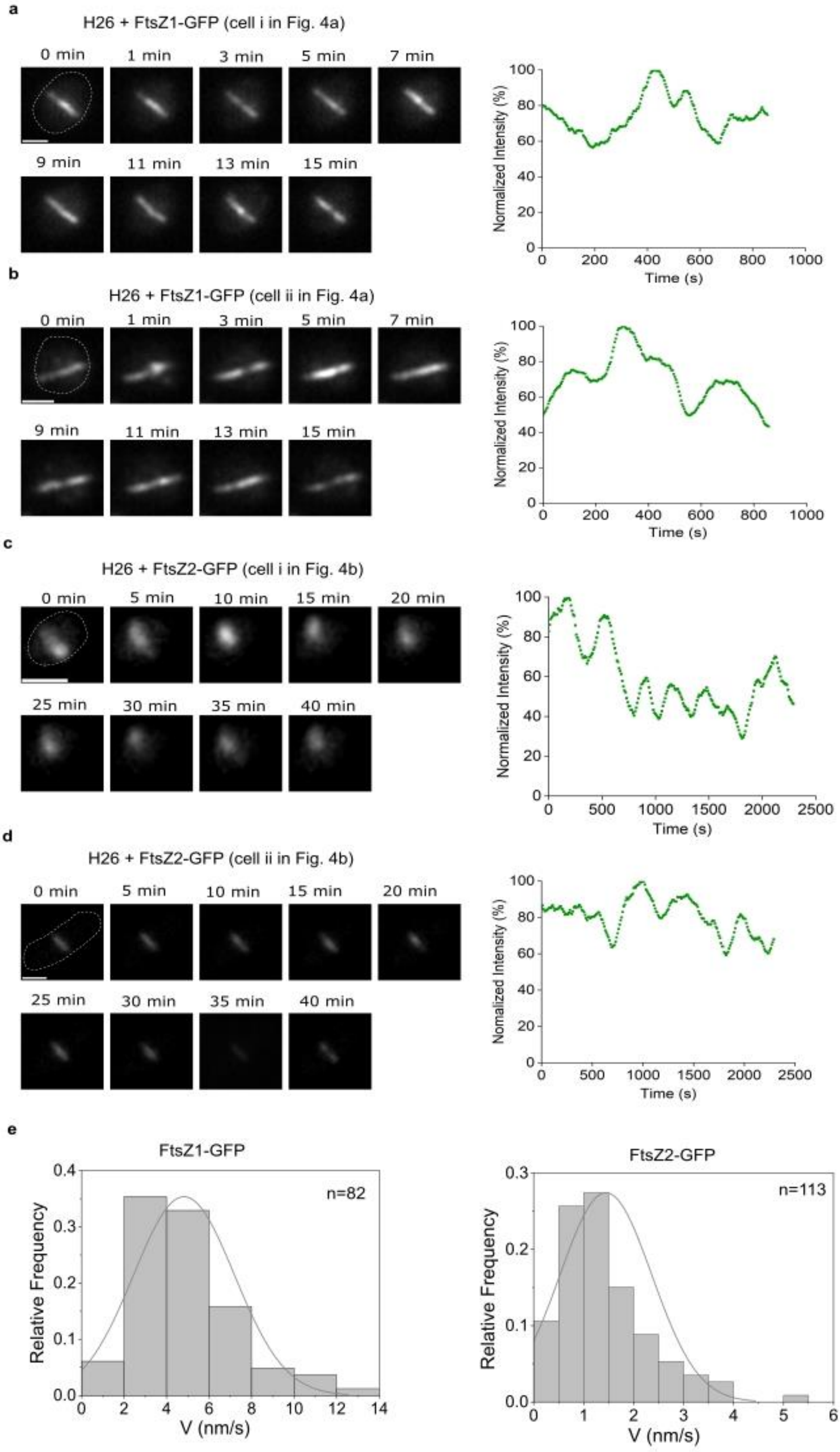
a
H26 + FtsZ1-GFP (cell i in Fig. 4a)
0 min
1 min
3 min
5 min
7 min
9 min
11 min
13 min
15 min
Normalized Intensity (%)
Time (s)
b
H26 + FtsZ1-GFP (cell ii in Fig. 4a)
0 min
1 min
3 min
5 min
7 min
9 min
11 min
13 min
15 min
Normalized Intensity (%)
Time (s)
c
H26 + FtsZ2-GFP (cell i in Fig. 4b)
0 min
5 min
10 min
15 min
20 min
25 min
30 min
35 min
40 min
Normalized Intensity (%)
Time (s)
d
H26 + FtsZ2-GFP (cell ii in Fig. 4b)
0 min
5 min
10 min
15 min
20 min
25 min
30 min
35 min
40 min
Nomalized Intensity (%)
Time (s)
e
FtsZ1-GFP
n=82
Relative Frequency
V (nm/s)
FtsZ2-GFP
n=113
Relative Frequency
V (nm/s)

**Supplementary Figure 14. Examples of analysing FtsZ-ring intensity fluctuations during time-lapse imaging and the quantification of velocity of FtsZ1-GFP an FtsZ2-GFP.** The wild-type H26 cells expressing FtsZ1-GFP or FtsZ2-GFP were grown in Hv-Cab medium without Trp. The mid-log cultures were sampled for TIRF imaging. The fluorescence image of representative cells was corrected for photobleaching, interpolated to 20 nm/pixel and moving averaged over a 12-frame window of time. Source data are provided as a Source Data file. **a-b,** Montages of representative H26+FtsZ1-GFP cells and the integrated fluorescence time trace along the rings. Scale bar, 1 µm. **c-d,** Montages of representative H26+FtsZ2-GFP cells and the integrated fluorescence time trace along the rings. Scale bar, 1 µm. **e,** Distributions of movement speeds of FtsZ1-GFP and FtsZ2-GFP as measured from ring kymographs of individual cells. The estimated average velocity (± SEM) of FtsZ1-GFP was 4.82 ± 0.26 nm/s, and that of FtsZ2-GFP was 1.45 ± 0.09 nm/s. n= number of clusters analyzed from two independent experiments.

**a**

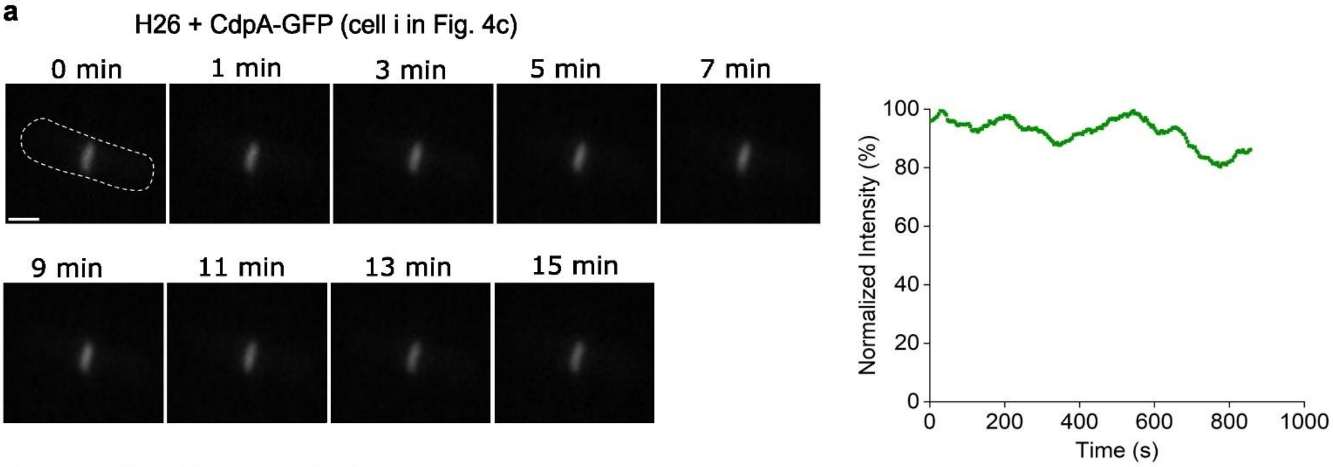
H26 + CdpA-GFP (cell i in Fig. 4c)
0 min
1 min
3 min
5 min
7 min
9 min
11 min
13 min
15 min
Normalized Intensity (%)
Time (s)
0
20
40
60
80
100
200
400
600
800
1000

**b**

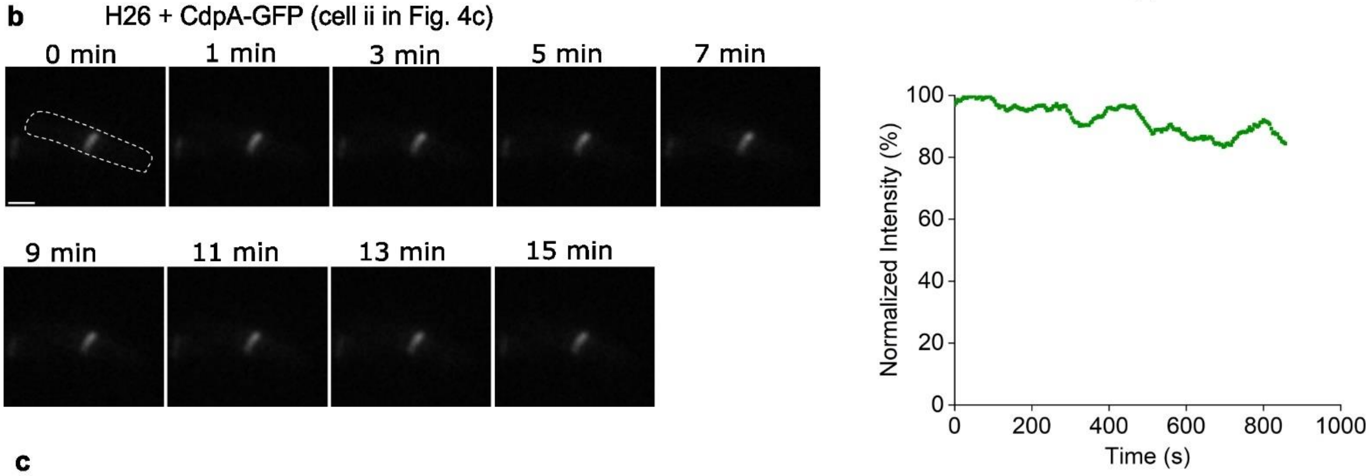
H26 + CdpA-GFP (cell ii in Fig. 4c)
0 min
1 min
3 min
5 min
7 min
9 min
11 min
13 min
15 min
Normalized Intensity (%)
Time (s)
0
20
40
60
80
100
200
400
600
800
1000

**c**

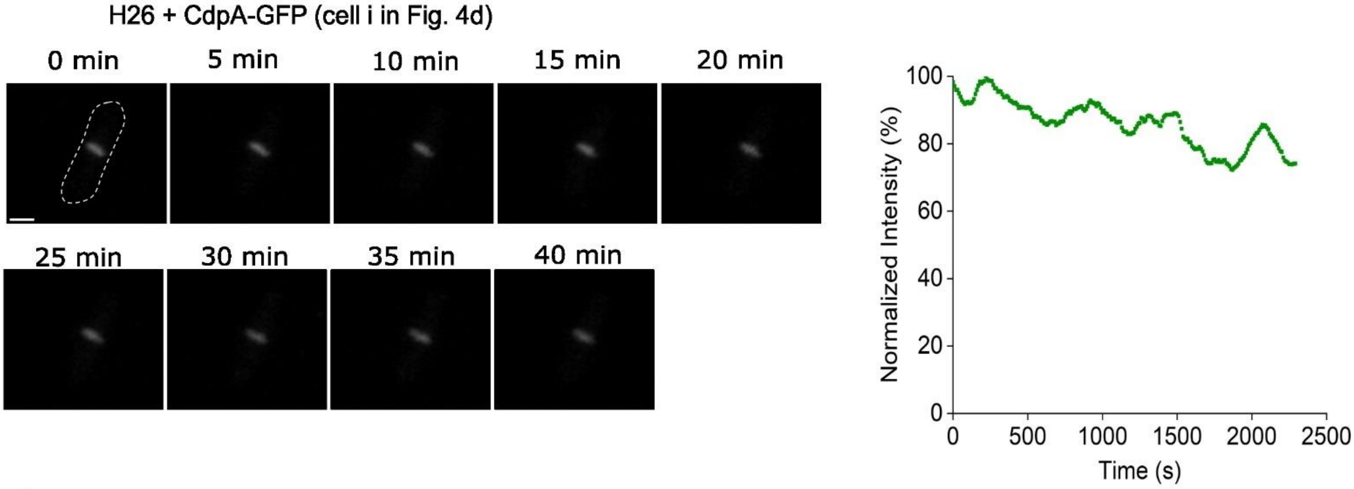
H26 + CdpA-GFP (cell i in Fig. 4d)
0 min
5 min
10 min
15 min
20 min
25 min
30 min
35 min
40 min
Normalized Intensity (%)
Time (s)
0
20
40
60
80
100
500
1000
1500
2000
2500

**d**

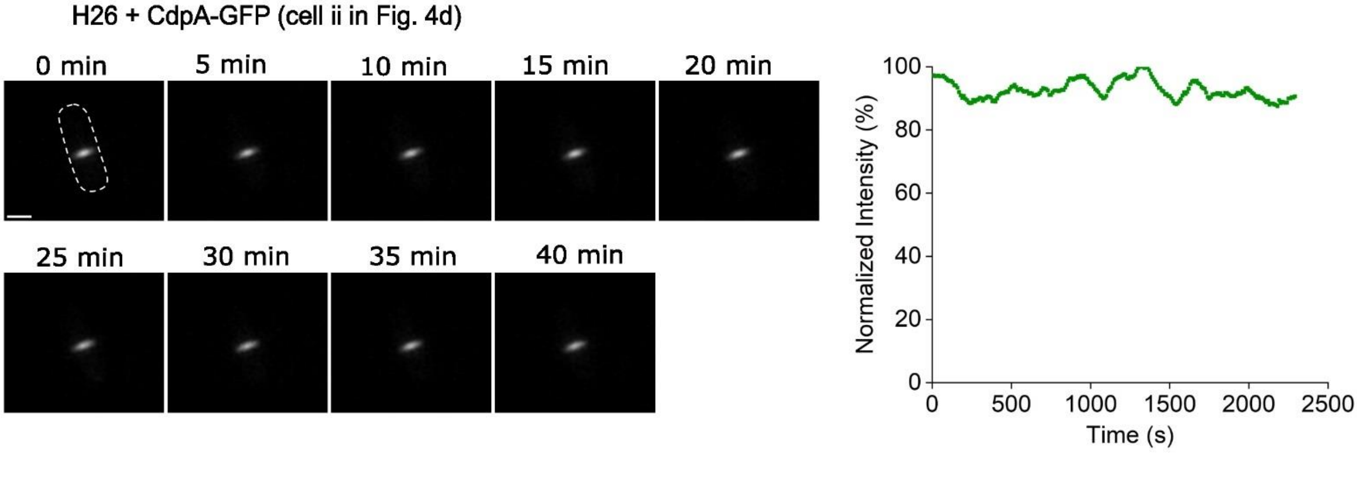
H26 + CdpA-GFP (cell ii in Fig. 4d)
0 min
5 min
10 min
15 min
20 min
25 min
30 min
35 min
40 min
Normalized Intensity (%)
Time (s)
0
20
40
60
80
100
500
1000
1500
2000
2500

**Supplementary Figure 15. Examples of analysing CdpA-ring intensity fluctuations during time-lapse imaging.** The growth medium and imaging pre-processing were same as per Supplementary Fig. 15. **a-b,** Montages of representative H26+CdpA-GFP cells and the integrated fluorescence time trace (15 min with 4s interval) along the rings. **c-d,** Montages of representative H26+CdpA-GFP cells and the integrated fluorescence time trace (40 min with 10s interval) along the rings. All scale bar, 1 µm. Source data are provided as a Source Data file.

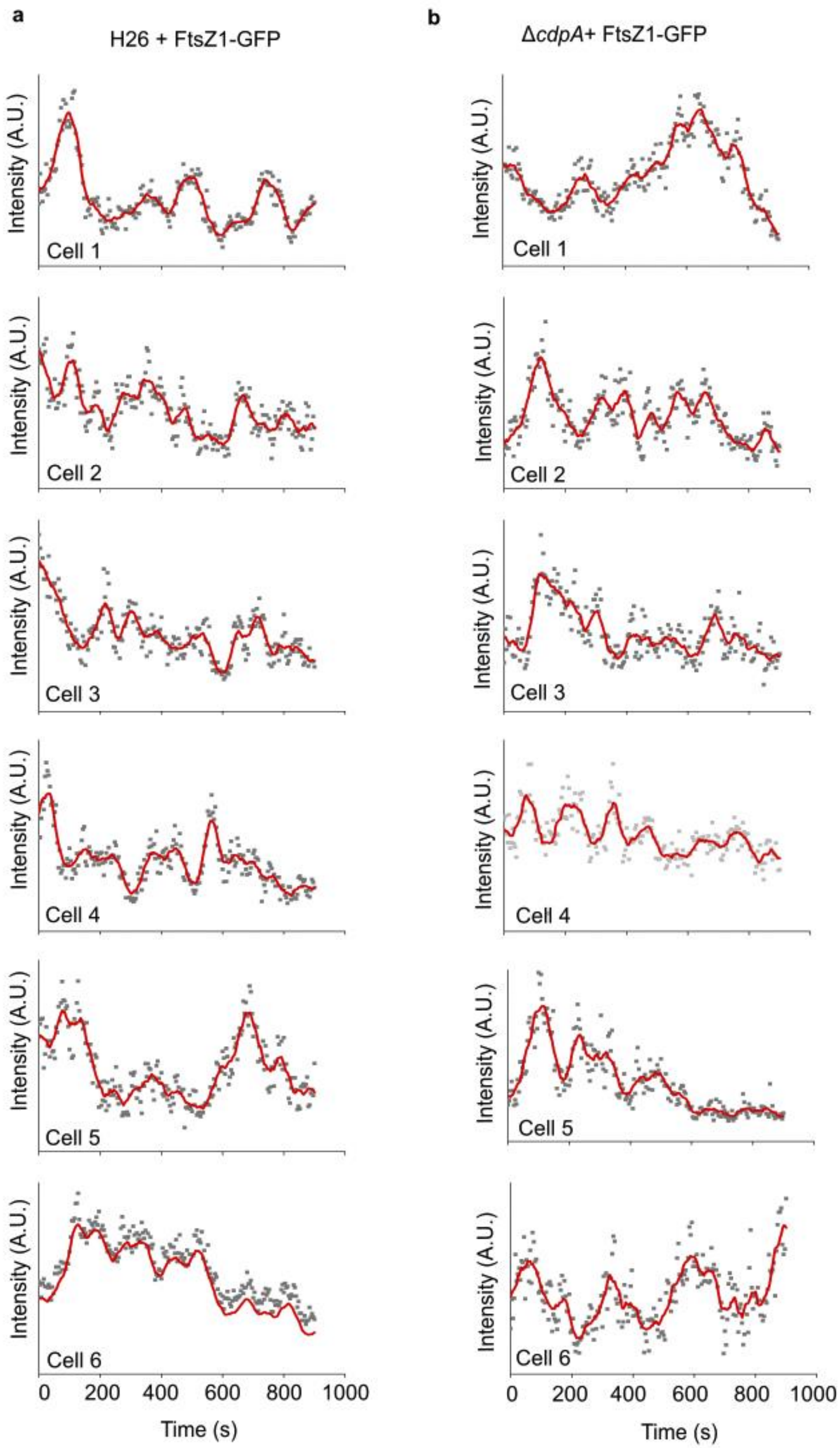


**Supplementary Figure 16. Additional representative fluorescence time traces of FtsZ1-GFP in H26 (a) and Δ*cdpA* (b) in the ROI region of Z-rings.** The intensity was measured in the ROI region within 3×3 pixels in the centre of the rings, as per Fig. 3e-f. Source data are provided as a Source Data file.

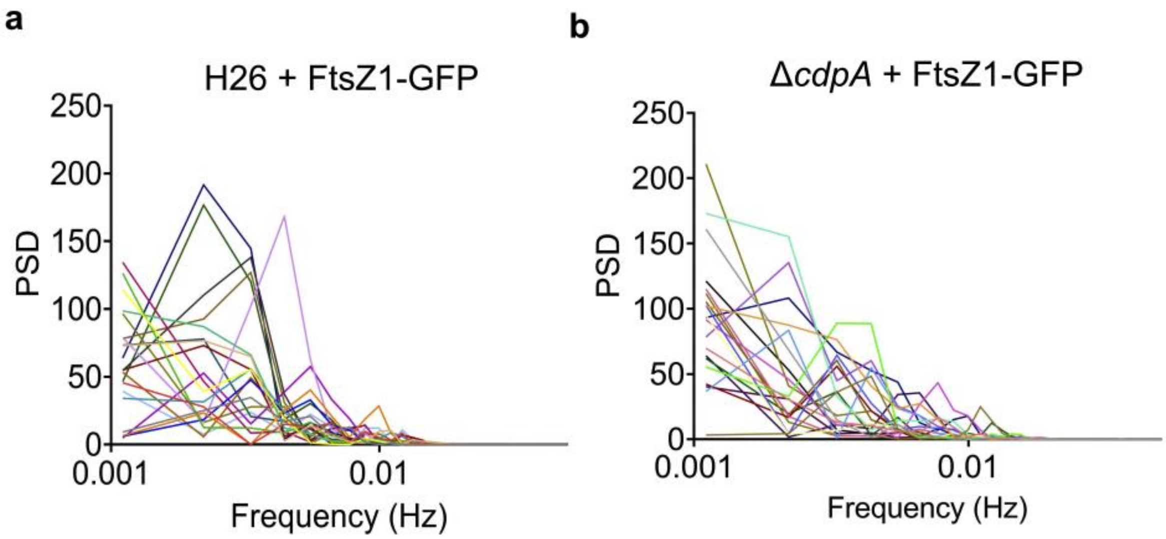


**Supplementary Figure 17. Power spectral density (PSD) curves for individual cells (n=21) of H26 + FtsZ1-GFP (a) and Δ*cdpA* + FtsZ1-GFP (b).** Source data are provided as a Source Data file.

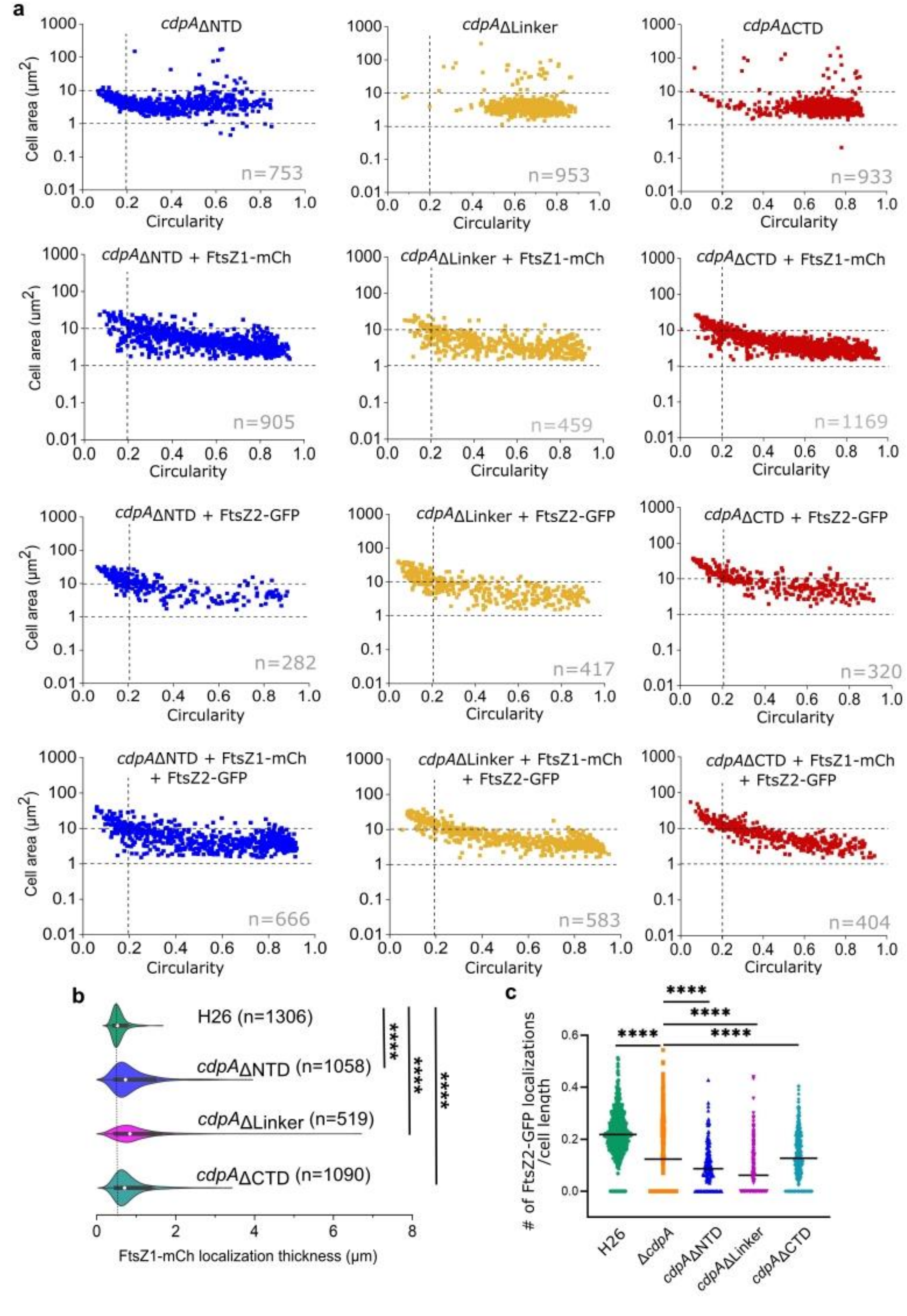

a
cdpAΔNTD
n=753
cdpAΔLinker
n=953
cdpAΔCTD
n=933
cdpAΔNTD + FtsZ1-mCh
n=905
cdpAΔLinker + FtsZ1-mCh
n=459
cdpAΔCTD + FtsZ1-mCh
n=1169
cdpAΔNTD + FtsZ2-GFP
n=282
cdpAΔLinker + FtsZ2-GFP
n=417
cdpAΔCTD + FtsZ2-GFP
n=320
cdpAΔNTD + FtsZ1-mCh + FtsZ2-GFP
n=666
cdpAΔLinker + FtsZ1-mCh + FtsZ2-GFP
n=583
cdpAΔCTD + FtsZ1-mCh + FtsZ2-GFP
n=404
Cell area (μm2)
Circularity
b
H26 (n=1306)
cdpAΔNTD (n=1058)
cdpAΔLinker (n=519)
cdpAΔCTD (n=1090)
****
FtsZ1-mCh localization thickness (μm)
c
### of FtsZ2-GFP localizations /cell length
H26
ΔcdpA
cdpAΔNTD
cdpAΔLinker
cdpAΔCTD
****

**Supplementary Figure 18. Effects of N-terminal domain, linker region and C-terminal domain deletion mutants on cell shape and division. a,** Cell size and shape quantification of the *cdpA*ΔNTD, *cdpA*ΔLinker, and *cdpA*ΔCTD strains during mid-log phase (in Hv-Cab medium + 0.2 mM Trp) with or without plasmids expressing the indicated FtsZ-FP(s). n = number of cells examined over one biologically independent experiment. Representative data are shown from one of two independent experiments. **b,** Quantification of FtsZ1-mCh localization thickness (along the cell's long axis) in wild-type H26, *cdpA*ΔNTD, *cdpA*ΔLinker, and *cdpA*ΔCTD backgrounds. n=number of cells examined over two biologically independent experiments shown. Statistical analysis was performed by Kolmogorov-Smirnov test for non-parametric analysis of difference (**** represents $P < 0.0001$)**. c,** Quantification of number of FtsZ2-GFP localizations per unit cell length, presented as a ratio (y-axis) in wild-type H26 (n=633 cells), Δ*cdpA* (n=446), *cdpA*ΔNTD (n=197), *cdpA*ΔLinker (n=273), and *cdpA*ΔCTD (n=320) backgrounds. Representative data shown from one experiment. Black horizontal lines represent the mean value. Statistical analysis was performed by Kolmogorov-Smirnov test for non-parametric analysis of difference (**** represents $P < 0.0001$). All four *cdpA* mutants have less FtsZ2-GFP localizations than the wild-type, with *cdpA*ΔLinker having lowest mean value. Source data are provided as a Source Data file.

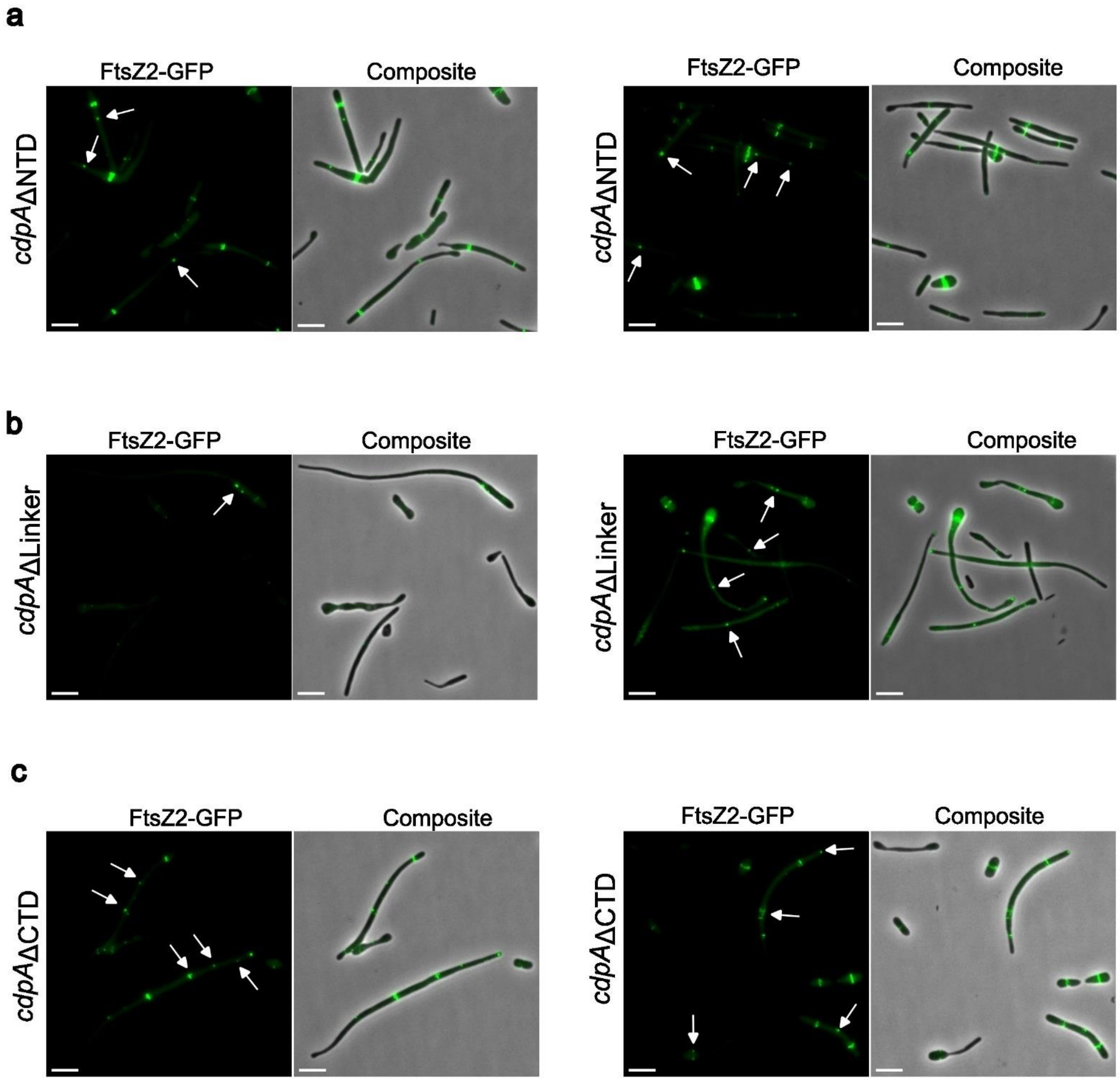


**Supplementary Figure 19. Examples of microscopy images of three CdpA domain deletion strains expressing FtsZ2-GFP. a,** *cdpA*ΔNTD expressing FtsZ2-GFP; **b,** *cdpA*ΔLinker expressing FtsZ2-GFP; **c,** *cdpA*ΔCTD expressing FtsZ2-GFP. The cells were grown in Hv-Cab medium with 0.2 mM Trp and sampled for imaging during mid-log phase. White arrows indicate mispositioned (short perpendicular patches, foci) localizations of FtsZ2-GFP. All scale bars, 5 µm.

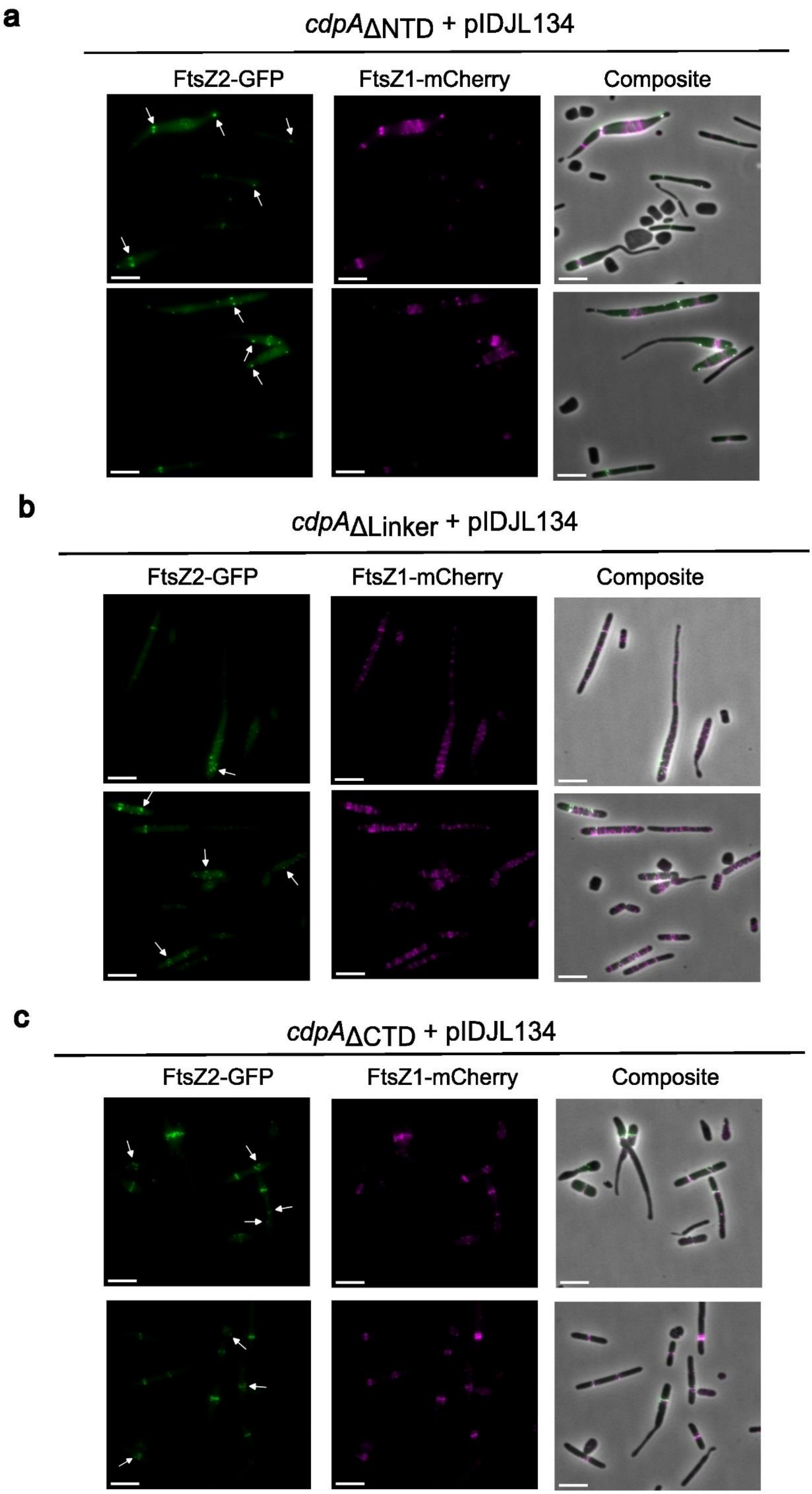

a
cdpAΔNTD + pIDJL134
FtsZ2-GFP
FtsZ1-mCherry
Composite
b
cdpAΔLinker + pIDJL134
FtsZ2-GFP
FtsZ1-mCherry
Composite
c
cdpAΔCTD + pIDJL134
FtsZ2-GFP
FtsZ1-mCherry
Composite

**Supplementary Figure 20. Additional examples of microscopy images of in three CdpA domain deletion strains expressing both FtsZ1-mCherry and FtsZ2-GFP (pIDJL134).** The cells were grown in Hv-Cab medium with 0.2 mM Trp and sampled for imaging during mid-log phase. White arrows indicate the mispositioned or perpendicular localizations of FtsZ2-GFP. All scale bars, 5 µm.

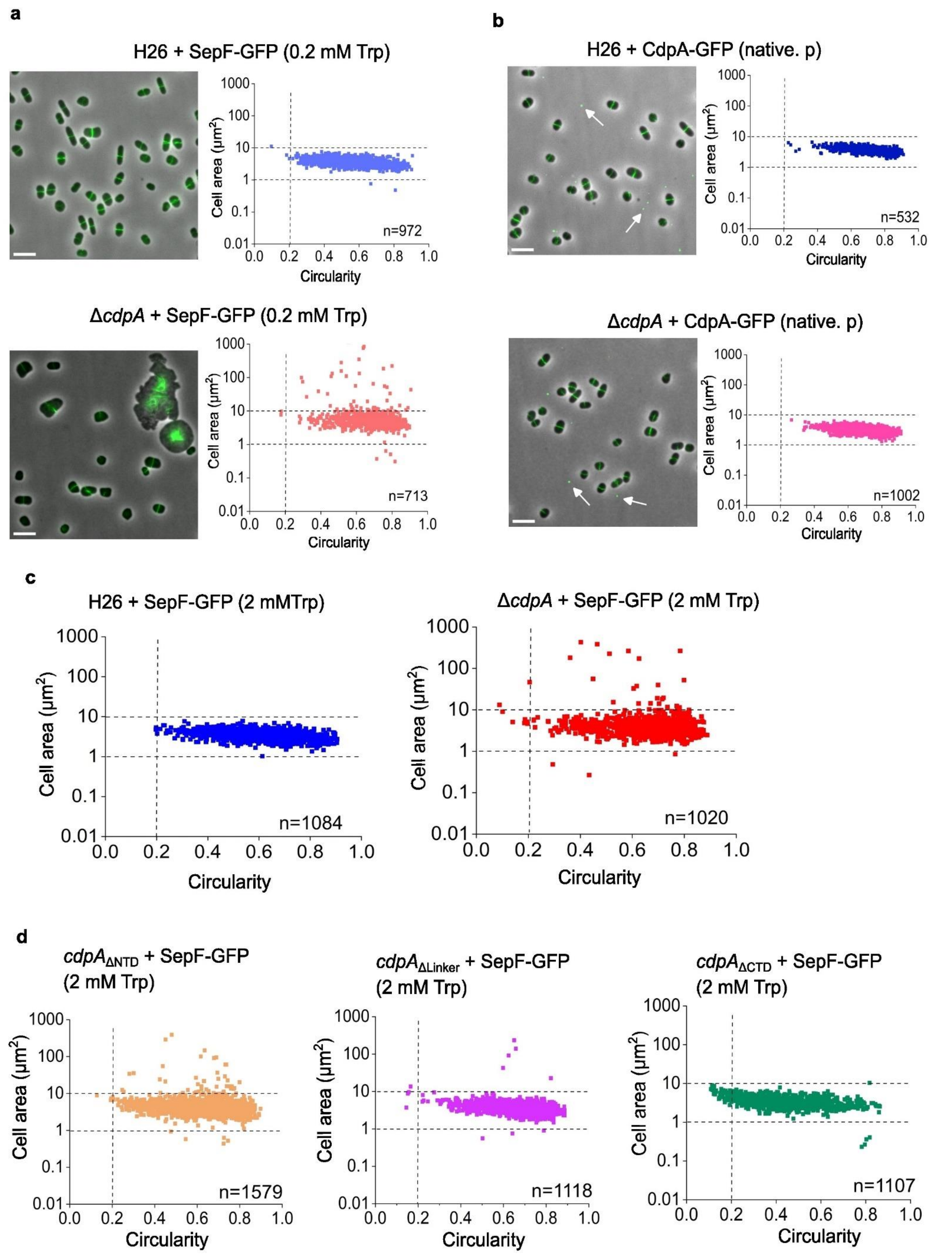


**Supplementary Figure 21. Expression of SepF-GFP in wild-type H26 and *cdpA* mutants. a,** Phase-contrast and fluorescence micrographs and corresponding cell size/shape analysis of the expression of SepF-GFP in H26 and Δ*cdpA* strains from steady mid-log cultures with Hv-Cab medium supplemented 0.2 mM Trp. Scale bars, 5 μm. **b,** Phase-contrast and fluorescence micrographs and corresponding cell size/shape analysis of the expression of CdpA-GFP under *cdpA* native promoter (native. p) in H26 and Δ*cdpA* strains. The expression of CdpA-GFP (native. p) fully corrected the cell division defect of the Δ*cdpA* strain. The strains were grown in Hv-Cab medium and sampled for imaging during mid-log phase. White arrows indicate the extracellular particles. Scale bars, 5 µm. **c,** Cell size/shape analysis of the expression of SepF-GFP in H26 and Δ*cdpA* strains from steady mid-log cultures with Hv-Cab medium supplemented 2 mM Trp. **d,** Cell size/shape analysis of the expression of SepF-GFP in three CdpA domain deletion strains from steady mid-log cultures with Hv-Cab medium supplemented 2 mM Trp. The corresponding micrographs for panel c and d can be found in Fig. 6. n=number of cells examined over one biologically independent experiment. The data shown is representative of at least two independent experiments. Expression of SepF-GFP (either under 0.2 mM or 2 mM Trp) failed to complement the cell division defects in Δ*cdpA* or in the three *cdpA* domain-deletion strains. Source data are provided as a Source Data file.

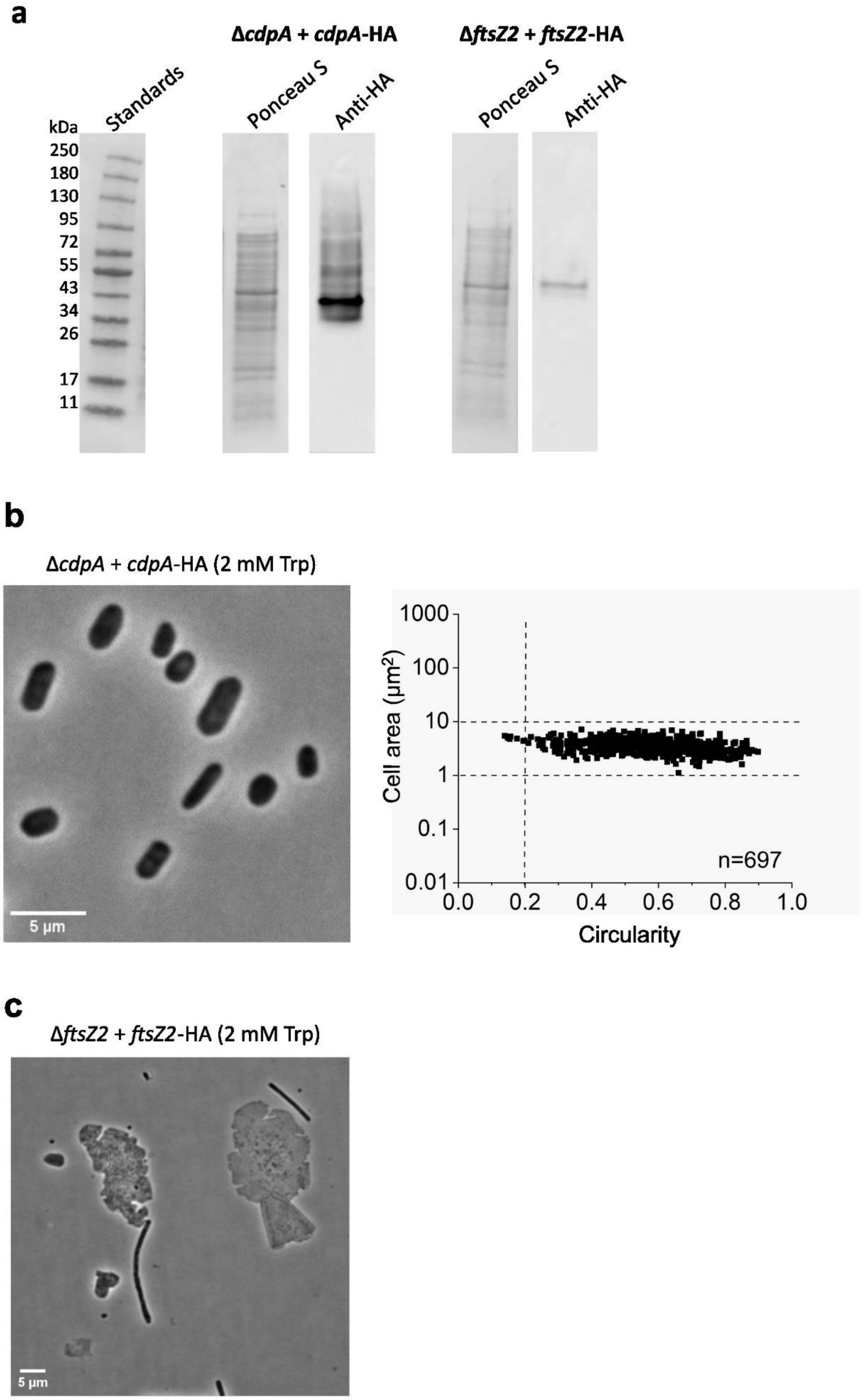


**Supplementary Figure 22. Production of CdpA-HA in *H. volcanii* and complementation of the Δ*cdpA* background by *cdpA*-HA. a,** Western blots for the detection of HA tagged proteins in strains carrying *cdpA*-HA or *ftsZ2*-HA expression plasmids in their corresponding knock-out background strains. Total cell protein (N.B. not crosslinked extract) was prepared for SDS-PAGE and western blotting. Membranes were stained with Ponceau S for total protein visualization and probed with anti-HA antibodies. CdpA-HA consistently appeared with a ladder of bands of higher apparent molecular weight than the expected monomer. FtsZ2-HA shows lower production amounts and no higher MW species detected. All lanes shown are from the same western blot with the same imaging exposure for the Ponceau S and anti-HA probe images, respectively. Unprocessed western blots are provided in the Source Data file. **b,** Phase contrast micrographs of cell producing CdpA-HA in the Δ*cdpA* background grown with 2 mM Trp, showing complementation of the division phenotype of Δ*cdpA* strain by expression of *cdpA*-HA. Scale bar, 5 µm. Cell area and circularity were determined for individual cells. n=number of cells examined over one biologically independent experiment. The data shown is representative of at least two independent experiments. Source data are provided as a Source Data file. **c,** Phase contrast micrographs of cell producing *ftsZ2*-HA in the Δ*ftsZ2* background grown with 2 mM Trp, showing giant and filamentous cells. This suggests that the expression of *ftsZ2*-HA does not complement the cell division defect in the absence of *ftsZ2*.

#### Supplementary Tables

**Supplementary Table 1**. Plasmids and Oligonucleotides used in this study

| Plasmid | Description/function | Oligonucleotides used in construction (5' to 3') | Source |
|---|---|---|---|
| Plasmids for genomic modification of *H. volcanii* | | | |
| pTA131 | Cloning vector with *pyrE2* marker, for *H. volcanii* genetic modification | | [1] |
| pTA131_up_down_0739 | pTA131 with *cdpA*-upstream flank and *cdpA*-downstream flank for *cdpA* gene deletion | 0739_US_F (HindIII):<br>CCGGCCAAGCTTCGATGAGGGCGCGGTAGAGC<br>0739_US_R (EcoRI+BglII):<br>GAATTCGCCGCCCGAAGATCTGGCCCCGACTACGAAACCAGC<br>0739_DS_F (BglII + EcoRI):<br>AGATCTTCGGGCGGCGAATTCTCTGTCGACGACTCTGACGGC<br>0739_DS_R (BamHI):<br>CCGGCCGGATCCATCTGCGGATGTGCGGCTAC | This study |
| pTA131_CdpA_delNTD | pTA131 with upstream flank and downstream flank of CdpA N-terminal domain (NTD) for its deletion (17-149 aa) | Synthetic gene (Genscript) | This study |
| pTA131_CdpA_delLinker | pTA131 with upstream flank and downstream flank of CdpA linker region for its deletion (150-284 aa) | Synthetic gene (Genscript) | This study |
| pTA131_CdpA_delCTD | pTA131 with upstream flank and downstream flank of CdpA C-terminal domain for its deletion (285-329 aa) | CdpA_CTS_UP_F (HindIII):<br>CGACGGTATCGATAAGCTTGACCTCACGCTCGAAACC<br>CdpA_CTS_ UP_R _Overlap:<br>AGAGTCGTCGACAGACTAGCCGACGGTGGCGTTCGC<br>CdpA_CTS_DS_Forward: TAGTCTGTCGACGACTCTGACG<br>CdpA_CTS_DS_Reverse (BamHI):<br>CTCTAGAACTAGTGGATCCTATCTGCGGATGTGCGGC | This study |
| Plasmids for gene expression in *H. volcanii* | | | |
| pTA962 | *p.tna* expression vector for *H. volcanii* | | [2] |
| pIDJL40 | pTA962 with gfp (BamHI-NotI) | | [3] |
| pTA962_*cdpA* (HVO_0739) | *p.tna* control of *cdpA* | HVO_0739_f:<br>CCCCCGGGAATTCATATGACCAGTCTTTCGGAGGCG<br>HVO_0739_r:<br>CGCGGATCCCTACTCGCGGCGGCGCGGGAA | This study |
| pIDJL40_*cdpA* (HVO_0739) | *p.tna* control of *cdpA*-GFP | HVO_0739_f:<br>CCCCCGGGAATTCATATGACCAGTCTTTCGGAGGCG<br>HVO_0739 (NS)_r:<br>CGCGGATCCCTCGCGGCGGCGCGGGAA | This study |
| pIDJL40- *ftsZ1* | *p.tna* control of *ftsZ1-GFP* | | [4] |
| pTA962-*ftsZ1*-mCherry | *p.tna* control of *ftsZ1-mCherry* | | [4] |
| pIDJL40- *ftsZ2* | *p.tna* control of *ftsZ2-GFP* | | [4] |
| pTA962-*ftsZ2*-mCherry | *p.tna* control of *ftsZ2-mCherry* | | [4] |
| pIDJL134 | *p.tna* control of *ftsZ2*-GFP and *ftsZ1*-mCherry | | [4] |
| pIDJL40-*cdpA*+*ftsZ1*-mCherry | *p.tna* control of *cdpA*-GFP and *ftsZ1-mCherry* | | This study |

| | | | |
|---|---|---|---|
| pIDJL40-*cdpA+ftsZ2*-mCherry | *p.tna* control of *cdpA*-GFP and *ftsZ2-mCherry* | | This study |
| pIDJL40-*cdpA* (native. p) | *cdpA* native promoter control of *cdpA*-GFP | CdpA_nativeP_F: ATACGCGGGCCCGCGCGAGGGTTAGCCAC HVO_0739 (NS)_r: CGCGGATCCCTCGCGGCGGCGCGGGAA | This study |
| pIDJL40_CdpA_NTD | *p.tna* control of N-terminal domain of CdpA (1-149 aa) fused to GFP | HVO_0739_f: CCCCCGGGAATTCATATGACCAGTCTTTCGGAGGCG CdpA_NTD_R: CGCGGATCCGAAGTTGGCGACGCCGAC | This study |
| pIDJL40_CdpAΔNTD | *p.tna* control of CdpA without N-terminal domain fused to GFP | HVO_0739_f: CCCCCGGGAATTCATATGACCAGTCTTTCGGAGGCG HVO_0739 (NS)_r: CGCGGATCCCTCGCGGCGGCGCGGGAA | This study |
| pIDJL40_CdpAΔCTD | *p.tna* control of CdpA without C-terminal domain (1-284 aa) fused to GFP | HVO_0739_f: CCCCCGGGAATTCATATGACCAGTCTTTCGGAGGCG CdpA_delCTD_R: CGCGGATCCGCCGACGGTGGCGTTCGC | This study |
| pTA962-HA | *p.tna* control of HA tag | | This study |
| pTA962-CdpA-HA | *p.tna* control of CdpA with HA tag | | This study |
| pTA962-FtsZ2-HA | *p.tna* control of FtsZ2 with HA tag | | This study |
| pSVA3942 | *p.tna* control of SepF-GFP | | [5] |

**Supplementary Table 2**. Strains used in this study

| Strain | Genotype | Description | Source |
|---|---|---|---|
| | | *E. coli* | |
| DH5α | *fhuA2 Δ(argF-lacZ)U169 phoA glnV44 Φ80 Δ(lacZ)M15 gyrA96 recA1 relA1 endA1 thi-1 hsdR17* | General cloning strain for plasmid construction | Invitrogen |
| C2925 | *ara-14 leuB6 fhuA31 lacY1 tsx78 glnV44 galK2 galT22 mcrA dcm-6 hisG4 rfbD1 R(zgb210::Tn10)* Tet$^{S}$ *endA1 rspL136* (Str$^{R}$) *dam13::Tn9* (Cam$^{R}$) *xylA-5 mtl-1 thi-1 mcrB1 hsdR2* | DNA methylation-deficient strain for preparation of demethylated plasmids for *H. volcanii* transformation | New England Biolabs |
| | | *H. volcanii* | |
| H26 (ID3) | (DS70) Δ*pyrE2* | Auxotroph (uracil) | T. Allers |
| H98 (ID6) | (DS70) Δ*pyrE2* Δ*hdrB* | Auxotroph (uracil, hypoxanthine and thymidine) | T. Allers |
| ID41 | (H98) pTA962 | H98 carrying pTA962 | [4] |
| ID550 | (H98) pTA962- *cdpA* | Carries plasmid for Trp-regulated expression of *cdpA* (HVO_0739) | This study |
| YL29 | (H26) pIDJL40_*cdpA* | Carries plasmid for expression of *cdpA-gfp* | This study |
| YL6 | (H26) pIDJL40-*cdpA+ftsZ1*-mCherry | Carries plasmid for dual expression of *cdpA-gfp* and *ftsZ1-mCherry* | This study |
| YL7 | (H26) pIDJL40-*cdpA+ftsZ2*-mCherry | Carries plasmid for dual expression of *cdpA-gfp* and *ftsZ2-mCherry* | This study |
| YL24 | (H26) pIDJL40- *ftsZ1* | Carries plasmid for expression of *ftsZ1-gfp* | This study |
| YL26 | (H26) pIDJL40- *ftsZ2* | Carries plasmid for expression of *ftsZ2-gfp* | This study |
| YL25 | (H26) pTA962-*ftsZ1*-mCherry | Carries plasmid for expression of *ftsZ1-mCherry* | This study |
| YL72 | (H26) pIDJL40_CdpA_NTD | Carries plasmid for expression of N-terminal domain of CdpA fused to GFP | This study |
| YL97 | (H26) pIDJL40_CdpAΔNTD | Carries plasmid for expression of modified CdpA (without N-terminal domain) fused to GFP | This study |

| | | | |
|---|---|---|---|
| YL73 | (H26) pIDJL40_CdpAΔCTD | Carries plasmid for expression of modified CdpA (without C-terminal domain) fused to GFP | This study |
| YL28 | (H26) pIDJL134 | Carries plasmid for dual expression of *ftsZ2-gfp* and *ftsZ1-mCherry* | This study |
| ID750 | (H26) pTA962-HA | Carries plasmid for expression of HA | This study |
| ID562 | (H26) Δ*cdpA* | Deletion of *cdpA* | This study |
| ID568 | (Δ*cdpA*) pTA962_*cdpA* | Δ*cdpA* and plasmid for expression *cdpA* | This study |
| ID569 | (Δ*cdpA*) pIDJL40_*cdpA* | Δ*cdpA* and plasmid for expression *cdpA-gfp* | This study |
| ID570 | (Δ*cdpA*) pIDJL40 | Δ*cdpA* and plasmid for expression *gfp* | This study |
| ID563 | (Δ*cdpA*) pIDJL40- *ftsZ1* | Δ*cdpA* and plasmid for expression *ftsZ1-gfp* | This study |
| ID564 | (Δ*cdpA*) pIDJL40- *ftsZ2* | Δ*cdpA* and plasmid for expression *ftsZ2-gfp* | This study |
| ID565 | (Δ*cdpA*) pTA962-*ftsZ1*-mCherry | Δ*cdpA* and plasmid for expression *ftsZ1-mCherry* | This study |
| ID567 | (Δ*cdpA*) pIDJL134 | Δ*cdpA* and plasmid for dual expression of *ftsZ2-gfp* and *ftsZ1-mCherry* | This study |
| ID756 | (Δ*cdpA*) pTA962-CdpA-HA | Carries plasmid for expression of CdpA-HA | This study |
| ID76 | Δ*ftsZ1* | Deletion of *ftsZ1* | [4] |
| ID77 | Δ*ftsZ2* | Deletion of *ftsZ2* | [4] |
| ID112 | Δ*ftsZ1* Δ*ftsZ2* | Double deletion of *ftsZ1* and *ftsZ2* | [4] |
| ID555 | (Δ*ftsZ1*) pIDJL40_*cdpA* | Δ*ftsZ1* and plasmid for expression *cdpA-gfp* | This study |
| ID556 | (Δ*ftsZ2*) pIDJL40_*cdpA* | Δ*ftsZ2* and plasmid for expression *cdpA-gfp* | This study |
| ID754 | (Δ*ftsZ2*) pTA962-FtsZ2 -HA | Carries plasmid for expression of FtsZ2-HA | This study |
| ID112 | (Δ*ftsZ1* Δ*ftsZ2*) pIDJL40_cdpA | Δ*ftsZ1* Δ*ftsZ2* and plasmid for expression *cdpA-gfp* | This study |
| YL84 | *cdpA*ΔNTD | Deletion of N-terminal domain (17-149 aa) of CdpA | This study |
| YL85-1 | (*cdpA*ΔNTD) pTA962-*ftsZ1*-mCherry | *cdpA*ΔNTD and plasmid for expression *ftsZ1-mCherry* | This study |
| YL87 | (*cdpA*ΔNTD) pIDJL40- *ftsZ2* | *cdpA*ΔNTD and plasmid for expression *ftsZ2-gfp* | This study |
| YL89 | (*cdpA*ΔNTD) pIDJL134 | *cdpA*ΔNTD and plasmid for dual expression of *ftsZ2-gfp* and *ftsZ1-mCherry* | This study |
| YL85 | *cdpA*ΔLinker | Deletion of linker region (150-284 aa) of CdpA | This study |
| YL91 | (*cdpA*ΔLinker) pTA962-*ftsZ1*-mCherry | *cdpA*ΔLinker and plasmid for expression *ftsZ1-mCherry* | This study |
| YL93 | (*cdpA*ΔLinker) pIDJL40- *ftsZ2* | *cdpA*ΔLinker and plasmid for expression *ftsZ2-gfp* | This study |
| YL95 | (*cdpA*ΔLinker) pIDJL134 | *cdpA*ΔLinker and plasmid for dual expression of *ftsZ2-gfp* and *ftsZ1-mCherry* | This study |
| YL14 | *cdpA*ΔCTD | Deletion of C-terminal domain (285-329 aa) of CdpA | This study |
| YL18 | (*cdpA*ΔCTD) pTA962-*ftsZ1*-mCherry | *cdpA*ΔCTD and plasmid for expression *ftsZ1-mCherry* | This study |
| YL19 | (*cdpA*ΔCTD) pIDJL40- *ftsZ2* | *cdpA*ΔCTD and plasmid for expression *ftsZ2-gfp* | This study |
| YL21 | (*cdpA*ΔCTD) pIDJL134 | *cdpA*ΔCTD and plasmid for dual expression of *ftsZ2-gfp* and *ftsZ1-mCherry* | This study |
| HTQ239 | | SepF depleted strain | [5] |
| YL79 | (HTQ239) pIDJL40-*cdpA* (native. p) | SepF depleted strain and plasmid for expression of CdpA-GFP under *cdpA* native promoter | This study |
| YL71 | (H26) pIDJL40-*cdpA* (native. p) | H26 and plasmid for expression of CdpA-GFP under *cdpA* native promoter | This study |
| YL76 | (Δ*cdpA*) pIDJL40-*cdpA* (native. p) | Δ*cdpA* and plasmid for expression of CdpA-GFP under *cdpA* native promoter | This study |
| YL23 | (H26) pSVA3942 | H26 and plasmid for expression of CdpA-GFP under *cdpA* native promoter | This stsudy |
| YL98 | (*cdpA*ΔNTD) pSVA3942 | *cdpA*ΔNTD and plasmid for expression of SepF-GFP | This study |
| YL99 | (*cdpA*ΔLinker) pSVA3942 | *cdpA*ΔLinker and plasmid for expression of SepF-GFP | This study |
| YL2 | (Δ*cdpA*) pSVA3942 | Δ*cdpA* and plasmid for expression of SepF-GFP | This study |
| YL16 | (*cdpA*ΔCTD) pSVA3942 | *cdpA*ΔCTD and plasmid for expression of SepF-GFP | This study |
| YL182 | (Δ*ftsZ1*) pIDJL40-*cdpA*+*ftsZ2*-mCherry | Δ*ftsZ1* and plasmid for dual expression of *cdpA-gfp* and *ftsZ2-mCherry* | This study |
| YL183 | (Δ*ftsZ2*) pIDJL40-*cdpA*+*ftsZ1*-mCherry | Δ*ftsZ2* and plasmid for dual expression of *cdpA-gfp* and *ftsZ1-mCherry* | This study |
